# GPerturb: Gaussian process modelling of single-cell perturbation data

**DOI:** 10.1101/2025.03.26.645455

**Authors:** Hanwen Xing, Christopher Yau

**Affiliations:** Nuffield Department for Women’s and Reproductive Health, University of Oxford, Oxford, UK; Health Data Research UK, London, UK

## Abstract

Single-cell RNA sequencing and Clustered Regularly Interspaced Short Palindromic Repeats (CRISPR) screening facilitate the high-throughput study of genetic perturbations at a single-cell level. Characterising combinatorial perturbation effects, such as the subset of genes affected by a specific perturbation, is crucial yet computationally challenging in the analysis of single-cell CRISPR screening datasets due to the sparse and complex structure of unknown biological mechanisms. We propose Gaussian process based sparse perturbation regression (GPerturb) to identify and estimate interpretable gene-level perturbation effects for such data. GPerturb uses an additive structure to disentangle perturbation-induced variation from background noise, and can learn sparse, gene-level perturbation-specific effects from either discrete or continuous responses of perturbed samples. Crucially, GPerturb provides uncertainty estimates for both the presence and magnitude of perturbation effects on individual genes. We validate the efficacy of GPerturb on both simulated and real-world datasets, demonstrating that its prediction and generalisation performance is competitive with existing state-of-the-art methods. Using real-world datasets, we also show that the model reveals interesting gene-perturbation interactions and identifies perturbation effects consistent with known biological mechanisms. Our findings confirm the utility of GPerturb in revealing new insights into the complex dependency structure between gene expressions and perturbations.

## Introduction

Developments in Single-cell RNA sequencing (scRNA-seq) and Clustered Regularly Interspaced Short Palindromic Repeats (CRISPR) screening accelerate the discovery of association between genes and various biological processes such as immune responses, cell proliferation and drug resistance (Dixit et al., 2016; Srivatsan et al., 2020; Buquicchio and Satpathy, 2021; Shalem et al., 2014; Rosati and Giordano, 2022; Bock et al., 2022; **?**). In particular, technologies such as CROP sequencing (CROP-seq) (Datlinger et al., 2017) and Perturb sequencing(Dixit et al., 2016) (Perturb-seq) have made high-throughput, large scale cellular perturbation screens possible. Such cellular perturbation screens allow practitioners to investigate complex biological mechanisms such as regulatory dependencies and drug responses on a single-cell level using the comprehensive, fine-grained readouts of the target perturbations within single cells, and have found applications in studies such as combinatorial therapy (Al-Lazikani et al., 2012; Aissa et al., 2021), drug discovery (Van de Sande et al., 2023) and regulatory elements (Jin et al., 2020; Yao et al., 2023).

The growing granularity of measurements provided by single-cell CRISPR screening technologies motivates the need for novel computational methods to help extracting interpretable biological insights from generated data particularly in relation to the perturbation effects. However, it is a challenging task due to the high dimensionality, complex structure and sparse nature of the single-cell screening measurements. Analytically, the problem is to produce a prediction model that can be used to provide an estimate of the effect of a perturbation on expression for any particular cell type or cell. The model can be developed using a training dataset that consists of a series of expression measurements on single cells, in which each cell belongs to one of a finite number of cell types and has been subject to one of a finite number of perturbations (including unperturbed controls).

A common approach has been to apply deep learning-based techniques to learn the relationships between cell type, perturbation and expression output flexibly from sufficiently large datasets. To do this, expression data and cell type information are typically transformed into *embeddings* via deep neural networks (DNNs), which are learnt low-dimensional projections of the original measurements, and then the effect of the perturbations are described in this embedding space. Perturbed embeddings can then be remapped back to expression measurements. If there are a large number of perturbations, embeddings may also be formed for the perturbations themselves. Models are trained against an objective which seeks to minimise the discrepancy between the observed perturbation effect and that predicted via the model. A bottlenecking effect in the design of the model architectures, which perform the various embedding and output transformations, leads to the creation of compressed representations that are optimised towards the maximal retention of information.

The Compositional Perturbation Autoencoder (Lotfollahi et al., 2023) (CPA) is an example of such an approach. Given the measured unperturbed and perturbed expression of a cell, CPA predicts the counterfactual distribution of the expression of that cell had it been subjected to a different perturbation. CPA adopts an autoencoder learning framework and uses additive latent embedding of the cell and perturbation states. For instance, SAMS-VAE (Bereket and Karaletsos, 2023) using a sparse additive mechanism shift variational autoencoder to characterise perturbation effects as sparse latent representations. In SAMS-VAE, the latent representation of a perturbed expression vector is obtained by adding a sparse representation of the perturbation to a dense perturbation-independent basal state, and the decoder is trained to reconstruct the perturbed expression vectors from latent representations. Other other approaches have sought to embed additional external information about the expression features to improve predictions. GEARS (Roohani et al., 2023) uses a knowledge graph of gene-gene relationships to inform the prediction allowing it to simulate the outcomes of perturbing unseen single genes or combinations of genes. Although not deep learning-based, CellOT (Bunne et al., 2023) leverages DNNs for function estimation in a neural optimal transport framework (Korotin et al., 2022) to map between unperturbed and perturbed single-cell responses.

More recently, single cell foundation models (Theodoris et al., 2023; Hao et al., 2024; Cui et al., 2024; Chen and Zou, 2024) have emerged which promise to provide a multi-functional basis for many analytical applications (Ma et al., 2024). However, the benefits of current foundation model approaches are not yet clear (Kedzierska et al., 2023; Wenteler et al., 2024; Li et al., 2024a; Ahlmann-Eltze et al., 2024) and fair evaluation is complicated by emerging applications that integrate direct empirical data with knowledge extracted from scientific literature or pre-trained foundation models (Istrate et al., 2024; Li et al., 2024a).

Non-deep learning approaches have also been developed. Guided Sparse Factor Analysis (Zhou et al., 2023) (GSFA) models continuous observations and adopts a linearity assumption in its multivariate latent factor regression approach. While a variant of the “factorize-recover” algorithm was used to infer perturbation effects from composite sample phenotypes from compressed Perturb-seq experimental data using combination of sparse principal components analysis and LASSO regression (Yao et al., 2023). The attractiveness of such approaches is their comparative simplicity due to the use of linearity assumptions.

The plethora of computational perturbation modelling methods (Gavriilidis et al., 2024; Rood et al., 2024) disguises many practical issues that are only apparent at usage time. For instance, assumptions in experimental setup and data preprocessing can be implicitly built into models. CPA assumes categorical cell-level information and continuous gene expression inputs but SAMS-VAE is not able to incorporate additional cell-level information, such as batch information or cell type, and can only handle binary perturbation and count-based expression inputs. GSFA utilises its own particular approach to input data transformation and normalisation approaches. While CPA is able to process continuous perturbation levels (e.g. dosage), GEARS applies only to discrete perturbations as it uses perturbation embeddings and relational graphs between perturbations. Foundation models often require their own specific approaches for to-kenisation and input data embedding. These model design differences, on a practical level, mean direct and intuitive comparisons between methods may not be possible both in terms of their predictions but also in terms of the explanations underlying those.

In this work, we propose a more conceptually classical approach for perturbation modelling, called **GPerturb**, which utilises hierarchical Bayesian modelling (van de Schoot et al., 2021) and Gaussian Process regression (Williams and Rasmussen, 1995). We demonstrate that **GPerturb** can achieve high levels of predictive performance that is comparable to current state of the art perturbation models even using a sparse, additive modelling structure and without the use of latent embeddings or external information. Further, the modularity of the hierarchical construction allows us to examine the effect of swapping between an observational data model based on count-based expression data for one which uses continuous transformed values instead. Despite the abundance of perturbation modelling methods available, **GPerturb** offers a novel and scalable generative modelling approach with classical features which make prediction output and their interpretation more readily understandable compared to methods based on black box learning.

## Results

### Overview

We first provide an overview of **GPerturb** (a more detailed mathematical description is provided in **Methods**) which is a generative model that aims to directly identify and estimate sparse, interpretable gene-level perturbation effects, for analysing single-cell CRISPR screening data. In **GPerturb**, we assume that the each expression feature measured for each cell can be explained as a sample from a distribution. In the case where expression data is continuously value, a normal distribution is used (zero-inflated Poisson for count-based data) where the mean expression level is given by the combination of two components. The first is a feature-specific basal expression level which is determined by the cell-specific parameters (e.g. cell type or cell-specific sequencing information). The second component is a feature-specific perturbation effect which depends on the type of perturbation applied to the cell (which can be null). To make it explicit that some perturbations will only affect certain features, the perturbation component for each feature is controlled by a binary on/off switch. The relationships that map cell-specific parameters and perturbation type to the observed expression level are governed by nonlinear Gaussian processes.

**GPerturb** adopts a supervised learning approach to learn and disentangle the basal (unperturbed) expression distribution associated with a given cell type and the additive effect of perturbations given observed gene expression measurements (Figure 1a). Gaussian processes (Williams and Rasmussen, 1995) are used to model expression functions, and sparsity constraints aim to regularise the model and improve generalisation and robustness. The generative properties of **GPerturb** allow expression levels to be predicted (Figure 1b), the associative analysis of perturbation effects with biological pathways (Figure 1c), and facilitates users in identifying details about complex perturbation-gene dependencies (Figure 1d). Compared with existing methods, **GPerturb** does not require a latent variable construction and incorporates uncertainty propagation in an intuitive way due to the Bayesian framework. It can be applied to either raw count (**GPerturb**-ZIP, for zero-inflated Poisson) or continuous transformed expression measurements (**GPerturb**-Gaussian). Further and detailed information about the model development and relationships to existing methods can be found in **Methods**.

**Figure 1:**
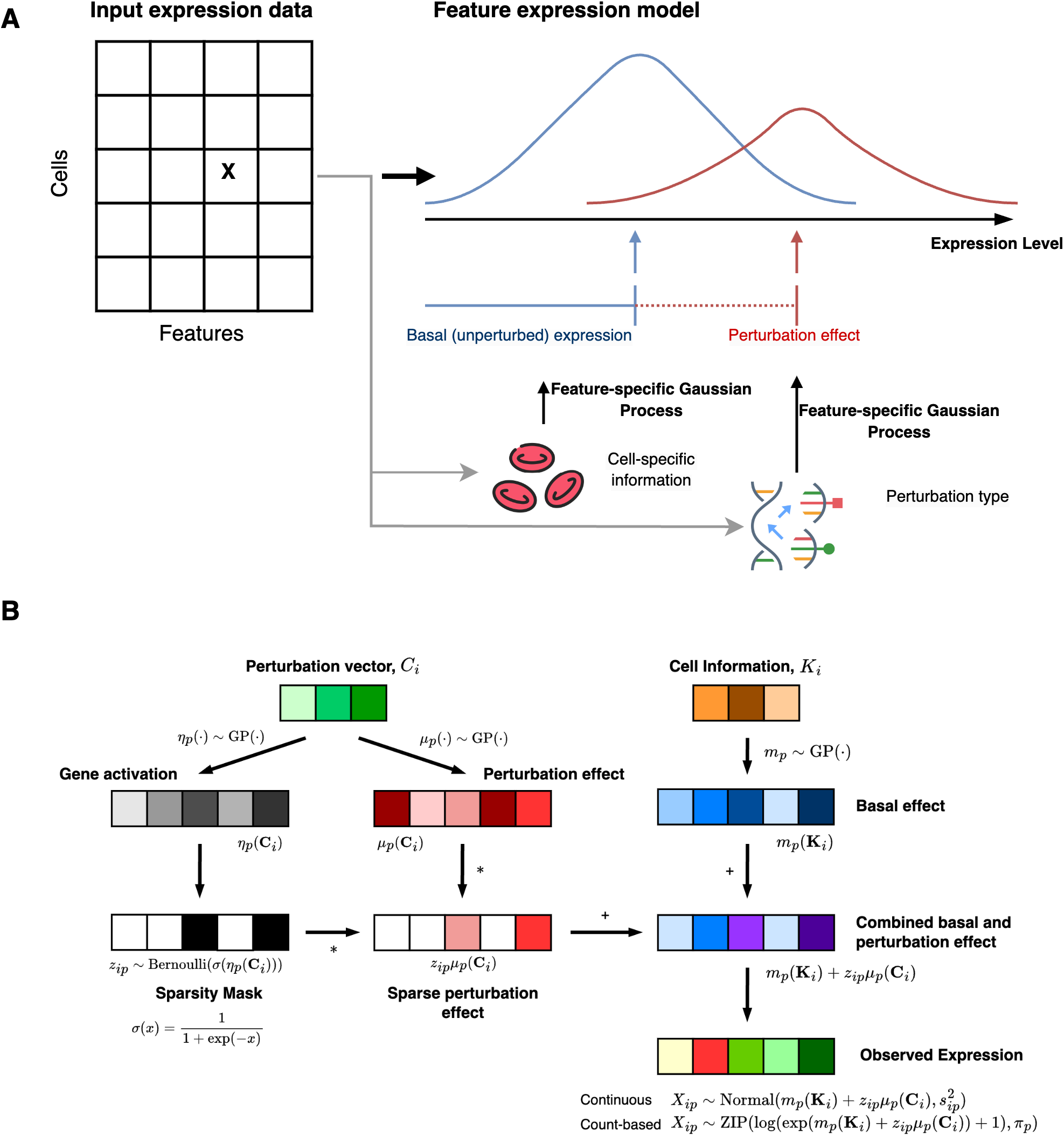
Overview of GPerturb. (A) For each cell-feature, **GPerturb** models the distribution over observed feature expression as the sum of a basal (unperturbed) expression and perturbation effect using feature-specific Gaussian Processes to transform of cell-specific information and perturbation applied. (B) Schematic of the mathematical construction of **GPerturb** showing the incorporation sparsity models and the ability to provide observation models for continuous and count-based expression data.

### Single-gene perturbation analysis

We first compared the predictive performance of GPerturb, CPA, GEARS and SAMS-VAE on a subset of the genome-wide CRISPR interference Perturb-seq dataset from Replogle et al. (2022) studied in Bereket and Karaletsos (2023) and Roohani et al. (2023). For all methods, the recommended settings are used. Since SAMS-VAE takes count-based data as inputs while CPA and GEARS require continuous expression inputs, we compare SAMS-VAE against **GPerturb**-ZIP and CPA and GEARS against **GPerturb**-Gaussian respectively. Similar to previous studies, we randomly select 20% of the dataset as the test set, and use the rest to train GPerturb.

We compared the *averaged* predictions and *averaged* observations for each of the unique perturbations. For CPA, SAMS-VAE and GEARS, we computed and store the *average* of 1,000 samples of reconstructed/predicted expressions drawn from the fitted model for each of the unique perturbations. Similarly, for **GPerturb**, we compute and store the *averaged* predicted mean expressions for each of the unique perturbations (i.e. averaged over all samples associated with a common perturbation). Table 1 shows the Pearson correlations between the predicted and observed expression levels for the perturbations which are illustrated in Figure 2A,C. We see **GPerturb**-ZIP attains better correlation than SAMS-VAE (*r*_GPerturb_ = 0.972, *r*SAMS-VAE = 0.944) for count-based inputs, while CPA-mlp achieved the best performance ahead of **GPerturb**-Gaussian and GEARS on continuous inputs (*r*_CPA-mlp_ = 0.984, *r*_GPerturb_ = 0.981, *r*_GEARS_ = 0.977).

**Table 1:**
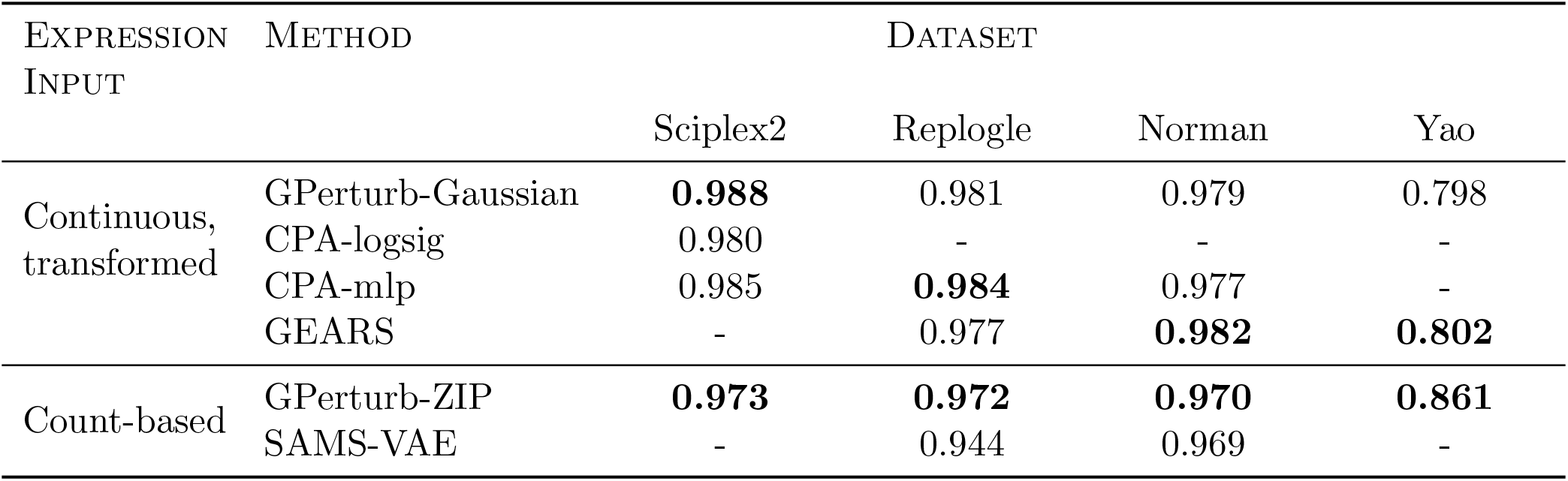
Comparison of predictive performance. Values show the Pearson correlation between predicted and observed expressions given by each method for each data set. GEARS and SAMS-VAE are not applicable to Sciplex2 due to non-binary perturbations. CPA and SAMS-VAE are not applicable to the multi-gene perturbation data from Yao et al (Yao et al., 2023) due to incompatible internal data preprocessing steps.

| EXPRESSION<br>INPUT | METHOD | DATASET |  |  |  |
| --- | --- | --- | --- | --- | --- |
|  |  | Sciplex2 | Replogle | Norman | Yao |
| Continuous,<br>transformed | GPerturb-Gaussian | **0.988** | 0.981 | 0.979 | 0.798 |
|  | CPA-logsig | 0.980 | - | - | - |
|  | CPA-mlp | 0.985 | **0.984** | 0.977 | - |
|  | GEARS | - | 0.977 | **0.982** | **0.802** |
| Count-based | GPerturb-ZIP | **0.973** | **0.972** | **0.970** | **0.861** |
|  | SAMS-VAE | - | 0.944 | 0.969 | - |

**Figure 2:**
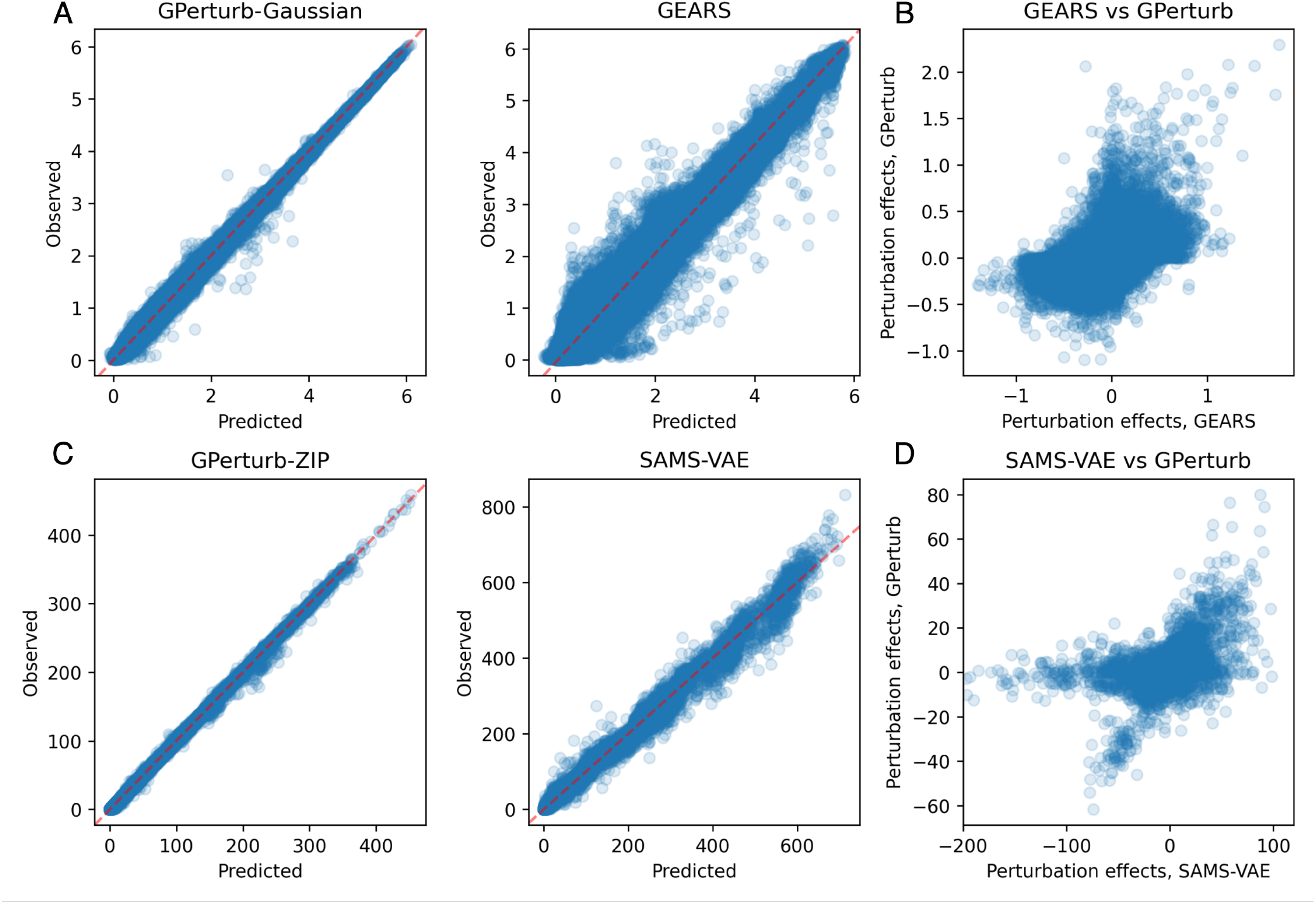
Single-gene perturbation analysis. (A) Comparison of predicted versus observed expression from **GPerturb** and GEARS using continuous expression inputs and (B) comparison of predicted perturbation effects. (C) Comparison of predictions from **GPerturb** and SAMS-VAE on count-based expression inputs and (D) comparison of predicted perturbation effects. Data from Replogle et al. (2022)

While the overall correlation between predicted and observed expression was high, Figure 2B,D shows that the directionality of the perturbation effects given by different models did not always agree, with instances where one method might report that a perturbation gives increased gene expression while another method indicates that the perturbation leads to decreased expression. We quantified this observation in Table 2 by examining the directionality agreement over all gene-perturbation pairs. Figure 3A shows the discrepancies in the exosome-related perturbation effects between **GPerturb**-Gaussian, CPA and GEARS for continuous expression input. In contrast, using count-based expression input, **GPerturb**-ZIP and SAMS-VAE showed greater consistency suggesting that the choice of pre-processing could have a considerable impact on perturbation modelling (Figure 3B). In order to further examine this, we were able to compare the outputs of **GPerturb**-Gaussian and **GPerturb**-ZIP on 345 perturbations grouped by pathways in Figure 4. This showed that given the same data set, the conversion from count-based to continuous-based expression (and the necessary changes in likelihood model in **GPerturb**) considerably changes the predicted perturbation effects.

**Table 2.**
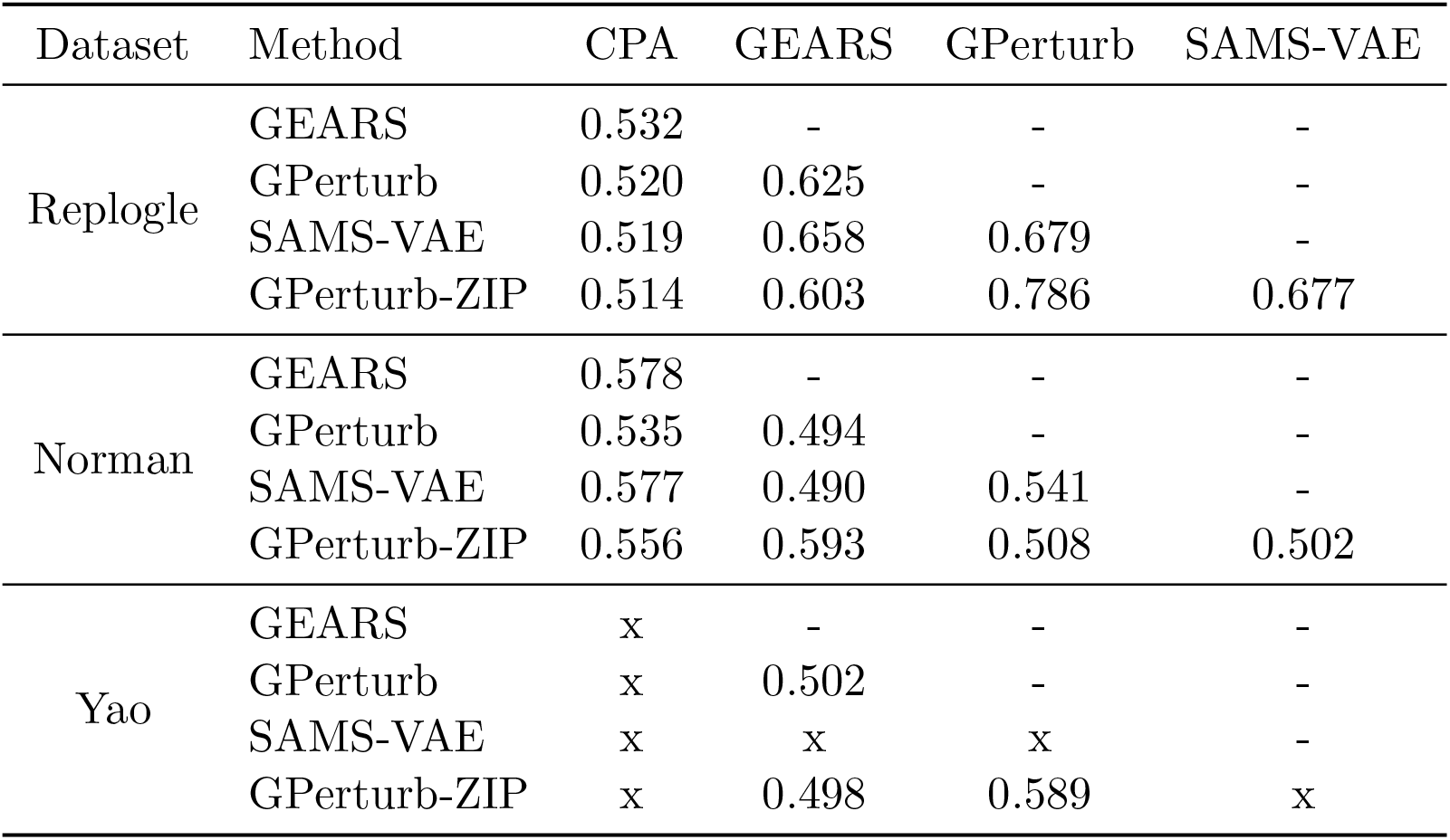
Proportion of gene-perturbation pairs with agreement on directionality between methods.

| Dataset | Method | CPA | GEARS | GPerturb | SAMS-VAE |
| --- | --- | --- | --- | --- | --- |
| Replogle | GEARS | 0.532 | - | - | - |
|  | GPerturb | 0.520 | 0.625 | - | - |
|  | SAMS-VAE | 0.519 | 0.658 | 0.679 | - |
|  | GPerturb-ZIP | 0.514 | 0.603 | 0.786 | 0.677 |
| Norman | GEARS | 0.578 | - | - | - |
|  | GPerturb | 0.535 | 0.494 | - | - |
|  | SAMS-VAE | 0.577 | 0.490 | 0.541 | - |
|  | GPerturb-ZIP | 0.556 | 0.593 | 0.508 | 0.502 |
| Yao | GEARS | x | - | - | - |
|  | GPerturb | x | 0.502 | - | - |
|  | SAMS-VAE | x | x | x | - |
|  | GPerturb-ZIP | x | 0.498 | 0.589 | x |

**Figure 3:**
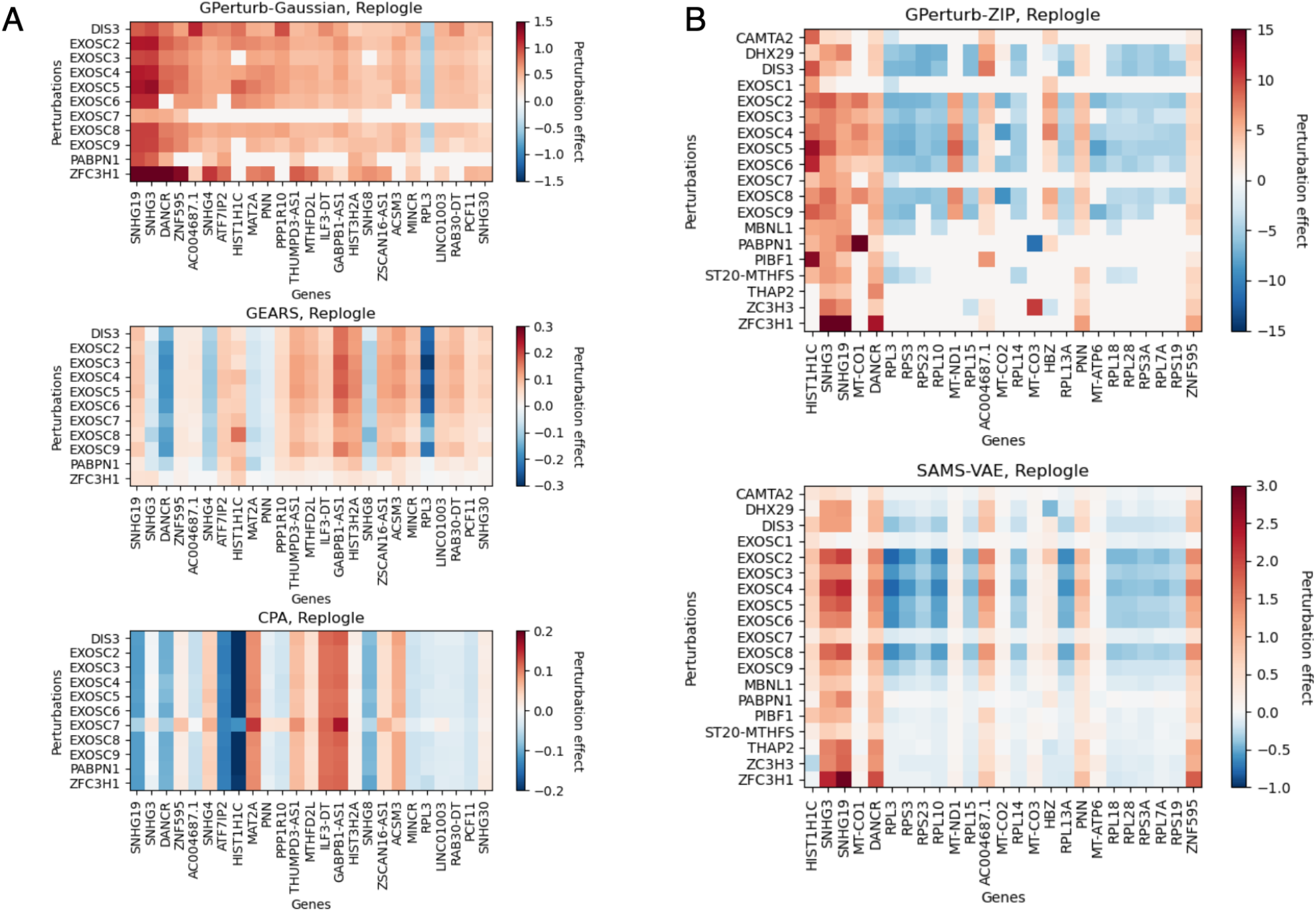
Differences in exosome-related perturbation effects associated with model and pre-processing selection. Top 25 most differentially expressed genes identified by (A) Gaussian and (B) Zero-Inflated Poisson versions of **GPerturb** and comparisons to perturbation effects inferred by GEARS, CPA and SAMS-VAE. Note that for the continuous case, we only include perturbations that are present in the GO graph of the current implementation of GEARS for sake of comparison. The difference in the scales of perturbation effects are due to the different internal data-preprocessing and normalising steps. Data from Replogle et al. (2022).

**Figure 4:**
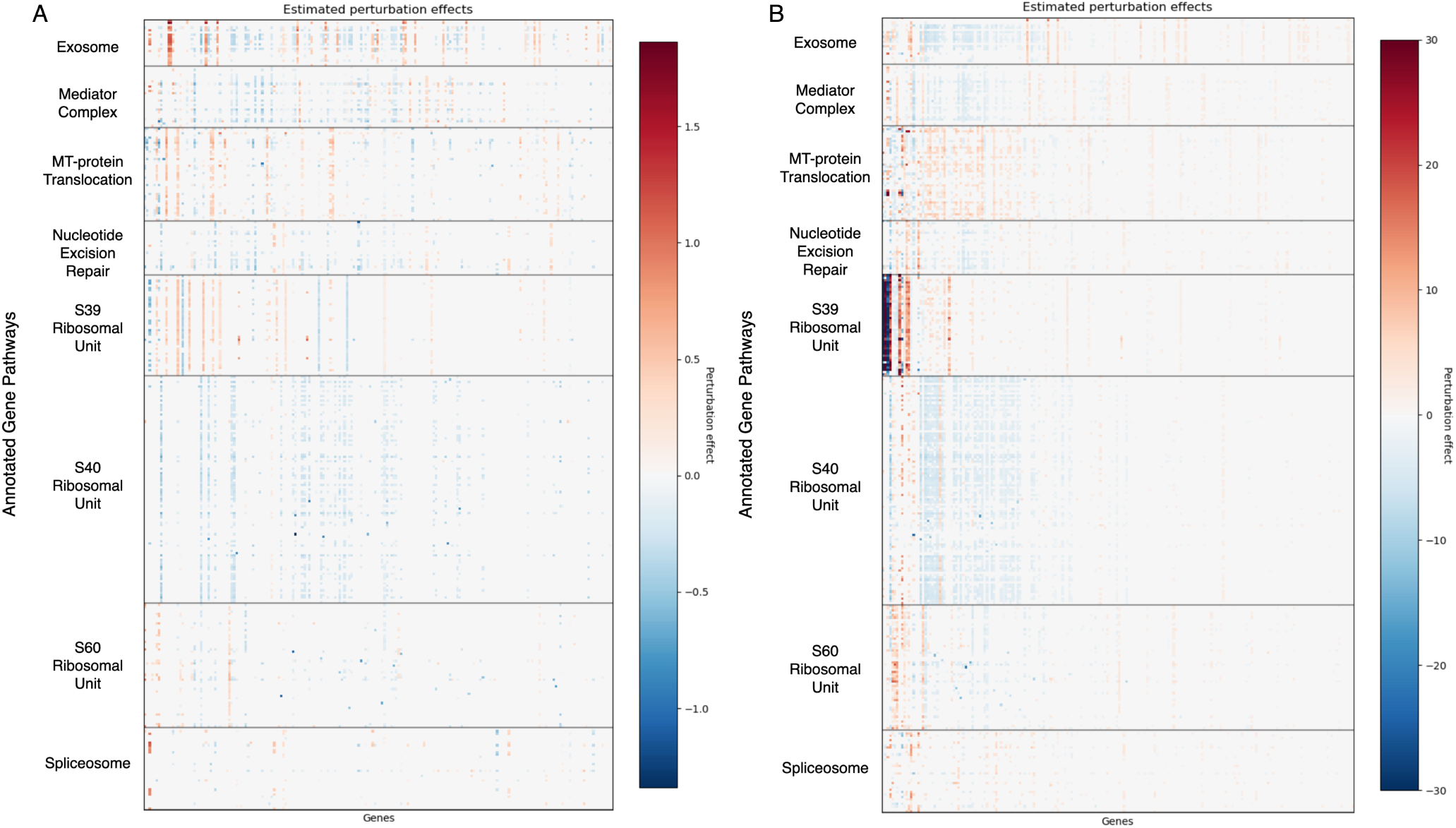
Perturbations by pathway annotations. Visualisation of 345 perturbations with pathway annotations and grouped by pathway highlights differences between (A) Gaussian- and (B) ZIP-based **GPerturb**. Each row corresponds to the perturbation effects of a unique perturbation. The perturbation effects for a gene is included only if the associated posterior inclusion probability greater than 0.95. Data from Replogle et al. (2022)

### Multi-gene perturbation analysis

We next considered a Perturb-seq dataset containing 131 two-gene perturbations from Norman et al. (2019) consisting of 89, 357 cells and 5, 045 genes. We compute the averaged predicted responses each method for each of the two-gene perturbations and compared to the corresponding averaged observations which are shown in Table 1 and Figure 5A,C. Note that unlike the other methods, GEARS is able to predict perturbation outcomes of previously unseen multi-gene perturbations by using biological knowledge encoded in its knowledge graph. Although **GPerturb** does not use additional prior information as in GEARS, it attains comparable correlation on predictions for two-gene perturbations and outperforms CPA and SAMS-VAE (Table 1). Interestingly, as in the previous experiments, the directionality of the perturbation effects between methods was not always consistent as illustrated in Figure 5B,D and quantified in Table 2.

**Figure 5:**
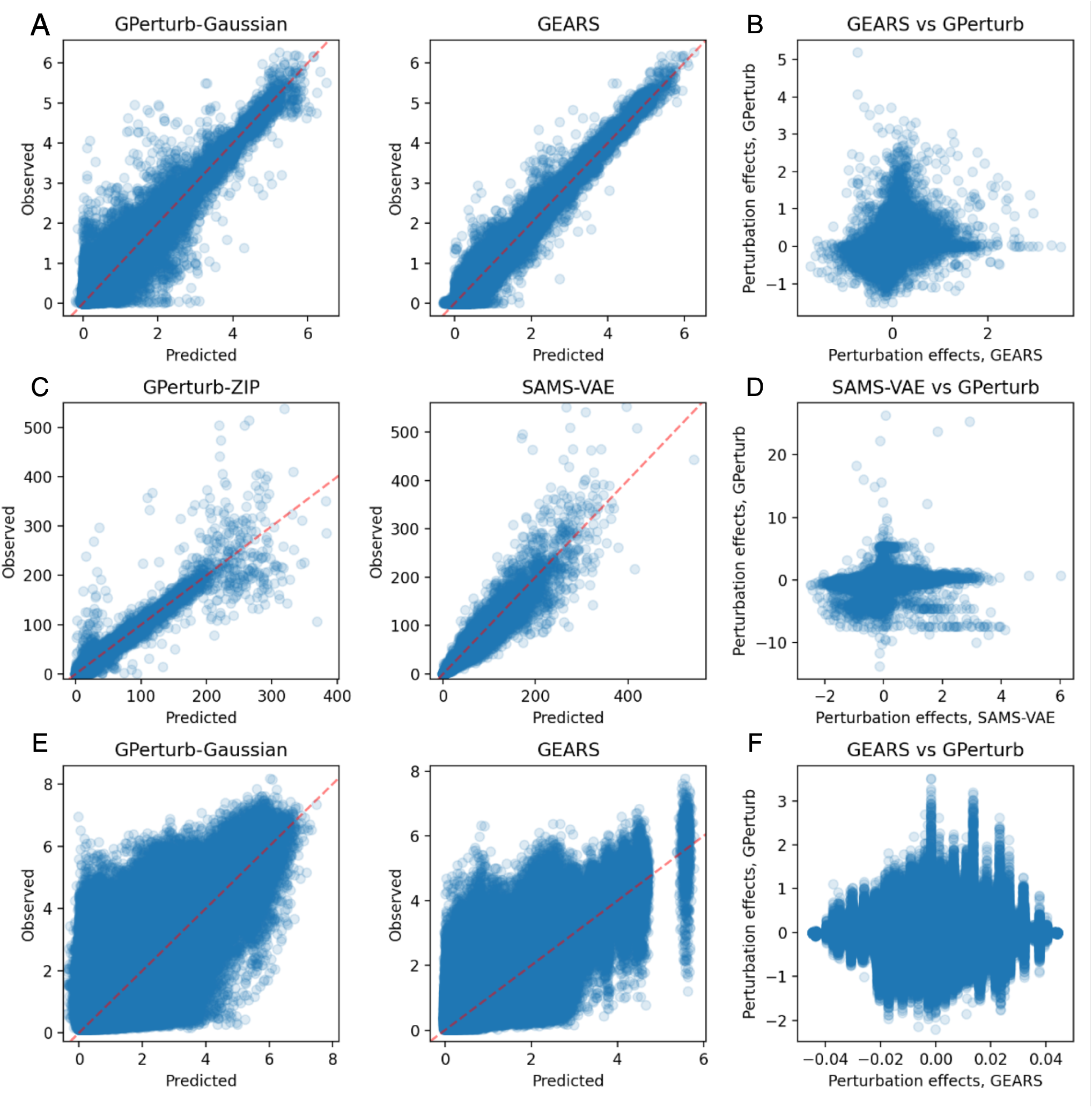
Multi-gene perturbation analysis. (A) Comparison of predicted and observed expression from **GPerturb** and GEARS using continuous expression inputs and (B) corresponding comparison of perturbation effects. (C) Comparison of predicted and observed expression from **GPerturb**-ZIP and SAMS-VAE using count-baed expression inputs and (D) corresponding comparison of perturbation effects. Data from Norman et al. (2019) (E) Comparison of predicted and observed expression from **GPerturb** and GEARS using continuous expression input and (F) corresponding comparison of perturbation effects. Data from Yao et al. (2023)

We further compare GEARS and **GPerturb** using the highly multiplexed Perturb-seq dataset from (Yao et al., 2023) under the same setup. CPA and SAMS-VAE could not be applied to this data set due to the large number perturbations. We report the averaged predictions against the corresponding averaged observations for both methods in Figure 5E. We see **GPerturb** and GEARS attain comparable predictions (*r*_GPerturb_ = 0.798, *r*_GEARS_ = 0.802) but predictions had high variance. Consequently, estimated perturbation effects for each perturbation-gene pair given by **GPerturb** and GEARS in Figure 5F showed weak correlation only. Unlike the previous examples, we see that even though the two methods have similar prediction accuracy, the scale of estimated perturbation effects given by GEARS is much smaller than **GPerturb**. The much more conservative perturbation estimations given by GEARS is likely due to the fact that less than 30% of the genetic perturbations in (Yao et al., 2023) is present in the Gene-Ontology (GO) knowledge graph in the current implementation of GEARS.

### Dosage-based perturbations

We next considered the SciPlex2 dataset (Srivatsan et al., 2020) where we examined a subset of A549 cells treated with one of the four compounds: dexamethasone (Dex), Nutlin-3a (Nutlin), BMS-345541 (BMS), or vorinostat (SAHA) across seven different doses. As a benchmark we conducted an analysis using CPA (Lotfollahi et al., 2023) which requires four inputs for each prediction: the cell property, a perturbation type, the expression profile of the cell corresponding to that perturbation and the perturbation type for which we want to predict the expression profile. We recorded the averaged counterfactual predictions of the negative control samples (no perturbation) under each of the 28 unique perturbations (4 compounds × 7 dosages) as counterfactual treatments. For **GPerturb** we recorded the averaged predictions (i.e. prediction values averaged over all cells associated with a common perturbation) for each of the 28 unique perturbations. We then compare the averaged predictions associated with all unique perturbations to the averaged observations in Table 1 and found **GPerturb** outperformed two variants of CPA: CPA-mlp and CPA-logsig. The latter enforces monotonicity of the dose-response relationship in its latent space. In the case of **GPerturb**, this was achieved in the absence of requiring a basal expression profile as input as needed in CPA. Note that comparison with GEARS and SAMS-VAE was not possible since neither account for non-binary perturbations. We then further investigated the ability of **GPerturb** to model the dosage relationships. Figure 6 illustrates that the predicted dosage-dependent expression levels given by **GPerturb** are more aligned to the measured expression values than both CPA variants particularly for non-monotonic dependencies between drug doses and expression levels. In particular, for PDE4D, CDKN1A and MDM2, expression varies non-monotonically for BMS which is not captured by the monotonicity constrained CPA-logsig.

**Figure 6:**
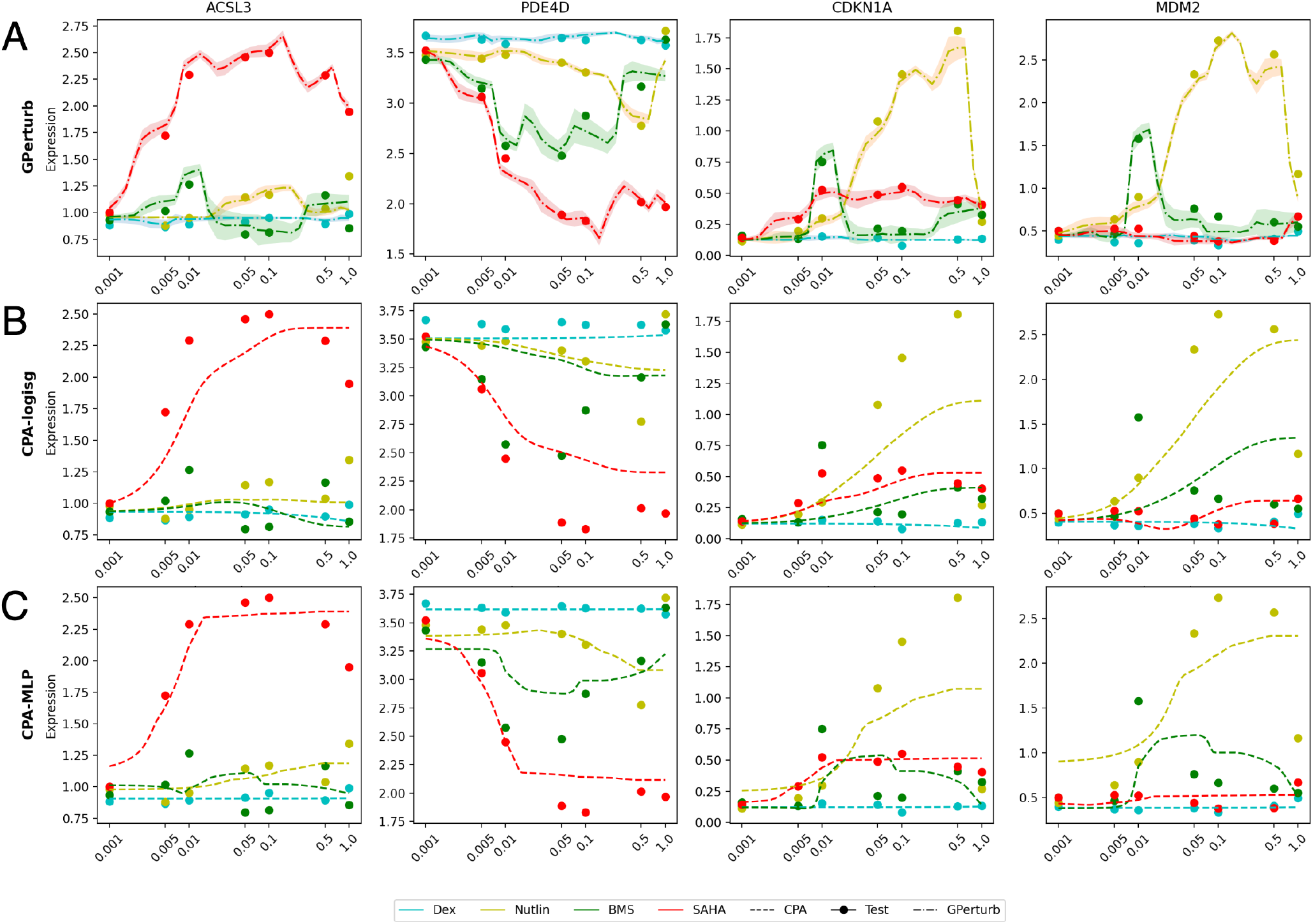
Analysis of continuous dosage-based perturbations. Predicted dosage-linked expression levels given by (A) **GPerturb**, (B) monotonically-constrained CPA and (C) unrestricted CPA on selected genes from the Sciplex2 datasetSrivatsan et al. (2020). Different colours are assigned to the four drugs (Dex, Nutlin, BMS, SAHA). Shaded regions are the corresponding 95% credible band given by GPerturb. Genes are selected to reproduce Fig S5 in Lotfollahi et al (Lotfollahi et al., 2023).

### Comparisons to linear models

In this section, we apply **GPerturb** to the LUHMES neural progenitor cell CROP-seq dataset studied in Zhou et al. (2023), and compare its performance to GSFA. This study targets 14 neurodevelopmental genes, including 13 autism risk genes, in LUHMES human neural progenitor cells. The raw data is preprocessed using the identical procedure described in Zhou et al. (2023). The resulting dataset consisting of *N* = 8, 708 samples and *P* = 6, 000 selected genes. The perturbations were encoded as one-hot vectors of length 14, each element corresponding to one of the 14 targeted neurodevelopmental genes (i.e. 14 distinct perturbations). The cell information is a real vector of length 4 (`lib_size`: number of total UMI counts, `n_features`: number of genes with non-zero UMI readings, `mt_percent`: percentage of mitochondrial gene expression and `batch`: batch ID). In addition to the one-hot perturbations, the dataset also consists of negative control gRNAs whose perturbations are encoded as zeros. For our proposed method, we randomly select 20% of the dataset as the test set, and use the rest to train **GPerturb**. For GSFA, the results are obtained based on the recommended settings given in Zhou et al. (2023).

We further note that in Zhou et al. (2023), the authors first removed cell level information from the continuous expression inputs by regressing the expression data on the cell information and then apply GSFA to the corresponding standardized residual matrix. In contrast, **GPerturb** disentangles and estimates cell-level and perturbation-induced variations simultaneously, and does not require any additional standardisation. We provided input into **GPerturb** with and without this standardisation but note that the analysis is more interpretable on the original form as additional transformations can affect prediction power and quality.

Figure 7A illustrates the predictions with **GPerturb** on the original data scale and after standardisation. While **GPerturb** shows good predictive performance without the GSFA standardisation applied to the data, it achieves similar correlative performance to GSFA when the data standardisation is used (Pearson correlation *r*_GPerturb_ = 0.248, *r*_GSFA_ = 0.182).

**Figure 7:**
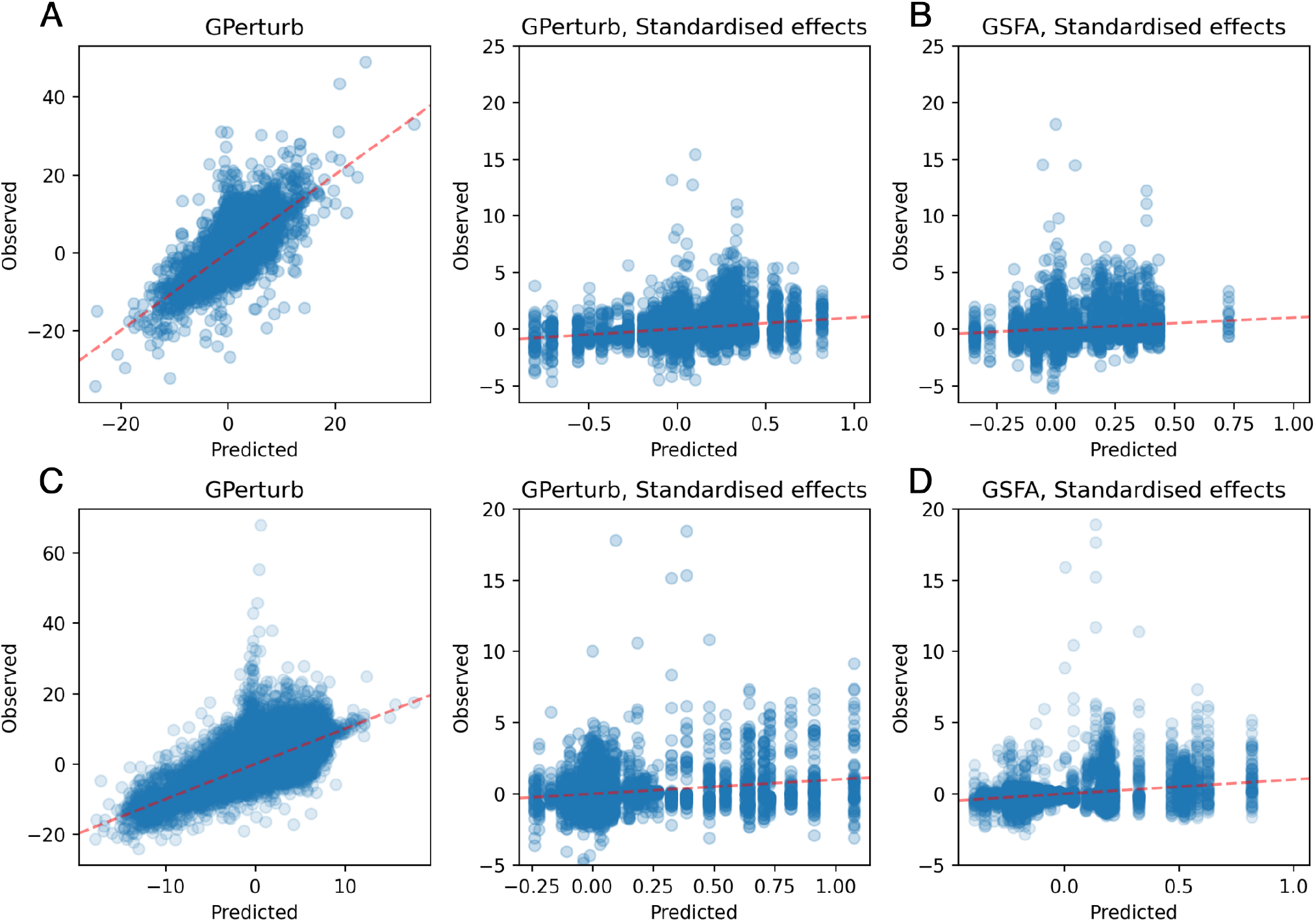
Comparison to GSFA. (A) Comparison of predicted and observed expression for LUHMES neural progenitor cell CROP-seq using **GPerturb** with and without GSFA-based standardisation and (B) GSFA reported predicted and observed standardised expression. (C) Comparison of predicted and observed expression for primary human CD8+ T cells datasets Zhou et al. (2023) given by **GPerturb** with and without GSFA-based standardisation and (D) GSFA reported predicted and observed standardised expression.

We also applied **GPerturb**-Gausain to the primary human CD8+ T cells dataset studied in Zhou et al. (2023) in a similar fashion. This study targets 20 genes associated with the T cell response, in both stimulated and unstimulated T cells. The processed dataset consists of *N* = 24, 955 samples and *P* = 6, 000 genes. The perturbations were encoded as one-hot vectors of length 20, which correspond to the 20 targeted genes in the study, and cell information was provided as a real vector of length 5 (`lib_size`: number of total UMI counts, `n_features`: number of genes with non-zero UMI readings, `mt_percent`: percentage of mitochondrial gene expression, `donor`: T Cell donor ID and `stimulated`: whether or not the T Cell is stimulated).

In Zhou et al. (2023), the authors hypothesised that perturbation effects are different in stimulated and unstimulated cells, and used a modified GSFA to capture such difference. In this example, we replicated the modification in **GPerturb** model to accommodate potentially different perturbation effects for stimulated and unsimulated T Cells in a similar fashion. Similar to the previous example, we randomly select 20% of the dataset as the test set, and use the rest to train GPerturb. We report the fitted results in Figure 7D. When comparing the predictive performance, GSFA showed greater correlation than **GPerturb** on the standardised data (*r*_GPerturb_ = 0.271, *r*_GSFA_ = 0.335). However, Figure 7C shows that the standardisation applied by GSFA was likely to be disadvantageous and unnecessary with **GPerturb** since it can be applied without the initial cell information regression step.

## Discussion

There are a number of state-of-the-art single-cell perturbation modelling methods currently available (including many not directly considered here) but a detailed analysis of the preprocessing, training and inference requirements of each method highlights significant differences in the approach and requirements associated with each method. While there has been considerable interest in deep learning based approaches, **GPerturb** adopts a more classical non-linear regression based offering which provides a non-deep learning approach to support model training and prediction by focusing on directly modelling individual genes rather than via the use of latent representations in many other recent methods. Our analysis shows that **GPerturb** is capable of state of the art performance despite these significant design differences and is highly versatile and computationally efficient (see Supplementary Table 1). A feature of our experimental results (Table 1) is that **GPerturb** in both forms could be applied to all four examples, while other methods could only be used for a subset of these, **GPerturb** is able to handle single, multi-gene and continuous perturbation inputs.

Our experiments show that direct performance comparisons between methods must be interpreted carefully and may not always be applicable. For example, comparisons between GSFA and other methods were not shown since GSFA operates and returns results in terms of standardised input data residuals. In addition, we also tried to compare the estimated sparse perturbation effects given by **GPerturb** with SAMS-VAE and CPA, but found no straight-forward way to do so due to the fact that **GPerturb** directly estimates sparse perturbation effects on the gene level, while CPA and SAMS-VAE focus on finding sparse low-dimensional embeddings of them. Furthermore, since our proposed **GPerturb** framework allows handling of both continuous normalised and count-based data using Gaussian and zero-inflated Poisson based likelihoods, we have observed that while the alternate versions of **GPerturb** attain comparable prediction accuracy with methods using comparable input data, the perturbation effects captured by the Gaussian and ZIP versions of **GPerturb** could be different. This highlights that variations in data processing and modelling could affect the conclusions drawn from the same raw data.

Our experiments are consistent with other recent more extensive evaluation studies, such as Li et al. (2024a,b); Ahlmann-Eltze et al. (2024), which have also found that prediction performance is highly context-dependent and that no single method excels across all scenarios. These evaluations includes recently developed single-cell foundation models which can also be applied for perturbation effect prediction. In some cases performance may be no better than simple linear models. The scalable Gaussian Process regression models we have introduced in **GPerturb** provide a highly-effective and complementary approach for single-cell perturbation modelling. These models can be of utility for direct prediction tasks or as a methodologically distinct benchmark for the development of new methods. Future work could examine extensions of this Gaussian Process framework as a credible non-deep learning based approach for handling multi-omics or spatially resolved molecular data.

## Methods

### GPerturb-Gaussian

We first discuss the model with **X** being a matrix of pre-processed continuous responses. We will give a counting data version of the model alongside with a schematic illustration of the additive modelling structure later.

Let 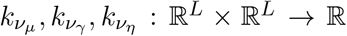 be Gaussian process kernels governed by kernel parameters *ν*_*µ*_, *ν*_*γ*_, *ν*_*η*_ respectively. Let *g*_*µ*_, *g*_*γ*_, *g*_*η*_ : ℝ^*L*^ → ℝ be the mean functions of the corresponding Gaussian processes.

We first define the gene-level additive perturbation model as follows:

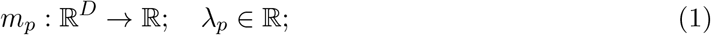

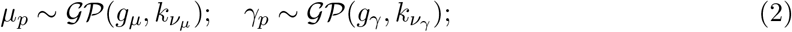

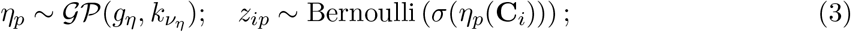

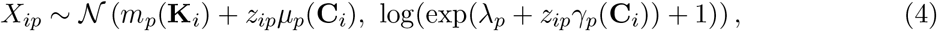

where 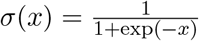.In this model setup, *m*_*p*_ is a fixed but unknown function that takes the cell-level information vector **K**_*i*_ associated with the *i*th sample as input, and returns *m*_*p*_(**K**_*i*_) as the expected basal expression level of the *p*th gene in the *i*th sample. *λ*_*p*_ is the basal variability parameter of the expression level of the *p*th gene shared across all samples *i* = 1,…, *N*. *z*_*ip*_ is a binary toggle controlling whether or not the expression level of the *p*th gene in the *i*th sample is perturbed by the perturbation vector **C**_*i*_. The success probability of *z*_*ip*_ depends on **C**_*i*_ through *η*_*p*_(**C**_*i*_), a random function *η*_*p*_ evaluated at **C**_*i*_ (Note that under our additive modelling setup, the binary toggles *z*_*ip*_ are the same for all cells receiving the same perturbation). *µ*_*p*_, *γ*_*p*_ are also random functions that take **C**_*i*_ associated with the *i*th sample as input, and return *µ*_*p*_(**C**_*i*_), *γ*_*p*_(**C**_*i*_) as the potential mean- and variability-level perturbation effects on the expression level of the *p*th gene in the *i*th sample. Schematic illustration and graphical representation of the Gaussian model is given in Supplementary Fig. 1.

We then assume the observed perturbed expression level *X*_*ip*_ associated with perturbation **C**_*i*_ and cell-level information **K**_*i*_ follows a Gaussian distribution with mean being the sum of basal mean *m*_*p*_(**K**_*i*_) and mean-level perturbation effect *z*_*ip*_*µ*_*p*_(**C**_*i*_), and variance being a positive function of the sum between the common basal variability *λ*_*p*_ and a variability-level perturbation *z*_*ip*_*γ*_*p*_(**C**_*i*_). We choose to use the function log(exp(·) + 1) to map the unconstrained variability parameters to the positive real variance parameters since it is approximately linear when the magnitude of input is large, and therefore would partially retain the additive structure between the basal states and perturbation effects in a similar fashion to the mean parameters in comparison with e.g. exp(·).

Under our modelling setup, the parameters naturally partition into two groups: the random perturbation-specific parameters 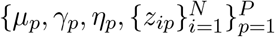, and the unknown but fixed basal level parameters 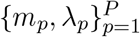. We do not treat the basal level parameters as random variables since the primary objective of the model is to learn how the gene expression levels *X*_*ip*_ respond to different perturbations. In the proposed model, the basal states only play the role of “intercept” or “offset”, and is not of primary interest to us. In addition, treating the basal level parameters as unknown but fixed model parameters also simplifies the inference procedure and reduces the computational cost of the proposed model.

### GPerturb-ZIP

We now discuss the model for expression count data. Similar to the continuous model, we define the count data based gene-level additive perturbation model using a zero-inflated Poisson likelihood as follows

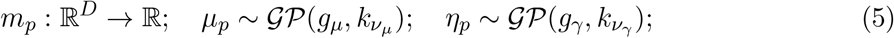

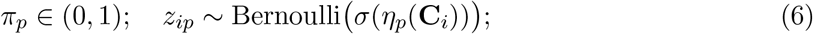

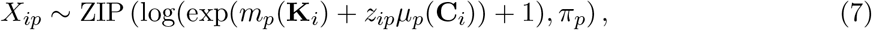

where *µ*_*p*_, *m*_*p*_, *η*_*p*_, *z*_*ip*_ have the same interpretation as in the deviance-based model, *π*_*p*_ is the proportion of excessive zeros on the *p*th gene shared across all samples *i* = 1,…, *N*, and ZIP(*µ, π*) denotes a zero-inflated Poisson distribution with expected Poisson rate *µ* and probability of excessive zeros *π*. Note that our ZIP model does not aim to estimate the pattern of excessive zeros of the dataset. Hence the quantity *z*_*ip*_*µ*_*p*_(**C**_*i*_)) should be interpreted as “the conditional perturbation effect given that the corresponding observation *X*_*ip*_ is not an excessive zero”. Schematic illustration and graphical representation of the Zero-inflated Poisson model is given in Supplementary Fig. 2.

We also considered handling potential over-dispersion by modelling **X** using a zero-inflated Gamma-Poisson likelihood (a different parameterisation of Negative Binomial). However, we find the Gamma-Poisson model and Poisson model achieved similar level of prediction performance on real datasets, and the majority of estimated dispersion parameters are far less than 1, showing no strong sign of over-dispersion. Hence we focus on the Poisson model in this section for sake of simplicity. The details of the zero-inflated Gamma-Poisson model can be found in **Supplementary Information**.

For both the deviance-based Gaussian and the Zero-inflated Poisson model, we recommend setting *k*_*µ*_, *k*_*γ*_, *k*_*η*_ to be RBF kernels 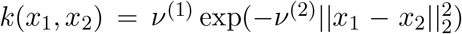 governed by kernel parameters 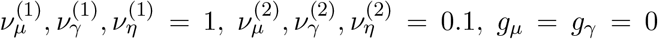 and *g* = −3 as these prior specifications give satisfactory results in all of our numerical experiments. These choices of priors reflect our belief that all *µ*_*p*_(**C**_*i*_) and *γ*_*p*_(**C**_*i*_) have the same marginal prior N (0, 1), and the prior on *σ*(*η*_*p*_(**C**_*i*_))), the inclusion probability of perturbation effect of **C**_*i*_ on the *p*th gene, is concentrated at around 0.05. Alternative choices are also discussed in the following sections.

### Posterior inference

In this section, we discuss the posterior inference strategy of the proposed models. We first give the posterior inference procedure of the deviance-based model. Let 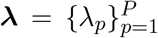.Let 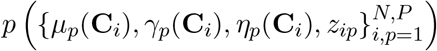 be the prior of the associated perturbation-specific parameters. Let

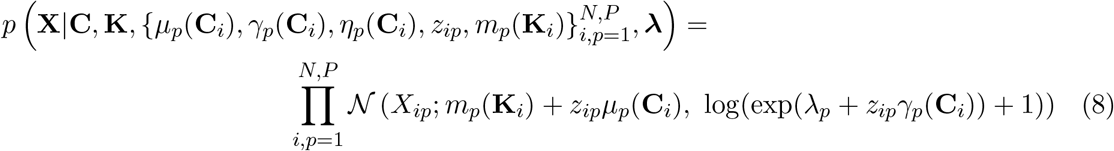

be the likelihood of the observed gene-expression level matrix **X** given the perturbation matrix **C**, the cell-level information matrix **K**, and all model parameters. Since the number of samples *N* and the number of genes *P* are usually large, jointly estimating the posterior 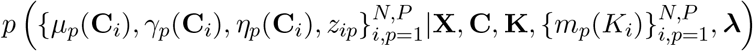 is computationally infeasible. We therefore use armortised variational inference (Ganguly et al., 2023) to address this issue: Let *f*_***ξ***_ : ℝ^*L*^ → ℝ^6*P*^ be a neural network parameterized by a real vector ***ξ***. We approximate the full posterior 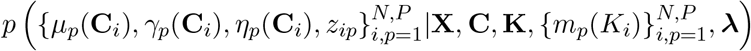 using the following variational posterior:

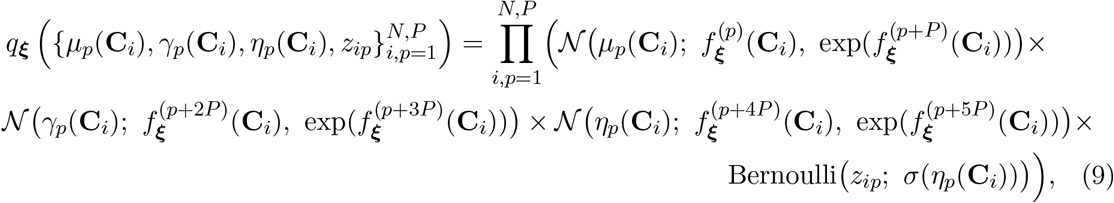

where 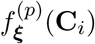 denotes the *p*th entry of *f*_***ξ***_(**C**_*i*_), 𝒩 (·; *µ, s*^2^) denotes a Gaussian p.d.f. with mean *µ* and variance *s*^2^, and Bernoulli(·; *π*) denotes a Bernoulli p.m.f. with success probability *π*. Similarly, for the fixed but unknown basal level functions 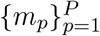,we let *f*_***ϕ***_ : ℝ^*D*^ → ℝ^*P*^ be a neural network parameterized by a real vector ***ϕ***, and use 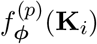 to parameterize *m*_*p*_(**K**_*i*_) for all *i* = 1,…, *N* and *p* = 1,…, *P*. The evidence lower bound (ELBO) of the deviance-based model then takes the form

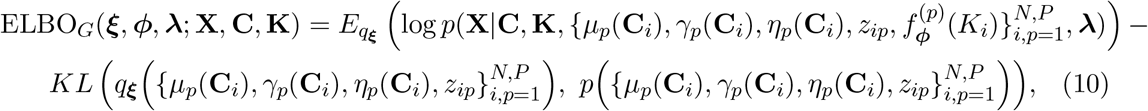

where 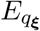 denotes expectation with respect to the variational posterior *q*_***ξ***_. We estimate the variational posterior *q*_***ξ***_ and all other model parameters by maximizing (an empirical version of) ELBO_*G*_(***ξ, ϕ, λ***; **X, C, K**). The Bernoulli random variables in 9 is approximated using Gumbel softmax (Jang et al., 2016).

Let {***ξ***^*^, ***ϕ***^*^, ***λ***^*^} = arg max_***ξ***,***ϕ***,***λ***_ ELBO_*G*_(***ξ, ϕ, λ***; **X, C, K**). Once the model has been fitted, we can then construct both approximate point and interval estimates of parameters of our interest. For example, let **C**^′^ be a generic perturbation vector. One can form approximate point or interval estimates of the posterior inclusion probability *σ*(*η*_*p*_(**C**^′^)), which controls if the expression level of the *p*th gene is perturbed by **C**^′^, using the variational posterior 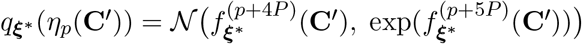. Compared with LSFR used in GSFR, identifying perturbation effects using posterior inclusion probability is more intuitive and interpretable thanks to the full Bayesian framework of the proposed model.

We now discuss the posterior inference of the Zero-inflated Poisson model. It can be parameterized and estimated in a similar fashion to the deviance-based model: Let 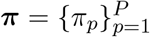. Let 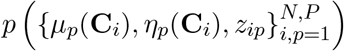 be the prior of the associated perturbation-specific parameters in the Zero-inflated Poisson model. Let

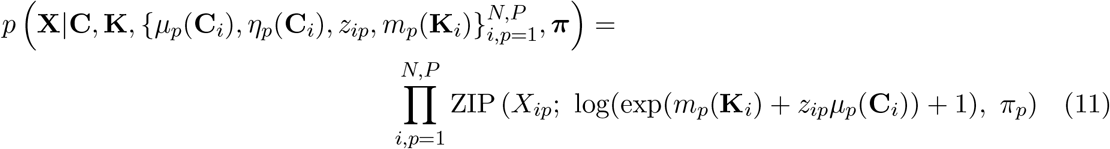

be the likelihood of the raw counting data **X**, where ZIP(·; *µ, π*) is the p.m.f. of a Zero-Inflated Poisson distribution with Poisson rate *µ* and probability of excessive zeros *π*. Similar to the deviance-based model, let *f*_***θ***_ : ℝ^*L*^ → ℝ^4*P*^ be a neural network parameterized by a real vector ***θ***. Let

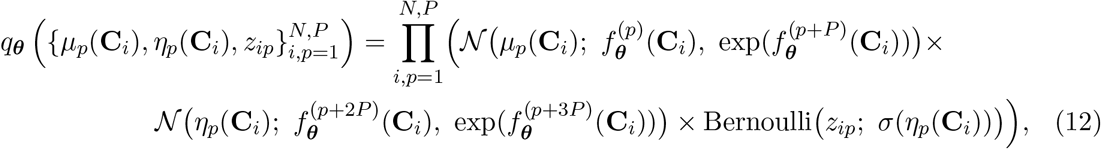

be the variational posterior of the perturbation-specific parameters 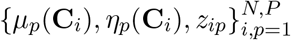. As in the deviance-based model, we use *f*_***ϕ***_(*K*_*i*_) to parameterize the basal level parameter *m*_*p*_(*K*_*i*_) for all *i* = 1,…, *N* and *p* = 1,…, *P*. Then the evidence lower bound of the Zero-inflated Poisson model ELBO_*P*_ (***θ, ϕ, π***; **X, C, K**) is defined in a similar fashion to Eqn (10), and the model parameters are estimated by maximizing (an empirical version of) ELBO_*P*_ (***θ, ϕ, π***; **X, C, K**) with respect to {***θ, ϕ, π***}.

### Magnitudes of perturbation vectors

In our proposed methods, the perturbation vector **C**_*i*_ can either be binary (indicating the presence of a perturbation) or continuous (representing e.g. dosage). When **C**_*i*_ represents the continuous dosage of a perturbation, we expect that the potential perturbation effects is positively correlated to the dosage (at least in a sensible range before some ceiling effects). Similarly, when **C**_*i*_ = **0** (i.e. no perturbation at all), we expect there is no potential perturbation effects. To impose these physical constraints, we recommend modifying the model and inference procedure as follows (Here we focus on the deviance based model. The zero-inflated Poisson model can be modified in a similar fashion): We first replace the standard RBF kernels on *ν*_*µ*_, *ν*_*γ*_ by a modified “zero-passing” RBF kernel used in Xing and Yau (2024). This modification ensures that samples 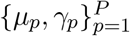 drawn from the modified Gaussian process prior would satisfy 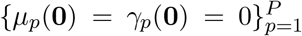. We also replace the generative process of *z*_*ip*_ with 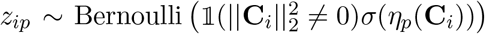,which ensures that no *z*_*ip*_ would be triggered when the input **C**_*i*_ = **0**. This choice also reflects our prior belief that even though the potential effects of a perturbation **C**_*i*_ depends on its magnitude, whether or not a gene is perturbed only depends on the presence of the perturbation (i.e. 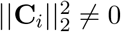), but not the scale of it. Similar constraints are also used in e.g. Lotfollahi et al. (2023) and Bereket and Karaletsos (2023).

In addition to the generative process, we also adjust the inference procedure accordingly. We modify the variational posterior in (9) as follows:

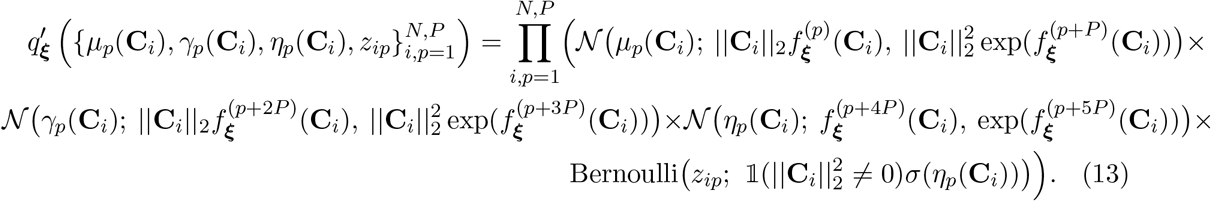

Here we rescale the variational posterior of the mean- and viability-level perturbation by a factor ||**C**_*i*_||_2_. This ensures that both terms would be zero when there is no perturbation, and would explicitly depend on the size of **C**_*i*_ otherwise. We also modify the variational distribution of *z*_*ip*_ in the same way as in the generative process. These modification ensures that both generative process and posterior inference are inline with the physical constraints discussed above. We use this modified generative process and inference procedure as our default model setup for the rest of the paper.

### Single gene perturbation

The Perturb-Seq dataset from Replogle et al. (2022) was pre-processed and filtered using the pre-processing steps described in Lopez et al. (2023) and Bereket and Karaletsos (2023). The resulting dataset **X** ∈ ℕ^*N* ×*P*^ consists of counting data of *N* = 118, 461 cells and *P* = 1, 187 genes. For *i* = 1,…, *N*, the perturbation **C**_*i*_ ∈ ℝ^*L*^ is either a length *L* = 722 one-hot vector, representing one of the 722 unique CRISPR guides (perturbations), or a zero vector, representing the perturbation associated with negative controls (non-targeting CRISPR guides). The non-targeting negative controls are treated as the baseline level. The cell information **K**_*i*_ ∈ ℝ^*D*^ is a length *D* = 4 real vector (`lib_size`: total number of UMI counts, `n_features`: number of genes with non-zero UMI readings, `mt_percent`: percentage of mitochondrial gene expression, `scale_factor`: core scale factor).

### Multigene perturbation prediction

We compared GPerturb’s performance on predicting multigene perturbation outcomes with the knowledge-graph informed GEARS using the Perturb-seq dataset (Norman et al., 2019) studied in Roohani et al. (2023). We followed the same data-preprocessing process used in Roohani et al. (2023). The resulting dataset **X** ∈ ℝ^*N* ×*P*^ consists of *N* = 89, 357 cells and *P* = 5, 045 genes. For *i* = 1,…, *N*, the perturbation **C**_*i*_ ∈ ℝ^*L*^ is a length *L* = 103 binary vector where the positions of ones encode the perturbed genes. (The dataset consists of 131 two-gene perturbations.) The cell information **K**_*i*_ ∈ ℝ^*D*^ is a length *D* = 2 real vector (`lib_size`: total number of UMI counts, `n_features`: number of genes with non-zero UMI readings). We randomly select 20% of the dataset as the test set, and use the rest as training set. For both our GPerturb and GEARS, the recommended settings are used to fit the models.

We further compared GPerturb with GEARS using the multiplexed Perturb-seq dataset from Yao et al. Yao et al. (2023) using the same procedure described above. We follow the data-preprocessing process given by the authors. The resulting dataset **X** ∈ ℝ^*N* ×*P*^ consists of *N* = 24, 192 cells and *P* = 15, 668 genes. For *i* = 1,…, *N*, the perturbation **C**_*i*_ ∈ ℝ^*L*^ is a length *L* = 600 binary vector where the positions of ones encode the perturbed genes. The cell information **K**_*i*_ ∈ ℝ^*D*^ is a length *D* = 3 real vector (`lib_size`: total number of UMI counts, `n_features`: number of genes with non-zero UMI readings and `mt_percent`: percentage of mitochondrial gene expression). We compare the performance of GEARS and GPerturb under the same setting described above.

### SciPlex2

The SciPlex2 dataset (Srivatsan et al., 2020) (GSM4150377) consists of A549 cells treated with one of the four compounds: dexamethasone (Dex), Nutlin-3a (Nutlin), BMS-345541 (BMS), or vorinostat (SAHA) across seven different doses. We follow the data pre-processing steps given in Lotfollahi et al. (2023). The resulting dataset **X** ∈ ℝ^*N* ×*P*^ consists of *N* = 20, 643 cells and *P* = 5, 000 genes. For *i* = 1,…, *N*, the perturbation **C**_*i*_ ∈ ℝ^*L*^ is a length *L* = 4 vector with only one non-zero entry whose position and value encode the compound type and dosage respectively. Similar to the previous sections, the perturbation associated with negative controls are encoded as **C**_*i*_ = **0** and treated as the baseline level. The cell information **K**_*i*_ ∈ ℝ^*D*^ is a length *D* = 2 real vector (`lib_size`: total number of UMI counts, `n_features`: number of genes with non-zero UMI readings). We randomly select 20% of the dataset as the test set, and use the rest to train GPerturb. For both Gaussian GPerturb and CPA, the recommended settings are used to fit the models.

## Supporting information

Supplementary figures, discussion and experiments

## Data Availability

The Sciplex2 dataset with associated metadata were obtained from https://www.ncbi.nlm.nih.gov/geo/query/acc.cgi?acc=GSM4150377. The raw Perturb-Seq dataset from Replogle et al. Replogle et al. (2022) was obtained from https://gwps.wi.mit.edu. The raw Perturb-Seq dataset from Norman et al. Norman et al. (2019) was obtained from https://dataverse.harvard.edu/api/access/datafile/6154020. The raw Perturb-Seq dataset from Yao et al. Yao et al. (2023) was obtained from https://www.ncbi.nlm.nih.gov/geo/query/acc.cgi?acc=GSE221321. All processed data, count matrices and results (including pre-trained models and visualisation) are available at https://figshare.com/articles/dataset/GPerturb_Gaussian_process_modelling_of_single-cell_perturbation_data/26491588.

## Code Availability

Python code for **GPerturb** and reproducible examples of analyses conducted in this study can be found on the Github repository (https://github.com/hwxing3259/GPerturb). All code is made available under MIT license.

## Acknowledgements

The authors ares supported by an EPSRC Turing AI Acceleration Fellowship (Grant Ref: EP/V023233/1).

## Supplementary Information

### Preamble

This supplementary document is structured as follows: In Section S1, we give further details into the background and setup of the problem expanding on the Methods description. In Section S2, we discuss its connection to some existing state-of-the-art methods. We then study the properties of the method in Section S3 using simulated data. Finally, in Section S4, we provide further details of the analysis of a number of datasets including some not included in the main manuscripts, and compare its performance with state-of-the-art methods discussed in Section S2.

#### S1 Further background and motivation

Our research is motivated by guided sparse factor analysis (GSFA) ^11^, a Bayesian sparse factor analysis model which aims to infer both the effects of genetic perturbations on individual genes, and the groups of genes or gene modules that are co-regulated. In GSFA, the groups of co-regulated genes are encoded as sparse loading vectors, and the effects of genetic perturbations on individual genes are assumed to be linear functions of the encoded loading vectors and the perturbation vectors. Before we give the details, we first introduce the notation. Let *N* be the total number of samples (usually cells). For *i* = 1,…, *N*, let **K**_*i*_ be the *D* dimensional cell-level information vector, **C**_*i*_ the *L* dimensional perturbation vector, and **X**_*i*_ the *P* dimensional observed gene-expression vector associated with the *i*th sample, where *P* is the total number of genes. Each entry *X*_*ip*_ in **X** represents the observed expression level of the *p*th gene in the *i*th sample. Let 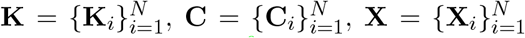.In GSFA, the authors first apply a deviance-statistics transformation^9^ to the raw counting data matrix, which returns to a continuous response matrix **X** ∈ ℝ^*N* ×*P*^. Then the cell-level information is decoupled/removed from **X** by first regressing each column of **X** on **K** using linear regression, then subtract the fitted value from **X**. In other words, the resulting matrix 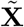 would be the residuals of the linear regressions described above. Finally, the author modelled the pre-processed 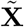 as

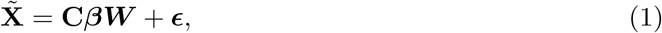

where *d* is the number of latent factors chosen by the user, ***β*** ∈ ℝ^*L*×*d*^ is a linear transformation of **C, C*β*** is the factor matrix informed by the perturbation vector **C, *W*** ∈ ℝ^*d*×*P*^ is the loading matrix, and ***ϵ*** ∈ ℝ^*N* ×*P*^ is the residual matrix.

To improve interpretability, the authors put sparsity inducing priors on ***β*** and ***W***. Even though GSFA gives promising results on multiple real-world datasets, it suffers from a few limitations. Firstly, we do not expect a perturbation **C**_*i*_ to affect all *P* genes in a dataset simultaneously. In particular, users are often interested in the questions “Given a perturbation, which subset of genes does it target?” Hence it would be desirable if a model could directly give sparse estimates of gene-level perturbation effects. However, even though the posterior of both ***β*** and ***W*** in GSFR are sparse, the resulting estimated perturbation effects on individual genes are not necessarily so, and users have to test for non-zero perturbation effects using LSFR^8^, which is not straightforward to interpret. Secondly, GSFR is only applicable to continuous response matrix **X**, which relies on some form of data-preprocessing. In practice, it is also of interest to study datasets in different formats. For example, users may be interested in directly analyzing the unique molecular identifier (UMI) counts instead of the continuous deviance statistics or *z*-scores. But it is not straightforward to generalize GSFR to non-Gaussian likelihoods in a computationally efficient way, which restricts GSFR’s applicability. In addition, the linear assumption in GSFR may not be flexible enough to capture the complex biological process behind the experiment results, and therefore would affect its prediction power. In the following section, we propose a model to address these limitations by directly estimating sparse gene-level perturbation effects using a Gaussian process based additive model and armortized variational inference.

##### S1.1 GPerturb Schematic Models

We present schematic illustrations of the Gaussian and ZIP-GPerturb in Fig 1 and 2.

**Supplementary Figure 1:**
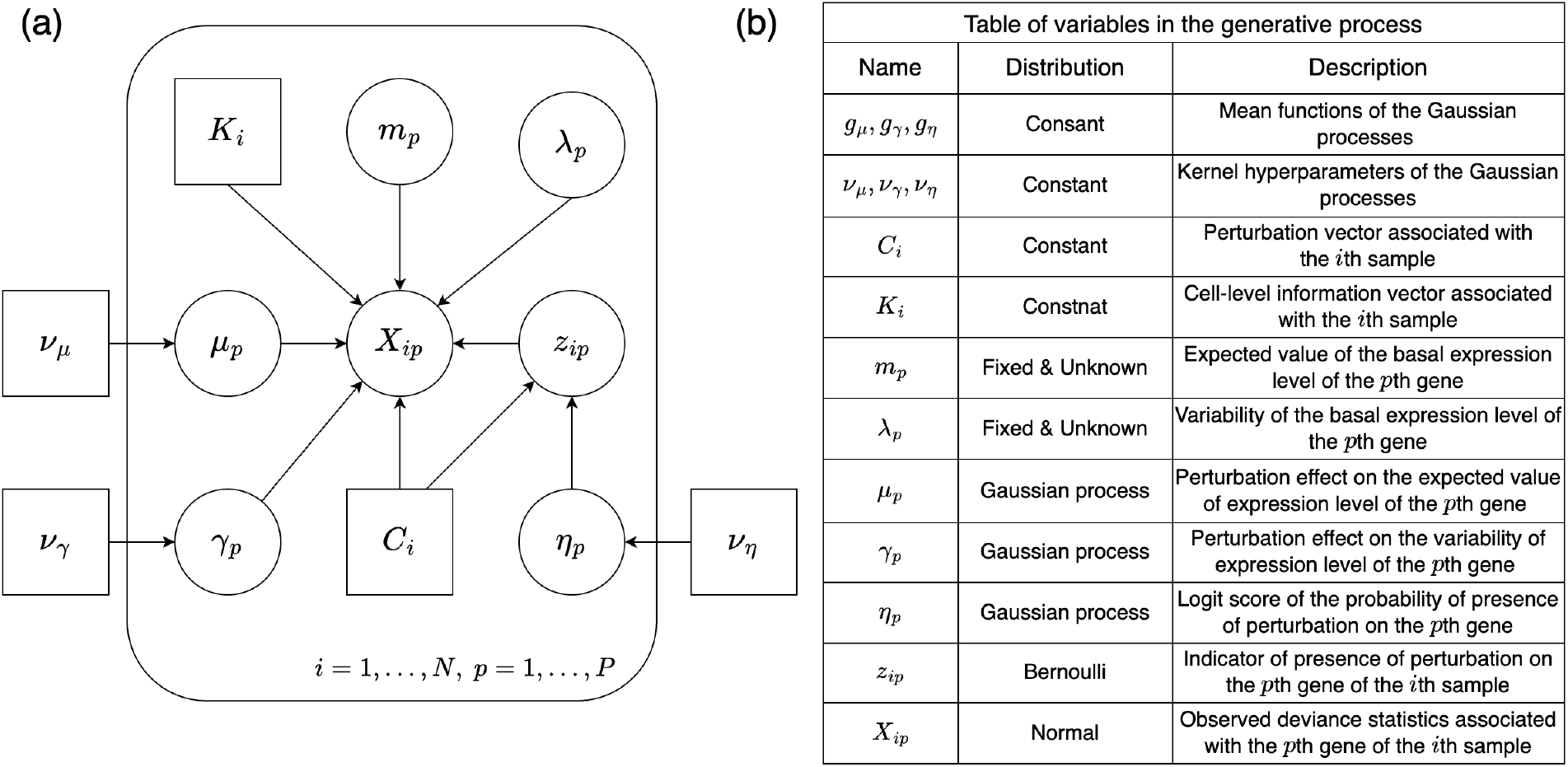
(a): A graphical representation of the proposed deviance-based Normal model. (b): Table of parameters used in the proposed model.

##### S1.2 Zero-inflated Gamma-Poisson model

Here we give details of the zero-inflated Gamma-Poisson model variant:

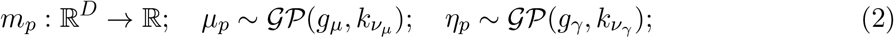

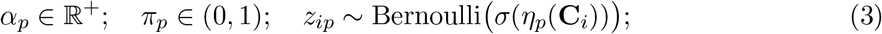

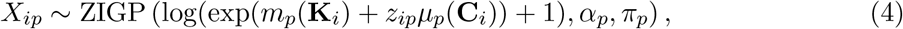

where *m*_*p*_, *µ*_*p*_, *η*_*p*_, *π*_*p*_, *z*_*ip*_ have the same interpretation as in the Zero-inflated Poisson model seen previously, *α*_*p*_ is the dispersion parameter associated with the *p*th gene, and ZIGP(*µ, α, π*) is a Zero-inflated Gamma-Poisson distribution with p.m.f.

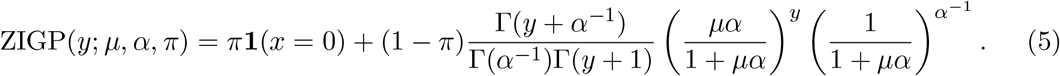

The ELBO estimate and variational posterior inference of the ZIGP model above can be carried out using the same procedure as the Zero-inflated Poisson model.

**Supplementary Figure 2:**
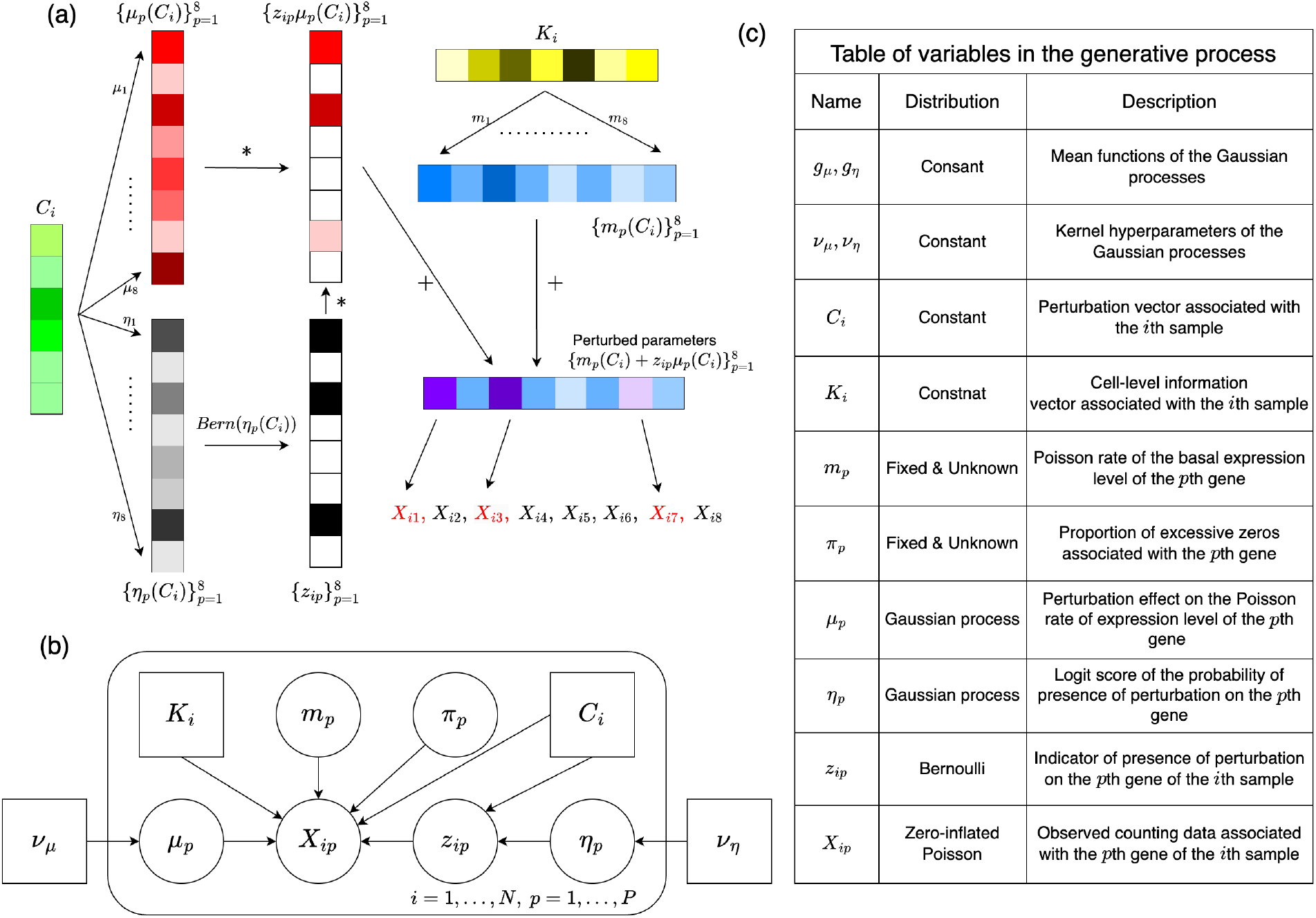
(a): A schematic illustration of the generative process of the raw counting data vector **X**_*i*_ associated with the *i*th sample. Here we assume *L* = 6, *D* = 7, *P* = 8. * denotes elementwise product. The perturbed gene expressions {*X*_*i*1_, *X*_*i*3_, *X*_*i*7_} are highlighted in red. (b): A graphical illustration of the proposed Zero-inflated Poisson model. (c): Table of variables used in the proposed Zero-inflated Poisson model

#### S2 Connection to existing methods

In this section, we discuss the connection between our proposed approach and some existing works.

##### S2.1 Guided sparse factor analysis (GSFA)

Our deviance-based additive model is motivated by guided sparse factor analysis (GSFA) ^11^. However, our methods differ from GSFA in the several ways: Firstly, our approach aims to directly give sparse estimates of gene-level perturbation effects, while GSFA focus on constructing a latent factor model that encodes the perturbation and co-regulated genes as sparse factor and loading vectors. Secondly, our modelling framework allows us to assess the inclusion/exclusion of gene-level perturbation effects on in an intuitive way using posterior inclusion probabilities of the binary toggles *z*_*ip*_s. In contrast, GSFA has to resort to multiple testing correction procedures such as LSFR ^8^. In addition, our proposed method estimates basal and perturbation effects simultaneously using flexible regression models, while GSFA estimates them in a sequential fashion using linear regression and factor analysis models. Hence we also expect our approach to be able to capture finer details of the dataset, and reveal more insight of the complex biological process of single-cell perturbations.

##### S2.2 Compositional perturbation autoencoder (CPA)

The Compositional Perturbation Autoencoder ^3^ (CPA) aims to predict counterfactual distributions of gene expression of a given cell under a generic unobserved perturbation using a variational autoencoder and additive latent embedding of the cell and perturbation states. In comparison with CPA, our approach is less flexible in term of perturbation effect estimation. However, our approaches focus more on disentangling non-sparse basal states and sparse gene-level perturbation effects for each perturbed sample, hence offering better interpretability than the black box decoder and non-sparse latent representations of cell and perturbation states in CPA. Both our proposed methods and CPA can utilize cell-level information to inform the basal states of the samples. However, CPA assumes categorical cell-level information and continuous gene expression responses, while our methods can handle both continuous and discrete cell-level information and gene expression responses.

##### S2.3 SAMS-VAE

With SAMS-VAE, Bereket and Karaletsos ^2^ introduce a Sparse additive mechanism shift variational autoencoder (SAMS-VAE) to characterise perturbation effects as sparse latent representations. In SAMS-VAE, the latent representation of a perturbed expression vector is obtained by adding a sparse representation of the perturbation to a dense perturbation-independent basal state, and the decoder is trained to reconstruct the perturbed expression vectors from latent representations. Compared with SAMS-VAE, our approach directly model and estimate sparse perturbation effects instead of a sparse latent representation of it. In addition, SAMS-VAE is not able to incorporate additional cell-level information into the model such as batch information or cell type (i.e. the latent basal states in SAMS-VAE can not be informed by cell-level information **K**), and can only handle binary perturbation and counting data. Our methods are not restricted by these constraints.

**Supplementary Figure 3:**
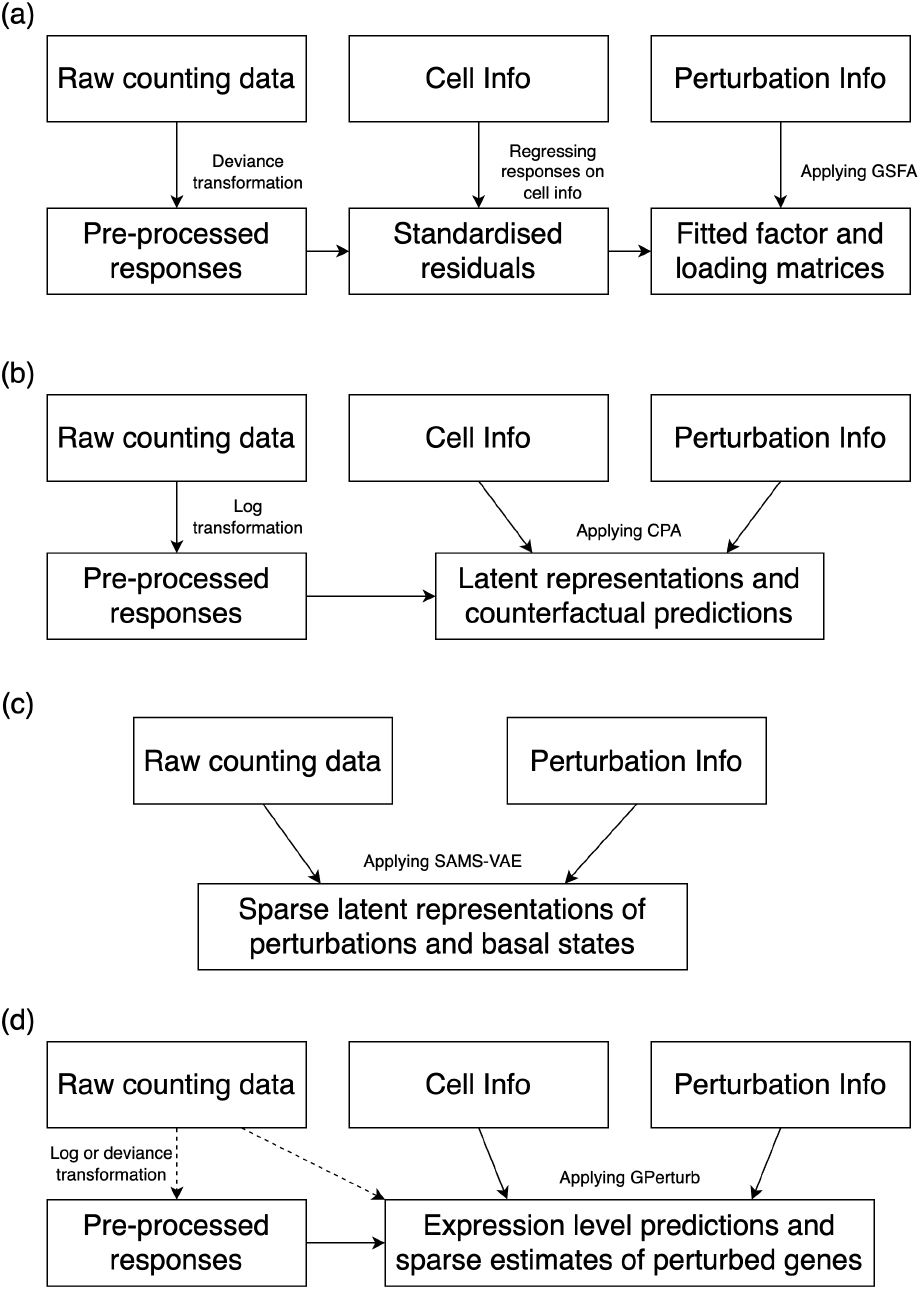
Schematic illustration of training pipelines (a) GSFA (b) CPA (c) SAMS-VAE and (d) GPerturb

**Supplementary Figure 4:**
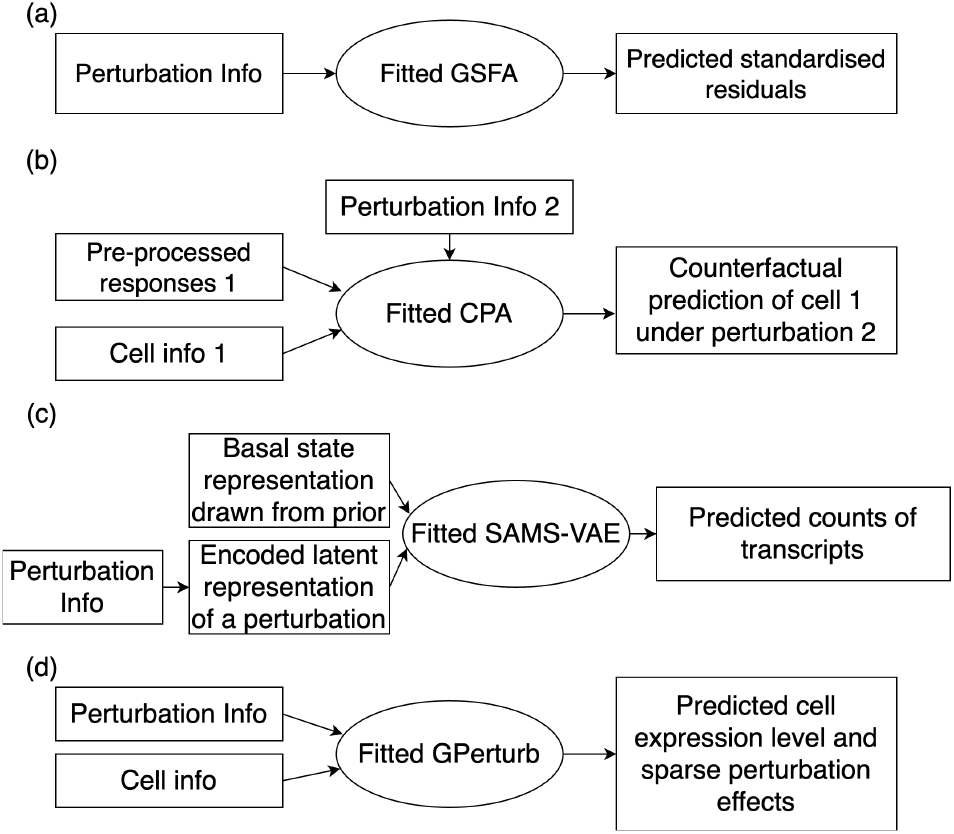
Schematic illustration of inference/prediction pipelines (a) GSFA (b) CPA (c) SAMS-VAE and (d) GPerturb.

##### S2.4 GEARS

In GEARS, Roohani et al. ^6^ proposed a graph-enhanced gene activation and repression simulator (GEARS), a computational model that predicts the gene expression outcomes of combinatorially perturbing a set of one or more genes. GEARS uses a prior knowledge graph of gene–gene relationships to inform the prediction, allowing it to simulate the outcomes of perturbing unseen single genes or combinations of genes. Comparing with our approach, predictions given by GEARS lack interpretability, and the model is only applicable to datasets with single- or multigene perturbations. In contrast, our approach is able to give probabilistic estimates of subsets of genes targeted by the perturbation in addition to the perturbed expression outcomes, and is applicable to both genetic and chemical perturbations.

##### S2.5 Training and inference pipelines of different methods

Figures 3 and 4 shows the setups for training and prediction respectively for GSFA, CPA, SAMS-VAE and GPerturb. For training, as discussed earlier, GSFA is characterised by the requirements to pre-process and standardise the input data for modelling. CPA also functions of transformed input data and maps to perturbations defined in latent representations. SAMS-VAE accepts count data and also uses latent perturbation spaces.

GPerturb can use either transformed or count data and models perturbed and normal expression levels directly. This means that the trained GPerturb model can be readily inverted for prediction and inference tasks and output can be reported on the original scale of the inputs including at count level.

In contrast, the use of standardised and processed inputs in GSFA limits interpretation of the output of the trained model. While it is necessary to have expression input from a perturbation to use as a reference for counterfactual prediction of expression under an alternate perturbation for CPA. With SAMS-VAE, training allows it learns to map perturbed expression profiles into an estimate of the (unperturbed) basal latent state via an encoder. However, at prediction time, it is not possible to make predictions unless some input data is available for the encoder and hence the cells.

These methods are designed for different purposes and hence require different inputs. Compared with existing state-of-the-art methods, the training and inference procedures of our proposed GPerturb is more intuitive.

#### S3 Simulation Experiments

Here we demonstrate that our proposed methods can learn sparse combinatorial perturbation effects using two simulated datasets.

##### S3.1 Gaussian GPerturb

We first demonstrate the efficacy of our proposed models using a simulated dataset consisting of continuous expression levels. Let the dimension of gene expression vector *P* = 6000, dimension of perturbation vector *L* = 15, dimension of cell-level information *D* = 4, number of cells *N* = 4000. The simulated data is generated as follows: Let **K** be a *N* × *D* matrix such that the entries in the first two columns of *K* are samples drawn from i.i.d. 𝒩 (0, 1), and the last two entries in each row of **K** are the one-hot encoding of a categorical sample drawn form 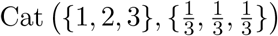. We generate **K** in this fashion since cell-level information can either be continuous or categorical in real world applications. Let **C** be a *N* × *L* binary matrix with each entry being a sample from Bernoulli(0.2). Let *λ*_*p*_ be samples from i.i.d. 𝒩 (0, 1) for *p* = 1,…, *P*. Let **H**_1_ ∈ ℝ^*K*×*P*^, **H**_2_, **H**_3_, **H**_4_ ∈ ℝ^*D*×*P*^ be matrices whose entries are drawn from i.i.d. N (0, 1). For each *i* = 1,…, *N* and *p* = 1,…, *P*, we set *m*_*p*_(*K*_*i*_) = (**KH**_1_)_*ip*_ (i.e. the corresponding entry in the matrix 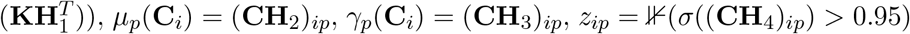), *µ*_*p*_(**C**_*i*_) = (**CH**_2_)_*ip*_, *γ*_*p*_(**C**_*i*_) = (**CH**_3_)_*ip*_, *z*_*ip*_ = ⊮(*σ*((**CH**_4_)_*ip*_) *>* 0.95), and **X**_*ip*_ ∼ 𝒩 (*m*_*p*_(**K**_*i*_) + *z*_*ip*_*µ*_*p*_(**C**_*i*_), log(exp(*λ*_*p*_ + *z*_*ip*_*γ*_*p*_(**C**_*i*_)) + 1). In the simulated dataset, roughly 5% of the *z*_*ip*_s are ones. In other words, roughly 5% of the sample-gene pairs in this simulated dataset are perturbed.

In order to assess the generalization performance of our proposed model on unobserved perturbation patterns, we split the simulated dataset into a training set and a test set in the following way: Let *E* be the set consisting of 40 unique perturbation vectors uniformly and randomly selected from the rows of *C*. Let *F* = {*i* ∈ {1,…, *N*}|**C**_*i*_ ∈ *E*} be the index set of all samples whose associated perturbation vector is in *E*. Let {**X**_−*F*_, **C**_−*F*_, **K**_−*F*_} be the training set (i.e. samples whose indices are not in *F*), and {**X**_*F*_, **C**_*F*_, **K**_*F*_} be the test set. By doing so, we ensure that the perturbation vectors in the test set are unseen in the model’s training process. The size of test set is roughly 15% of the full synthetic dataset. Performance of the fitted model on test set is reported in Supplementary Fig 5. We see the model estimates both basal states and perturbation effects associated with unseen perturbation vectors accurately. In addition, the model also accurately identifies whether or not a sample-gene pair is perturbed by an unseen perturbation vectors (AUC = 0.966).

##### S3.2 Zero-inflated Poisson GPerturb

In this section, we demonstrate the efficacy of the proposed zero-inflated Poisson model using a simulated example. The dimension of the dataset and the synthetic data **C, K, H**_1_, **H**_2_, **H**_3_ are chosen and generated in the same fashion as in the previous example. Let *π*_*p*_ be samples from i.i.d. Beta(2, 10) for *p* = 1,…, *P*. In this example, we set *m*_*p*_(**K**_*i*_) = 5(**KH**_1_)_*ip*_ + 50, *µ*_*p*_(**C**_*i*_) = 5(**CH**_2_)_*ip*_, *z*_*ip*_ = ⊮(*σ*((**CH**_3_)_*ip*_) *>* 0.95) and **X**_*ip*_ ∼ ZIP(log(exp(*m*_*p*_(**K**_*i*_) + *z*_*ip*_*µ*_*p*_(**C**_*i*_)) + 1), *π*_*p*_). The synthetic basal rates and perturbation effects are scaled to mimic the size of counts in real datasets. The training and test set are split in the same way as in the previous example. Performance of the fitted model on test set is reported in Supplementary Fig 6. We see it predicts unseen basal states and perturbation patterns accurately, and is able to correctly identify the sparse perturbation effects of unseen perturbation vectors (AUC = 0.901).

**Supplementary Figure 5:**
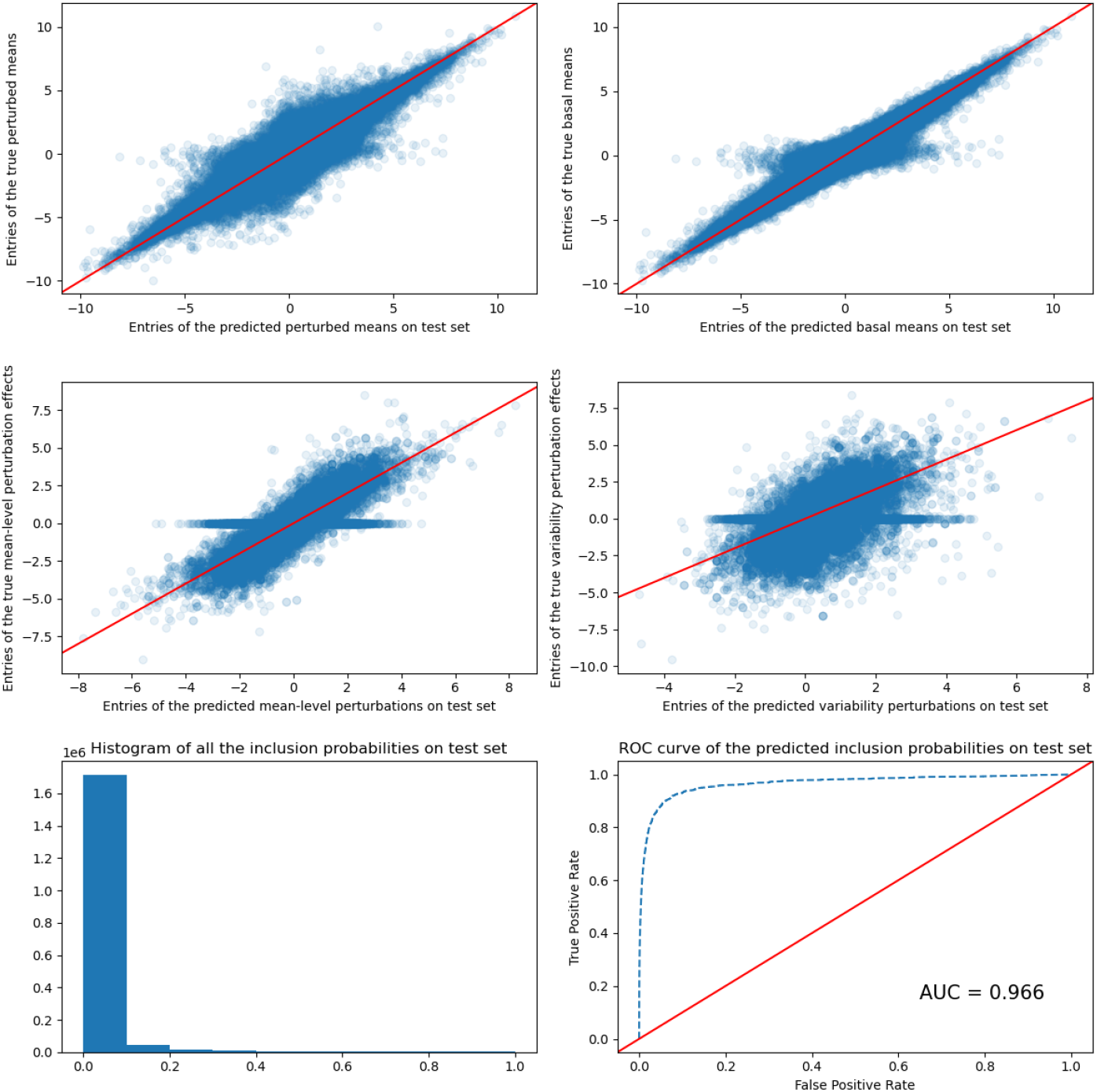
Estimated perturbation effects and inclusion probabilities on the test set consisting of unseen perturbation patterns. Top: Scatter plots of the estimated perturbed mean v.s. the true perturbed mean and estimated basal mean v.s. true basal mean on test set. Mid: Scatter plots of estimated mean-level and variability perturbation vs the truth (i.e. *z*_*ip*_*µ*_*p*_(**C**_*i*_) and *z*_*ip*_*γ*_*p*_(**C**_*i*_) respectively) on test set. Bottom left: Histogram of estimated inclusion probabilities on test set. Bottom right: ROC curve and AUC indicating how well the estimated inclusion probabilities predict the binary toggle *z*_*ip*_s on test set.

**Supplementary Figure 6:**
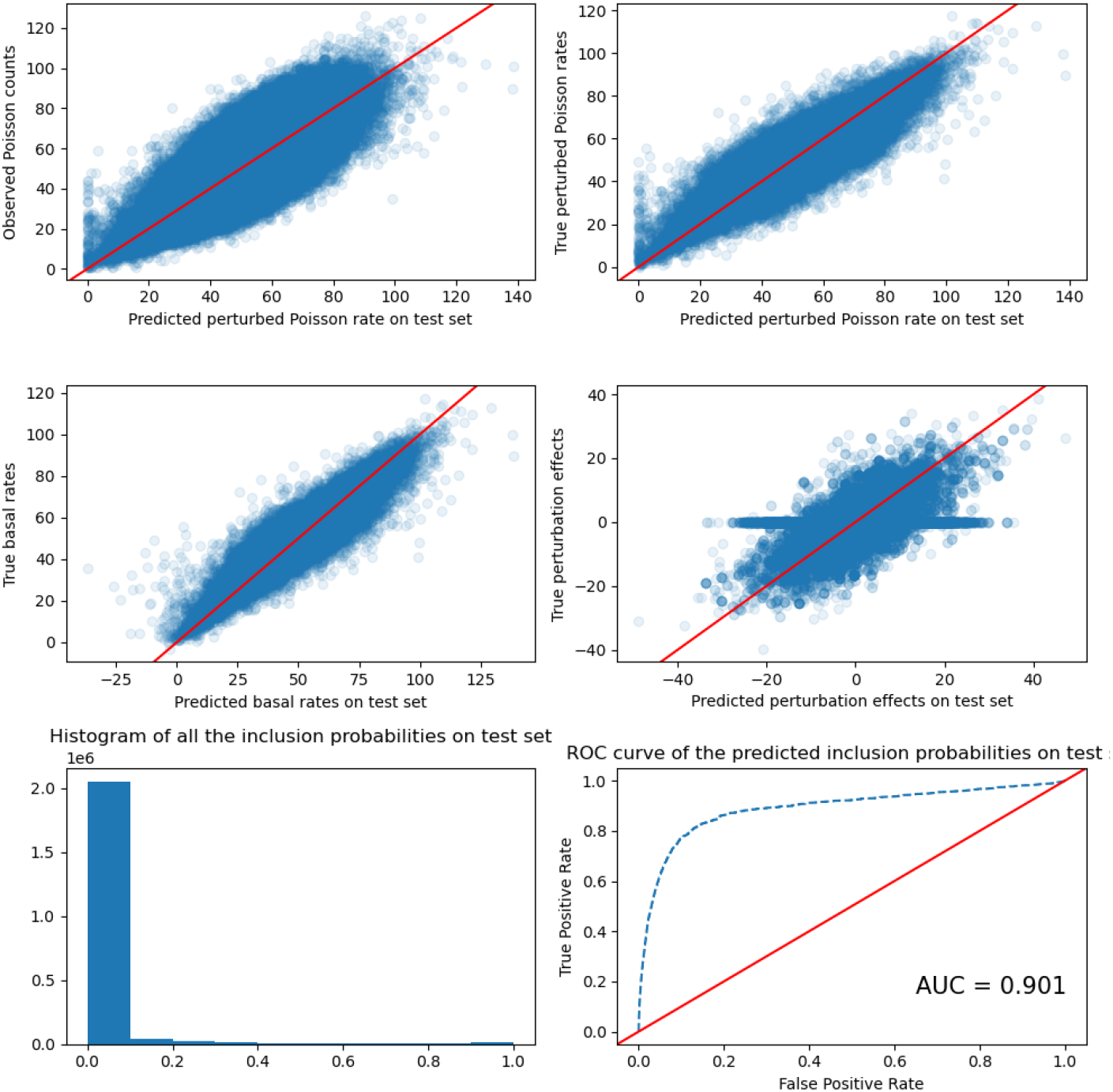
Estimated perturbation effects and inclusion probabilities on the test set consisting of unseen perturbation patterns. Top: Scatter plots of the estimated perturbed Poisson mean v.s. the true observations and the estimated perturbed Poisson mean v.s. true Poisson means of the non-zero entries in test set. Mid: Scatter plots of the estimated basal rate v.s. true basal rate and estimated mean-level perturbation vs the truth perturbation effects of the non-zero entries in test set. Bottom left: Histogram of estimated inclusion probabilities of the non-zero entries in test set. Bottom right: ROC curve and AUC indicating how well the estimated inclusion probabilities predict the binary toggle *z*_*ip*_s of the non-zero entries in test set.

#### S4 Further Results

##### S4.1 Zero-inflated Gamma-Poisson GPerturb

In this section, we provide further details of the analysis of datasets discussed in the main text using the Zero-inflated Gamma Poisson variant of GPerturb. Supplementary Figures 7, 8, 9 and 10 show comparative performance between this and the Zero-inflated Poisson model used in the main text. Supplementary Figure 11 shows the empirical distribution of the estimated dispersion parameters *α*_*p*_) across all genes. The results indicate that due to the low dispersion, the simpler Zero-inflated Poisson model used in the main text is sufficient for these datasets.

**Supplementary Figure 7:**
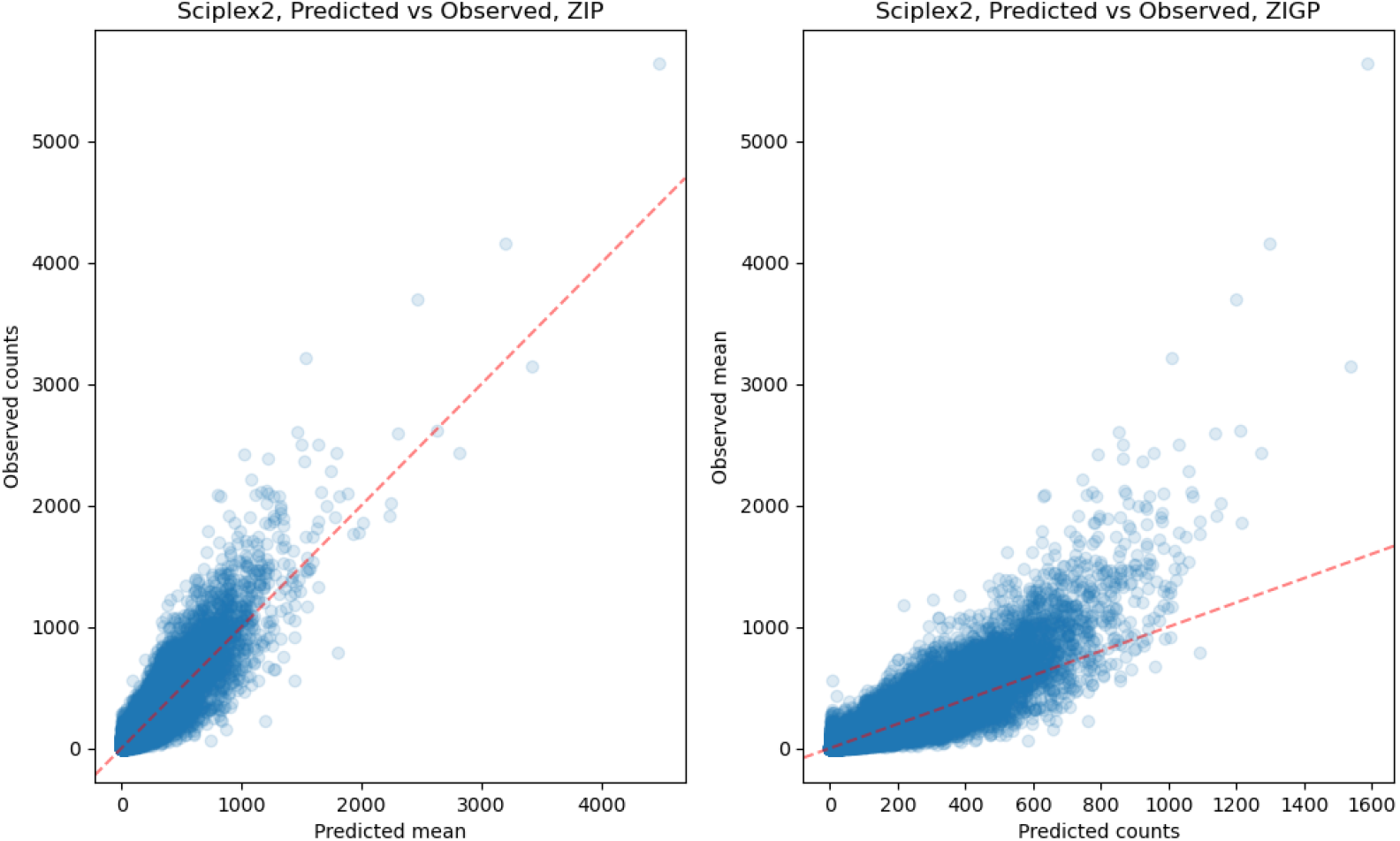
Non-zero observed counts for each cell-gene pair vs corresponding estimated mean for each cell-gene pair given by ZIP and ZIGP GPerturb on test set.

**Supplementary Figure 8:**
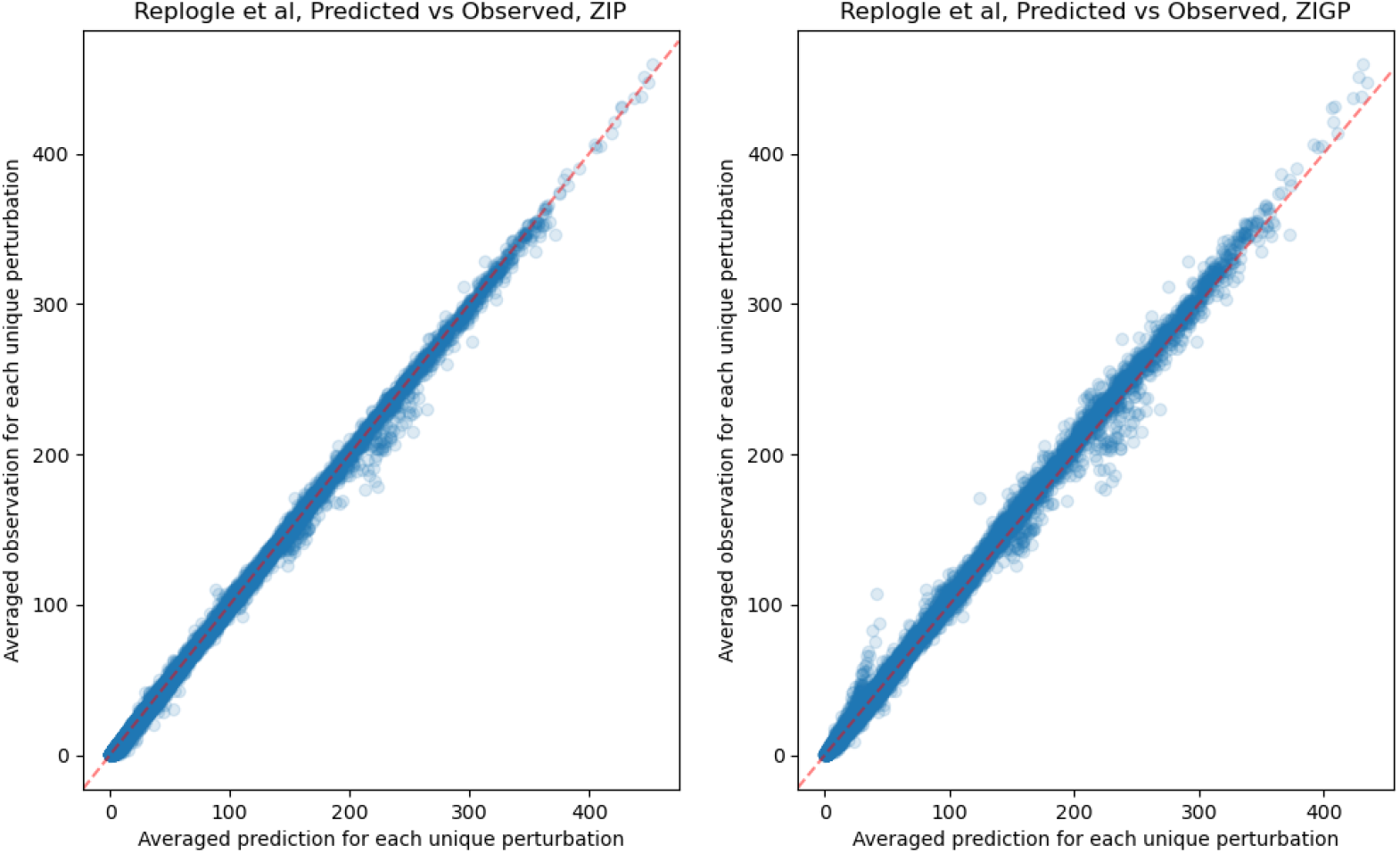
Non-zero observed counts for each cell-gene pair vs corresponding estimated mean for each cell-gene pair given by ZIP and ZIGP GPerturb on test set.

**Supplementary Figure 9:**
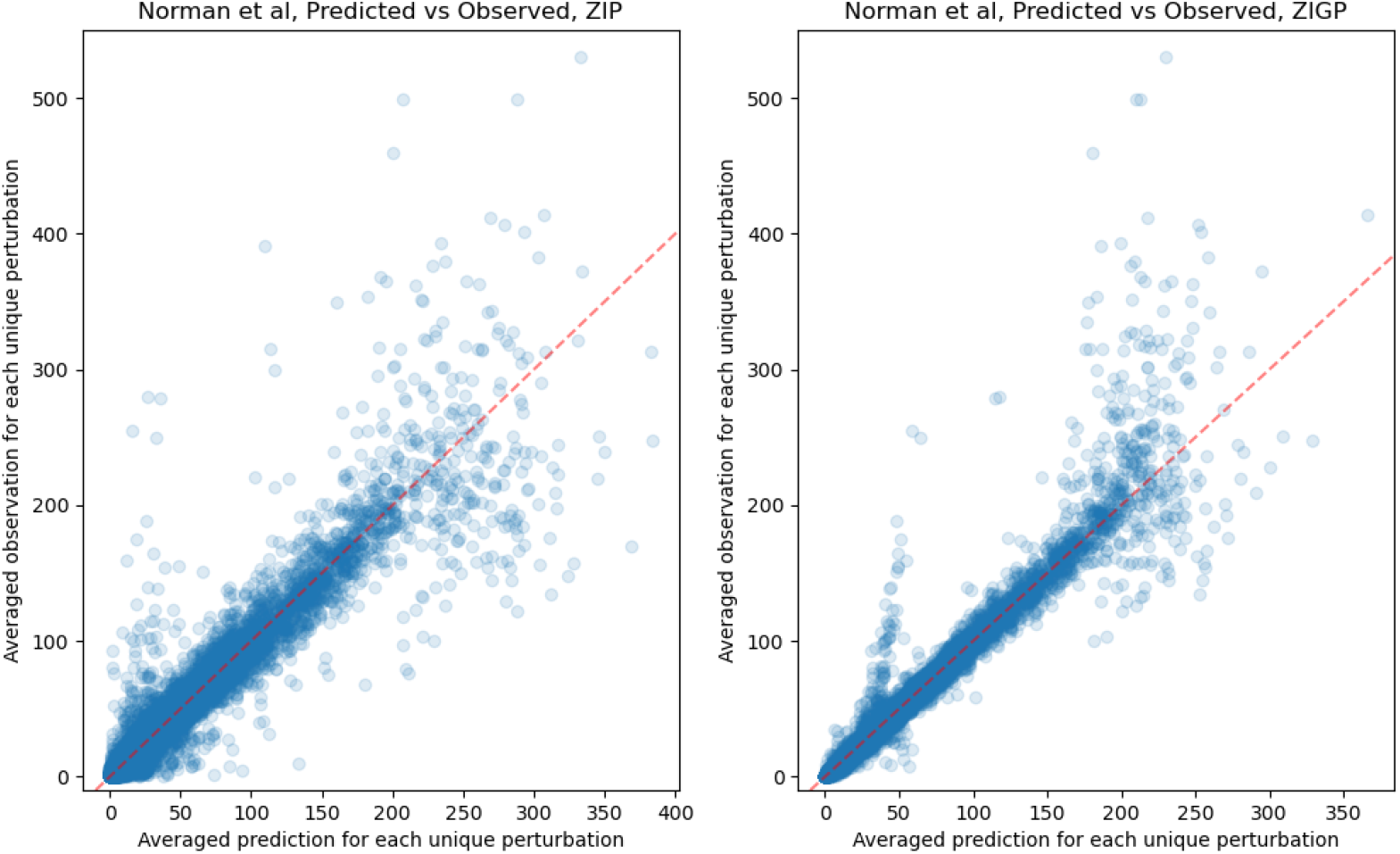
Non-zero observed counts for each cell-gene pair vs corresponding estimated mean for each cell-gene pair given by ZIP and ZIGP GPerturb on test set.

**Supplementary Figure 10:**
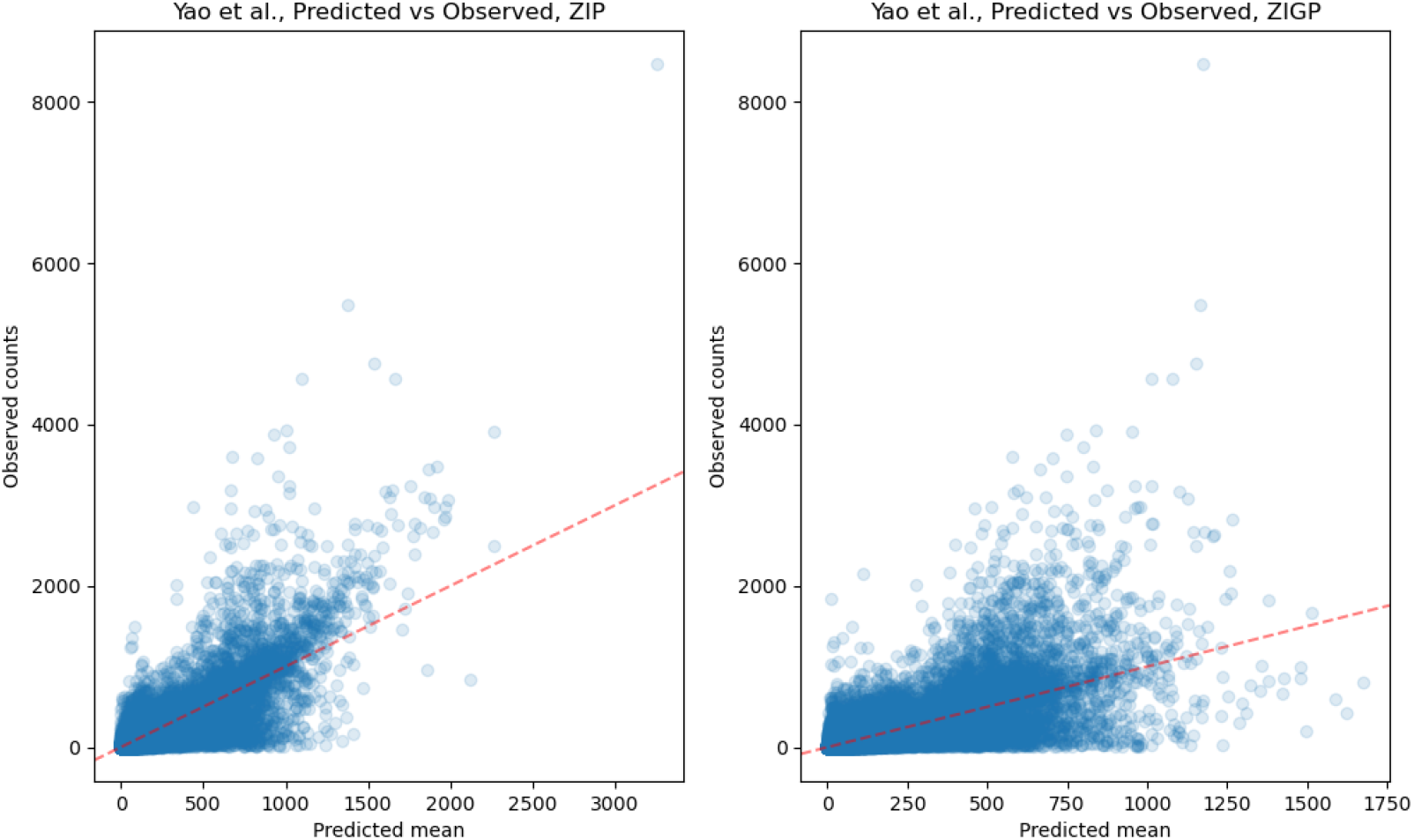
Non-zero observed counts for each cell-gene pair vs corresponding estimated mean for each cell-gene pair given by ZIP and ZIGP GPerturb on test set.

##### S4.2 Computation cost

In this section we compare the computation cost of GPerturb with existing methods. Since distinct modeling architecture are used in different methods, we choose to report the wall clock running time as a fair benchmark for computation cost (Supplementary Table 1). We see that the computation cost of GPerturb in term of running time is on a similar level to existing methods. All experiments are conducted on our machine with an AMD Ryzen 7 2700 CPU and an NVidia RTX 2060 GPU.

**Supplementary Figure 11:**
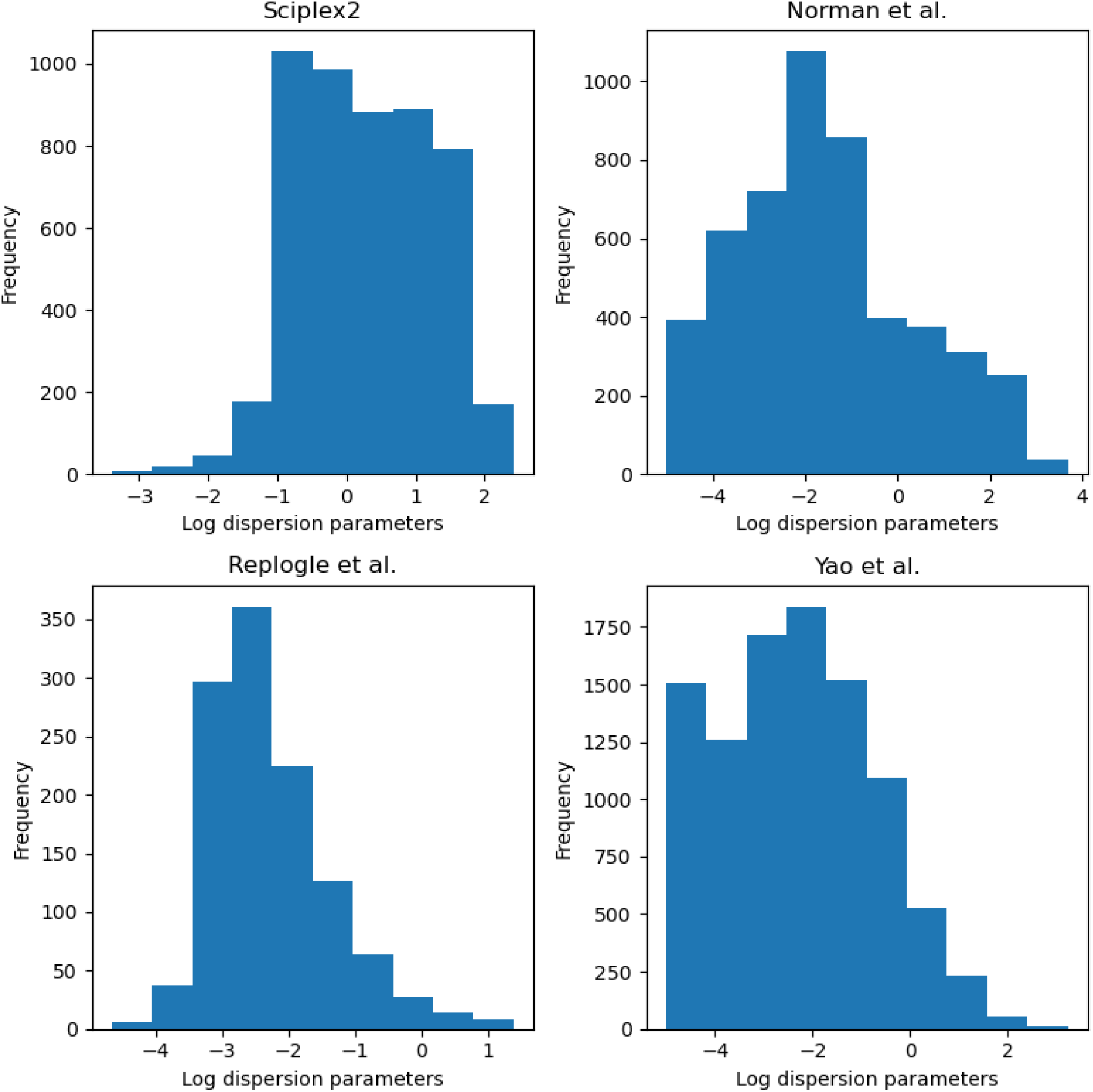
Histograms of estimated log dispersion parameters for each of the datasets we studied in the paper.

##### S4.3 GPerturb’s Bayesian probabilistic modeling framework

The probabilistic Bayesian modeling framework improves interpretability and uncertainty quantification of the results. Existing methods such as SAMS-VAE^2^ and CPA^3^ focus on predicting counterfactual perturbed gene expressions using latent embeddings which lack natural biological meaning. GEARS^6^ is able to predict the effect of a given perturbation on individual genes, and give ad-hoc uncertainty quantification. However, the lack of sparsity in GEARS predictions makes it difficult for users to identify genes responsive to different perturbations in a principled way. Our probabilistic Bayesian modeling framework address these issues. For example, GPertrub provides easy-to-interpret uncertainty quantification (credible intervals) of the predicted perturbation effects (See Fig 2c). In addition, GPerturb is able to identify and select responsive genes to different perturbations in a straightforward and interpretable fashion using the estimated posterior inclusion probabilities (PIP) for each gene-perturbation pair (i.e. the probability of a gene being responsive to a perturbation) thanks to the probabilistic modeling framework (See Methods). To further illustrate this point, we report the estimated perturbational effects of exosome-related perturbations on the top-25 most differentially expressed genes identified by GPerturb in Replogle et al ^5^ data at different PIP inclusion thresholds (Supplementary Fig 12. We see that by increasing the PIP inclusion threshold from 0.5 to 0.99, users can easily filter out the subset of gene-perturbation pairs that are most confidently selected by the model based on this interpretable threshold. This means users can use GPerturb to directly handle queries such as “how likely is a gene responsive to a given perturbation” or “what subset of genes are most likely to be responsive to a given perturbation” without any ad-hoc post-processing steps used in e.g. GEARS and CPA.

**Supplementary Table 1:**
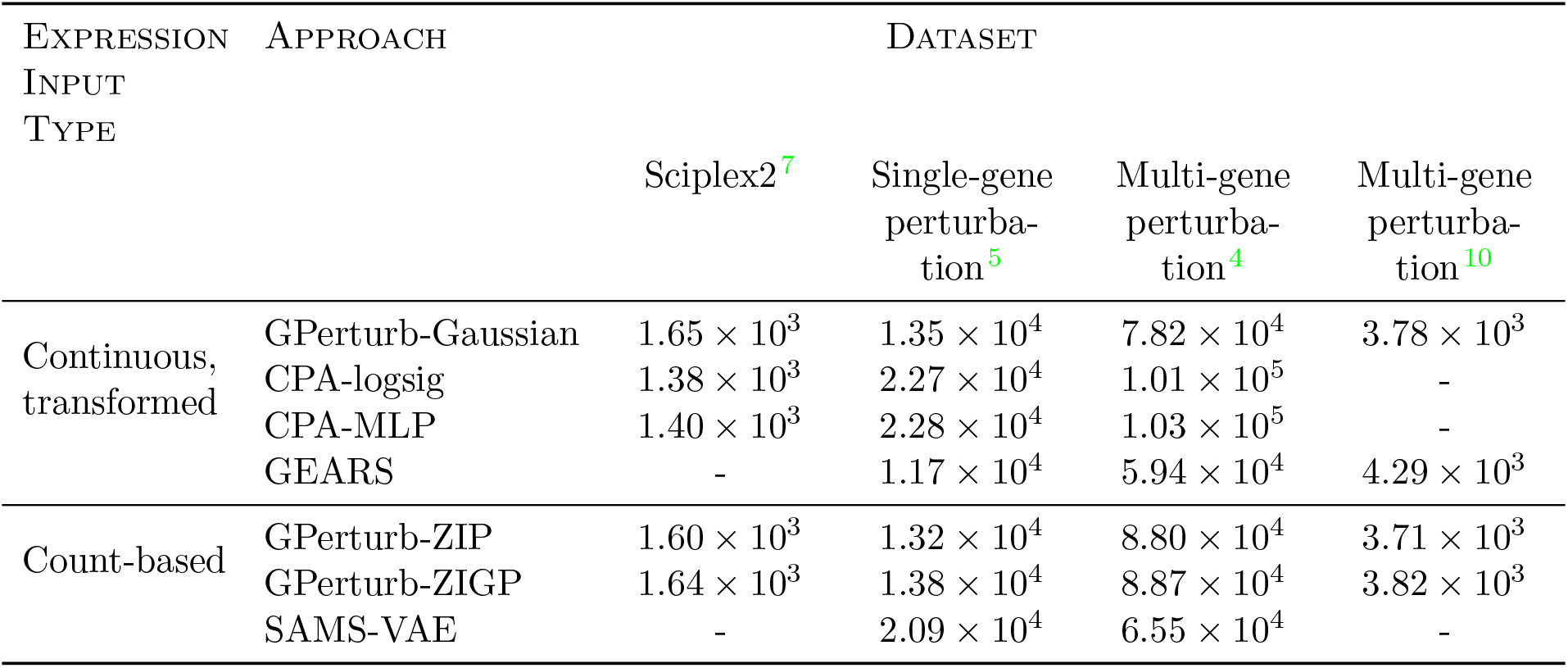
Comparison of wall clock running time. Values show the averaged wall clock running time in seconds for each method and each data set over 3 repetitions.

#### S5 Additional Experiments

The following describes additional experiments not described in the main manuscript and particularly focuses on comparison to GSFA.

##### S5.1 LUHMES neural progenitor cell CROP-seq study

We applied Gaussian GPerturb to the LUHMES neural progenitor cell CROP-seq dataset (GSE142078) studied in Zhou et al. ^11^, and compare its performance to GSFA. This study targets 14 neurodevelopmental genes, including 13 autism risk genes, in LUHMES human neural progenitor cells. The raw data is preprocessed using the identical procedure described in Zhou et al. ^11^. The resulting dataset **X** ∈ ℝ^*N* ×*P*^ consists of *N* = 8708 samples and *P* = 6000 selected genes. For *i* = 1,…, *N*, the perturbations **C**_*i*_ ∈ {0, 1}^*L*^ are encoded as one-hot vectors of length *L* = 14, each corresponds to one of the 14 targeted neurodevelopmental genes (i.e. 14 distinct perturbations). The cell information **K**_*i*_ ∈ ℝ^*D*^ is a real vector of length *D* = 4 (`lib_size`: number of total UMI counts, `n_features`: number of genes with non-zero UMI readings, `mt_percent`: percentage of mitochondrial gene expression and `batch`: batch ID). In addition to the one-hot perturbations, the dataset also consists of negative control gRNAs whose perturbations are encoded as **C**_*i*_ = **0**. Recall that our choices of generative process and variational family ensure that the negative controls with **C**_*i*_ = **0** have zero perturbation effects. By doing so, users can view the negative controls as the baseline level, and the perturbation effects associated with the non-zero **C**_*i*_s as the perturbation strength relative to the negative controls. In Zhou et al. ^11^, the authors first remove cell level information from the transformed responses by regressing **X** on **K** using linear regression, then apply GSFA to the corresponding standardized residual matrix. In contrast, our proposed method disentangles and estimates cell-level and perturbation-induced variations simultaneously, and does not require any standardisation. For our proposed method, we randomly select 20% of the dataset as the test set, and use the rest to train GPerturb. The default priors discussed in Methods are used here. For GSFA, the results are obtained based on the recommended settings given in Zhou et al. ^11^.

**Supplementary Figure 12:**
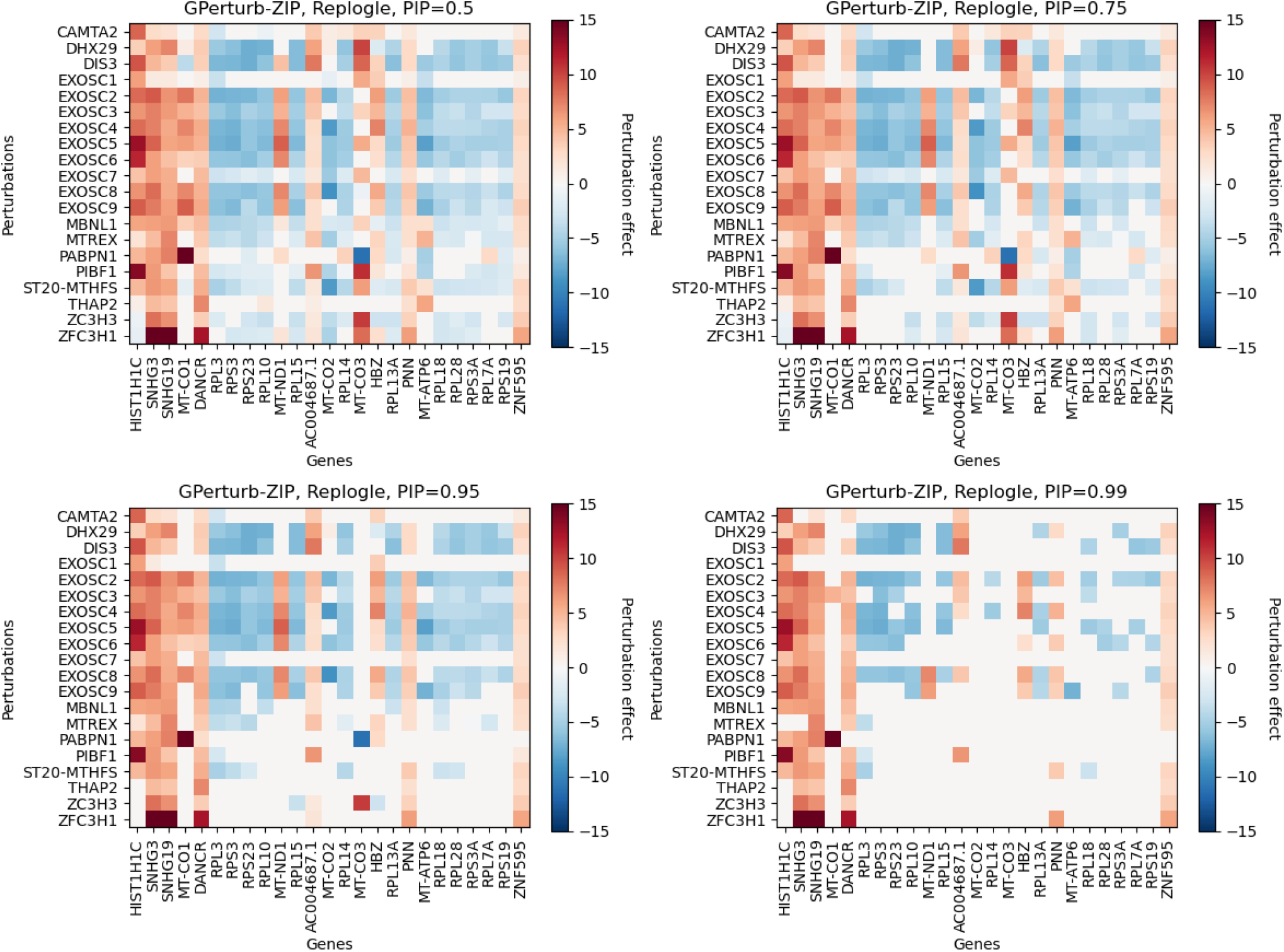
Estimated perturbation effects associated with exosome-related perturbations in Replogle et al ^5^ on a subset of differentially expressed genes identified by the model under different posterior inclusion probability thresholds. Note that as the threshold increases, less gene-perturbation pairs are deemed to be responsive.

In this example, we are interested in comparing the perturbation-induced variations captured by the two methods. Compared with our approach, GSFA requires additional dataset-dependent pre-processing and standardisation steps, which could potentially restrict its interpretability and generalisation power. (See Supplementary Fig 3, 4 for an illustration of the different training and inference pipelines of the methods discussed in this paper.) To make the results of the two methods comparable, we apply the following transformations to the fitted Gaussian GPerturb, mimicking the pre-processing steps in GSFA^11^: Let *N* ^′^ = 0.2⌊*N* ⌋ be the number of samples in the test set. For each sample *i* = 1,…, *N* ^′^ in the test set, we first let 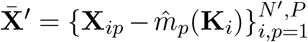 to be the residual matrix (i.e. subtract the estimated cell-level variation 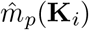 from the observed response **X**_*ip*_), then we standardise the columns of 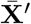,and apply the same standardisation to the estimated mean perturbation effect matrix 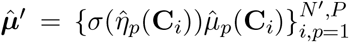. Let 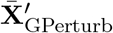 and 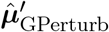 be the standardised residual matrix and estimated mean perturbation effect matrix given by GPerturb. Let 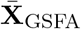 and 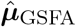 be the corresponding standardized residual matrix and estimated gene-level perturbation effect given by GSFA on the full dataset (GSFA uses the *entire* dataset to estimate the factor/loading matrices). In other words, one can view the standardised residuals 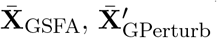 as transformed, noisy observations associated with perturbation treatments, and 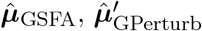 as the estimated mean perturbation effects. We stress that the fitted Gaussian GPerturb is more interpretable on the original scale, and transformations applied to it may affect its prediction power. The purpose of the transformations above is only to map the fitted results onto a scale comparable with GSFA, and is not necessary in practice.

In this comparison, we focus on predictive performance on cell-gene pairs whose expression levels are more likely to be perturbed. To do so, we first select entries in 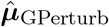 whose corresponding estimated posterior inclusion probability 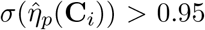,then report the scatter plot of the selected entries in perturbation effects 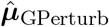 verses the same set of selected entries in standardised residuals 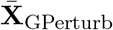 in Supplementary Fig 13. We also report the scatter plot of the same set of selected entries in 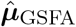 verses 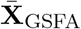. We find that under this comparison framework in favour of GSFA, the post-processed GPerturb achieves similar performance to GSFA on the set of selected entries (Pearson correlation *r*_GPerturb_ = 0.248, *r*_GSFA_ = 0.182) in term of Pearson correlation between the transformed observations 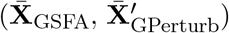 and predictions 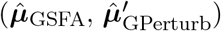,indicating that GPerturb is able to capture details of perturbation effects. The fitted verses observed scatter plot on test set is reported in Supplementary Fig 13.

We remind the reader that this comparison is qualitative and not rigorous as the two models are estimated using different pre-processing steps and objectives.

**Supplementary Figure 13:**
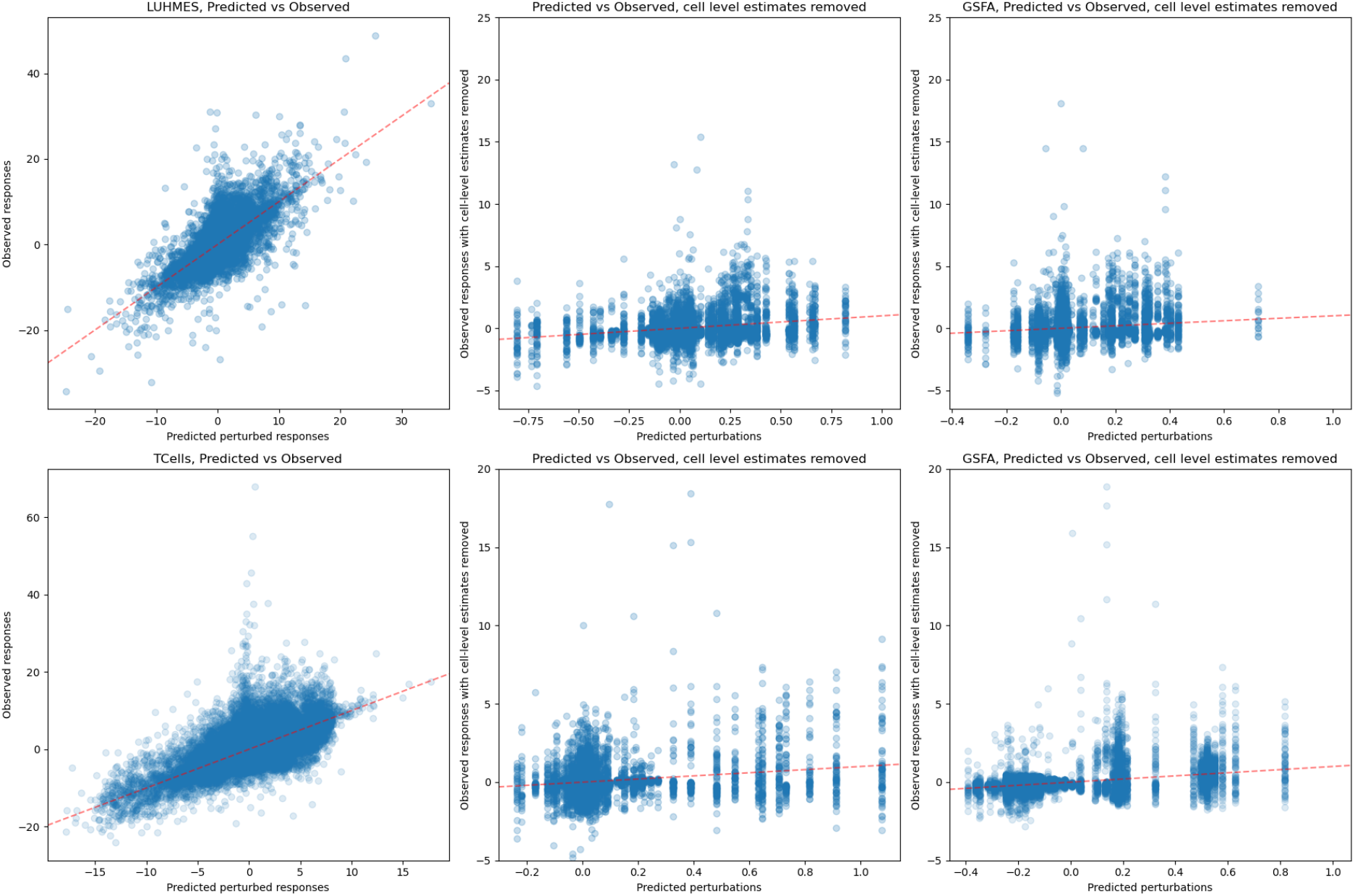
**Top**: Fitted values of Gaussian GPerturb and GSFA on the LUHMES dataset. **Top left**: Predicted expression level given by Gaussian GPerturb vs observed expression level in the LUHMES test set. **Top middle**: Standardised perturbation effects 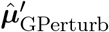 verses standardised residuals 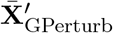 of the selected “active” gene-sample pairs whose corresponding posterior inclusion probability 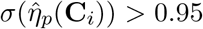.**Top right**: Standardised perturbation effects 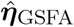 verses standardised residuals 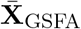 of the same set of selected “active” gene-sample pairs. **Bottom**: Fitted values of Gaussian GPerturb and GSFA on the human T Cells dataset. Figures have the same interpretation as in the top row.

We also report heat map of the estimated perturbation effects associated with each of the 14 unique perturbations 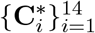 given by Gaussian GPerturb in Supplementary Fig 14. Here a perturbation-gene pair is considered “active” only if the corresponding estimated posterior inclusion probability 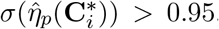,and a gene *p* is included only if at least one of the 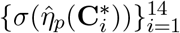 is greater than 0.95. The resulting heat map provides an intuitive visualization of perturbation effects on different genes. In addition to the genes associated with large estimated perturbation effects, we also report a similar heat map on a collection of marker genes studied in Zhou et al. ^11^ in Supplementary Fig 15. Comparing with Zhou et al. ^11^, we found that GPerturb identifies less active perturbation-gene pairs than GSFA (See also Supplementary Fig 19). To further investigate this difference, we select three marker genes (NES, CRABP2, HDAC2) from different pathway groups, and compare the observed expression level of these genes with the GPerturb prediction under three different perturbations (CHD2, PTEN, SETD5) (Supplementary Fig 16). For each perturbation-gene pair, we use a two-sample *t*-test to test if the mean of the expression levels under the perturbation is different from the baseline (i.e. mean expression level under the non-targeting perturbation). The *p*-values are then corrected using Benjamini–Hochberg procedure (B-H) ^1^, and a gene-perturbation pair is considered active if the corrected *p*-value is less than 0.05. We then compare the subset of active perturbation-gene pairs identified by B-H with the ones identified by GPerturb and GSFA. From Supplementary Fig 16 we see GPerturb’s predicted expression levels agree with the observed values, and the subset of pairs identified by GPerturb agrees better with the one identified by B-H, suggesting that GSFA may consist of more false positives.

**Supplementary Figure 14:**
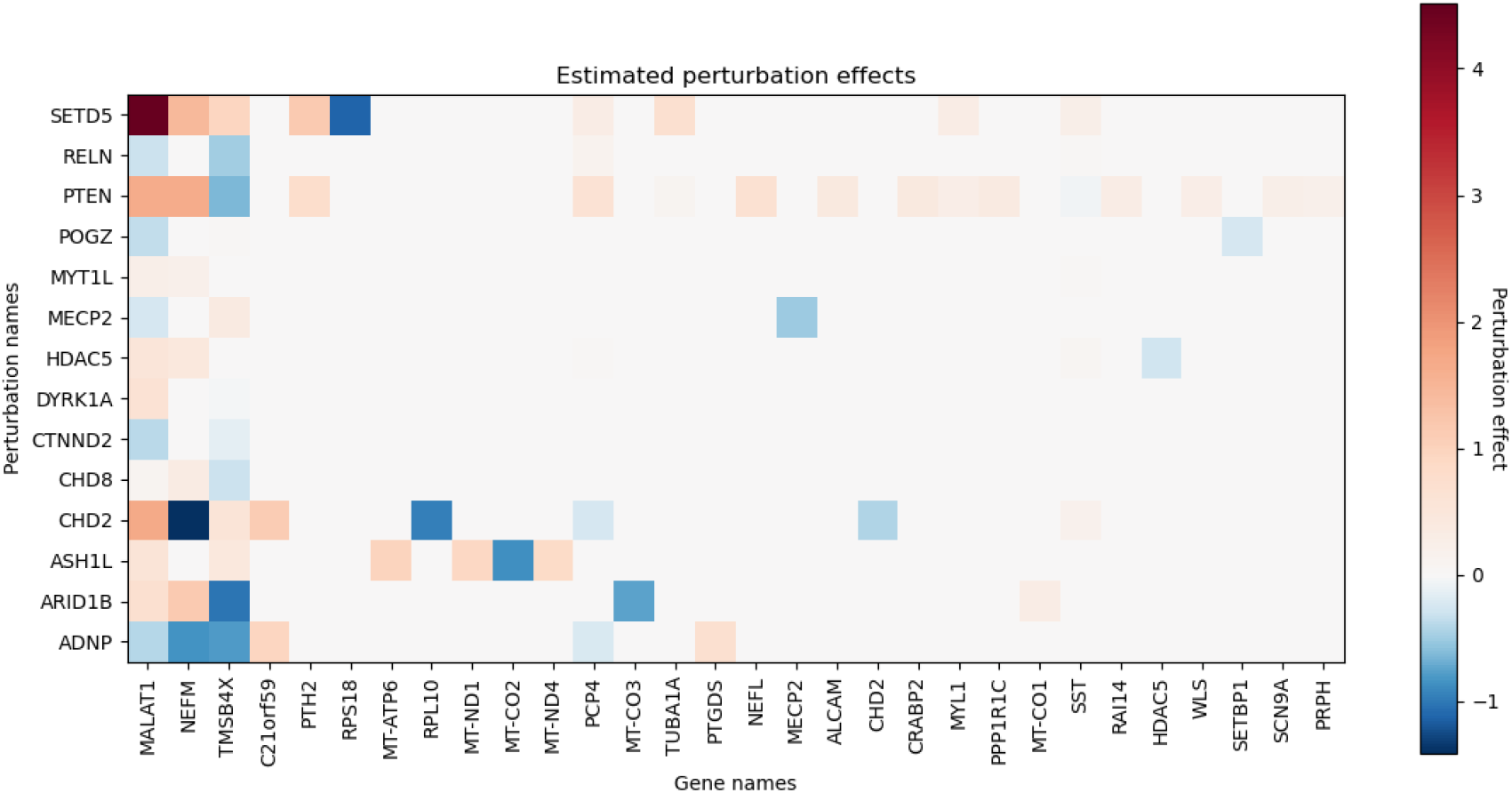
Heat map of selected perturbation effects estimated from the LUHMES dataset. Each row corresponds to one of the unique perturbation 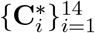.The perturbation effect of 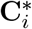 on gene *p* is included only if the associated posterior inclusion probability 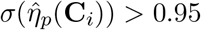.

To demonstrate scalability and versatility of our methods, we also apply Zero-inflated Poisson GPerturb to the raw counting data. We removed all zero columns from the raw counting matrix (i.e. genes with zero count across all samples), and apply Poisson GPerturb to the resulting data matrix consisting of *N* = 8708 samples and *P* = 21688 genes. The results are reported in Supplementary Fig 17, 18. Even though both the Gaussian and Poisson GPerturb fit the corresponding datasets reasonably well, the two approaches give slightly different estimates of the “active” perturbation-gene pairs, which is likely due to the data pre-processing and transformation steps. Hence we recommend users to analyse the dataset with different data pre-processings and compare their results. We also report in Supplementary Fig 19 histogram of the number of differentially expressed genes identified by Gaussian-GPerturb and existing methods similar to Supplementary Fig 5e in Zhou et al. ^11^.

##### S5.2 CD8+ T cell CROP-seq study

In this section, we apply Gaussian GPerturb to the primary human CD8+ T cells dataset (GSE119450) studied in Zhou et al. ^11^ in a similar fashion to the previous section. This study targets 20 genes associated with the T cell response, in both stimulated and unstimulated T cells. The processed dataset **X** ∈ ℝ^*N* ×*P*^ consists of *N* = 24955 samples and *P* = 6000 genes. For *i* = 1,…, *N*, the perturbations **C**_*i*_ ∈ {0, 1}^*L*^ are one-hot vectors of length *L* = 20, which correspond to the 20 targeted genes in the study, and **K**_*i*_ ∈ ℝ^*D*^ is a real vector of length *D* = 5 (`lib_size`: number of total UMI counts, `n_features`: number of genes with non-zero UMI readings, `mt_percent`: percentage of mitochondrial gene expression, `donor`: T Cell donor ID and `stimulated`: whether or not the T Cell is stimulated). In Zhou et al. ^11^, the authors hypothesised that perturbation effects are different in stimulated and unstimulated cells, and used a modified GSFA to capture such difference.

**Supplementary Figure 15:**
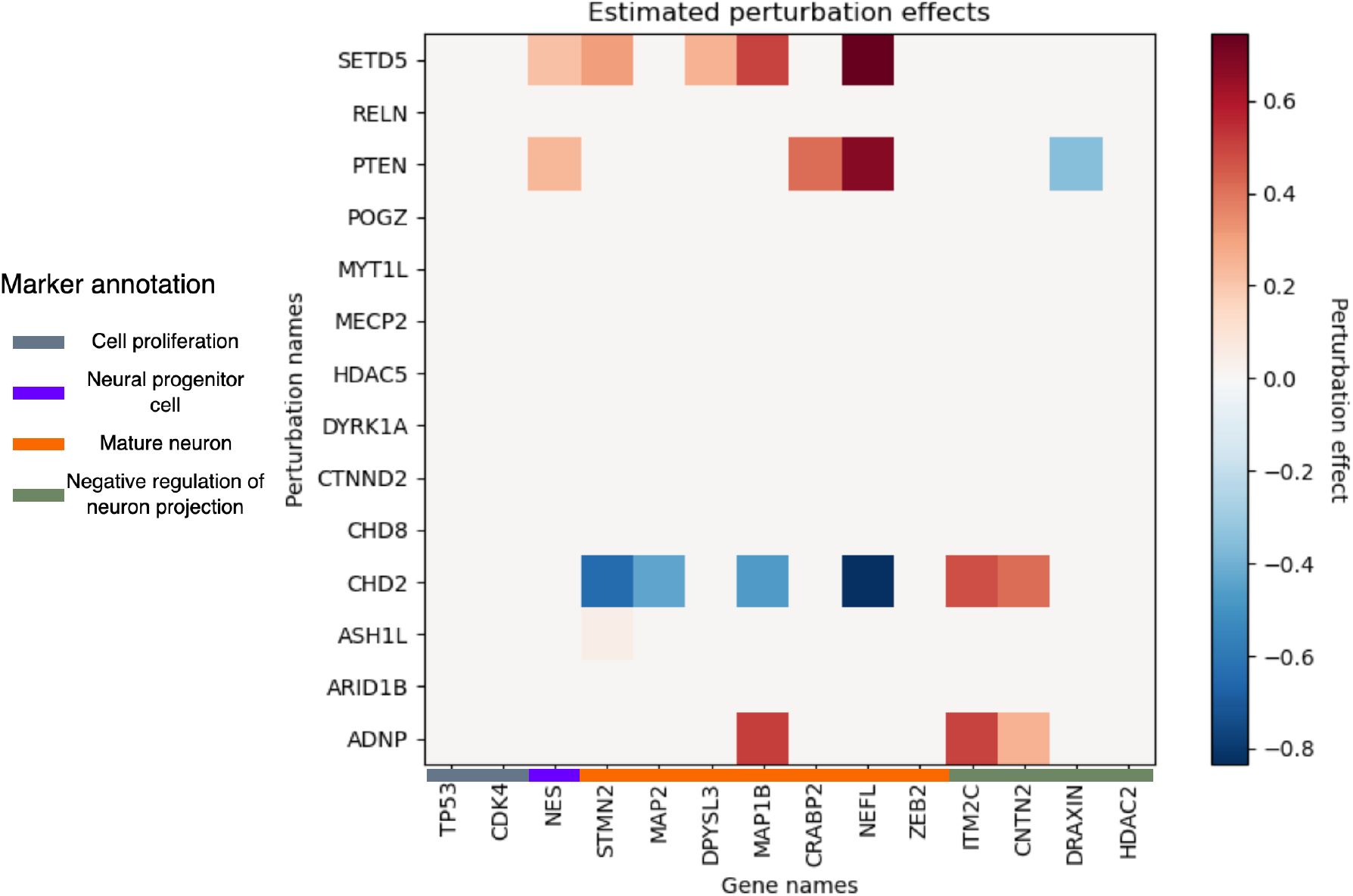
Heat map of perturbation effects on the marker genes studied in Zhou et al. ^11^ estimated from the LUHMES dataset. Each row corresponds to one of the unique perturbation 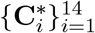. The perturbation effect of 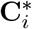 on gene *p* is included only if the associated posterior inclusion probability 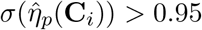.

In this example, we use a modified Gaussian GPerturb model to accommodate potentially different perturbation effects for stimulated and unsimulated T Cells in a similar fashion. Our objective is to capture potentially different perturbation effects associated with different cell groups under the same perturbation. For simplicity, we demonstrate the modified model with two cell groups, which corresponds to the stimulated and unstimulated T cells. We assume 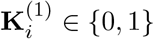,the first entry of cell information, encodes the cell group indicator. Extension to larger number of cell groups or other cell conditions is straightforward.

**Supplementary Figure 16:**
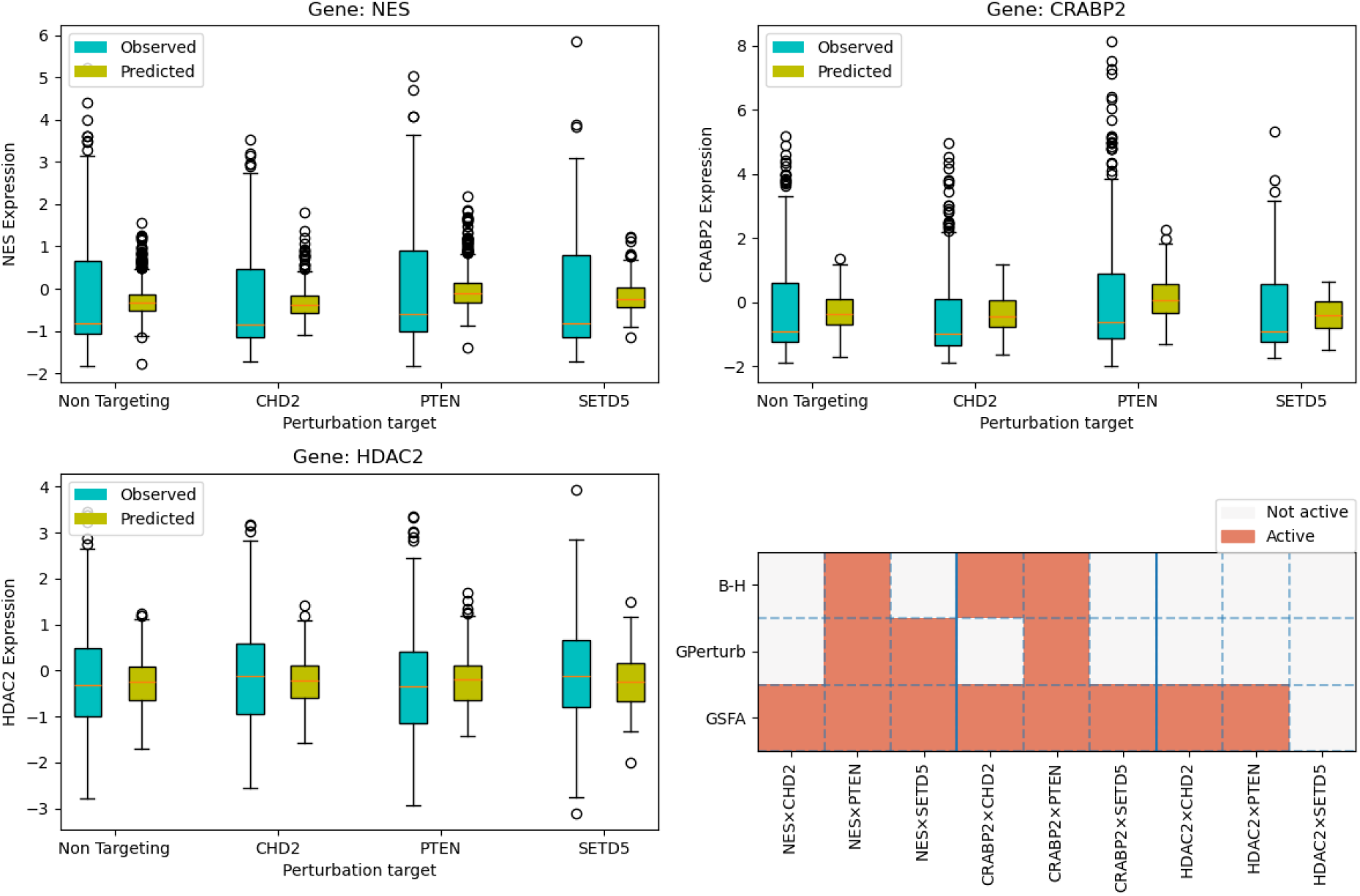
Comparing the expression levels under different perturbations. **Top left**: Boxplots of the observed and GPerturb predicted expression levels of gene NES in test set. **Top right**: Boxplots of the observed and GPerturb predicted expression levels of gene CRABP2 in test set. **Bottom left**: Boxplots of the observed and GPerturb predicted expression levels of gene HDAC2 in test set. **Bottom right**: Subset of gene-perturbation pairs selected by Benjamini–Hochberg, GPerturb and GSFA respectively.

We start from the modified generative process

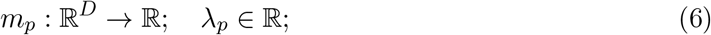

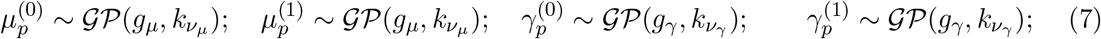

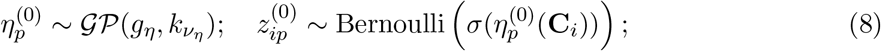

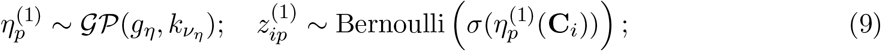

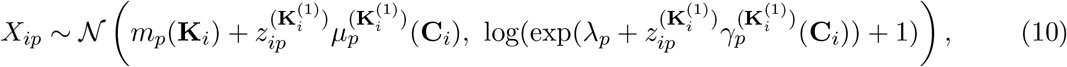

In other words, the generative process assumes that two cell groups share the common basal mean function *µ*_*p*_ (whose output also depends on the cell group indicator), but are associated with different perturbation effects 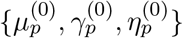 and 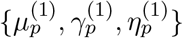.

The variational family is then modified accordingly: We replace *f*_***ξ***_ : ℝ^*L*^ → ℝ^6*P*^ in Eqn (13) by *g*_***ξ***_ : ℝ^*L*+1^ → ℝ^6*P*^, which takes the augmented perturbation vector 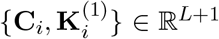 as its input. The new function *g*_***ξ***_ alongside with *f*_***ϕ***_ and ***λ*** are estimated by minimizing the ELBO given in Eqn (10) in a similar fashion to the original Gaussian GPerturb. The Poisson GPerturb is modified in a similar fashion.

**Supplementary Figure 17:**
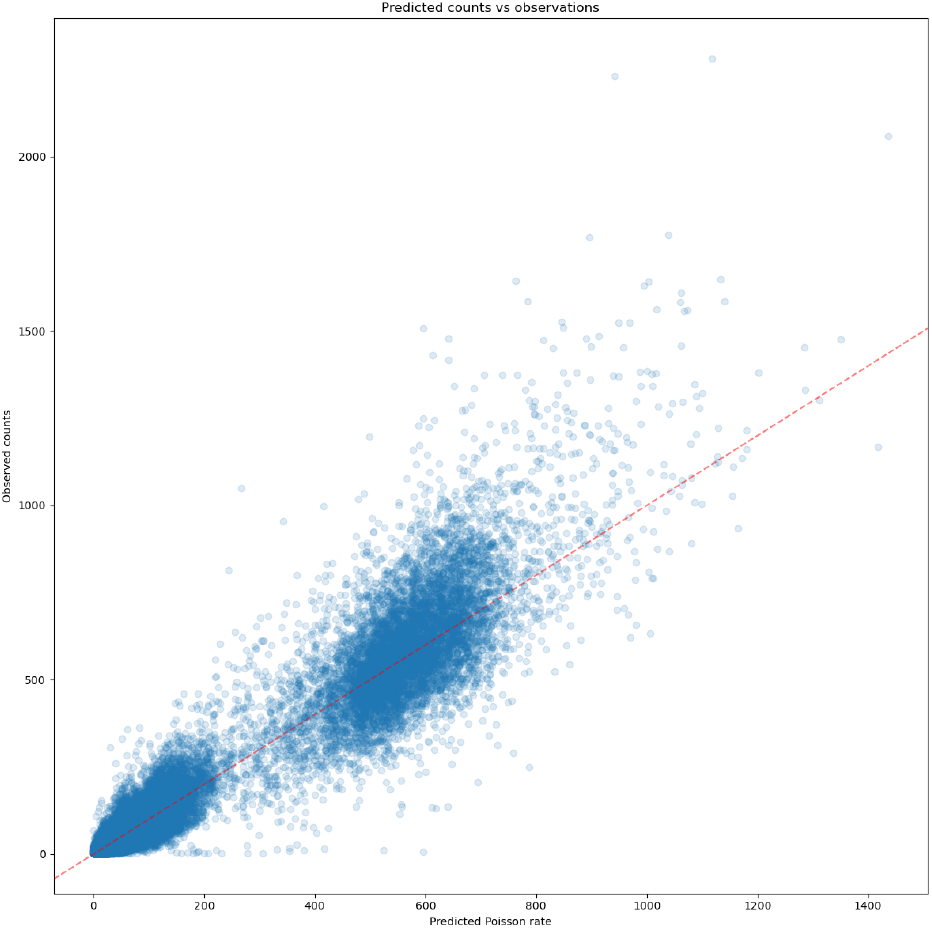
Non-zero observed counts for each cell-gene pair vs corresponding estimated Poisson rate for each cell-gene pair given by Poisson GPerturb

**Supplementary Figure 18:**
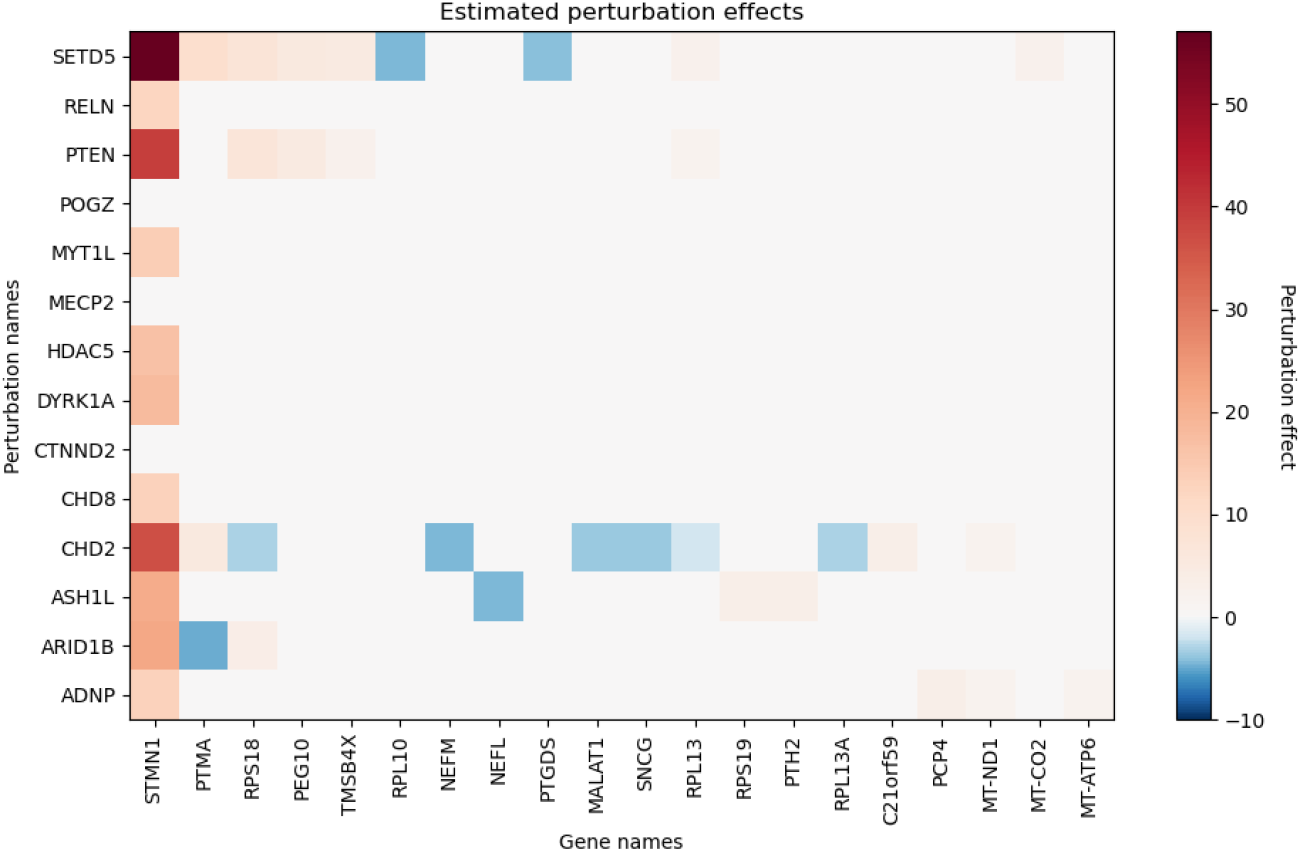
Heat map of the estimated perturbation effects given by Poisson GPerturb. Similar to Supplementary Fig 14, the perturbation effects of 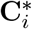 on gene *p* is included only if the associated posterior inclusion probability 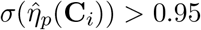.

Similar to the previous example, we randomly select 20% of the dataset as the test set, and use the rest to train GPerturb. We report the fitted results in Supplementary Fig 13. Similar to the previous section, we compare the predictive performance between GPerturb and GSFA on the cell-gene pairs in test set whose estimated posterior inclusion probability is greater than 0.95. Here we found post-processed Gaussian GPerturb and GSFA achieve comparable prediction performance (*r*_GPerturb_ = 0.271, *r*_GSFA_ = 0.335).

The heat maps of estimated perturbation effects associated with unique perturbations are reported in Supplementary Fig 20. Here we only include the top 30 genes sorted by the magnitude of overall perturbation effects in the heat map for sake of visual. We also report the heat map of perturbation effects on the collection of marker genes studied in Zhou et al. ^11^ in Supplementary Fig 21. In Supplementary Fig 22 and 23, we compare the expression level of three marker genes (IL7R, TNFRSF18, MKI67) under three perturbations (LCP2, CBLB, TCEB2) in a similar fashionn to the previous section. Again we see that GPerturb agrees better with the B-H corrected two-sample *t*-tests, and GSFA tends to pick up more perturbation-gene pairs whose mean expression levels are not significantly different from the baseline. We also apply the Zero-inflated Poisson GPerturb model to the raw counting data consists of *N* = 8708 samples and *P* = 22400 genes. The results are reported in Supplementary Fig 24, 25.

**Supplementary Figure 19:**
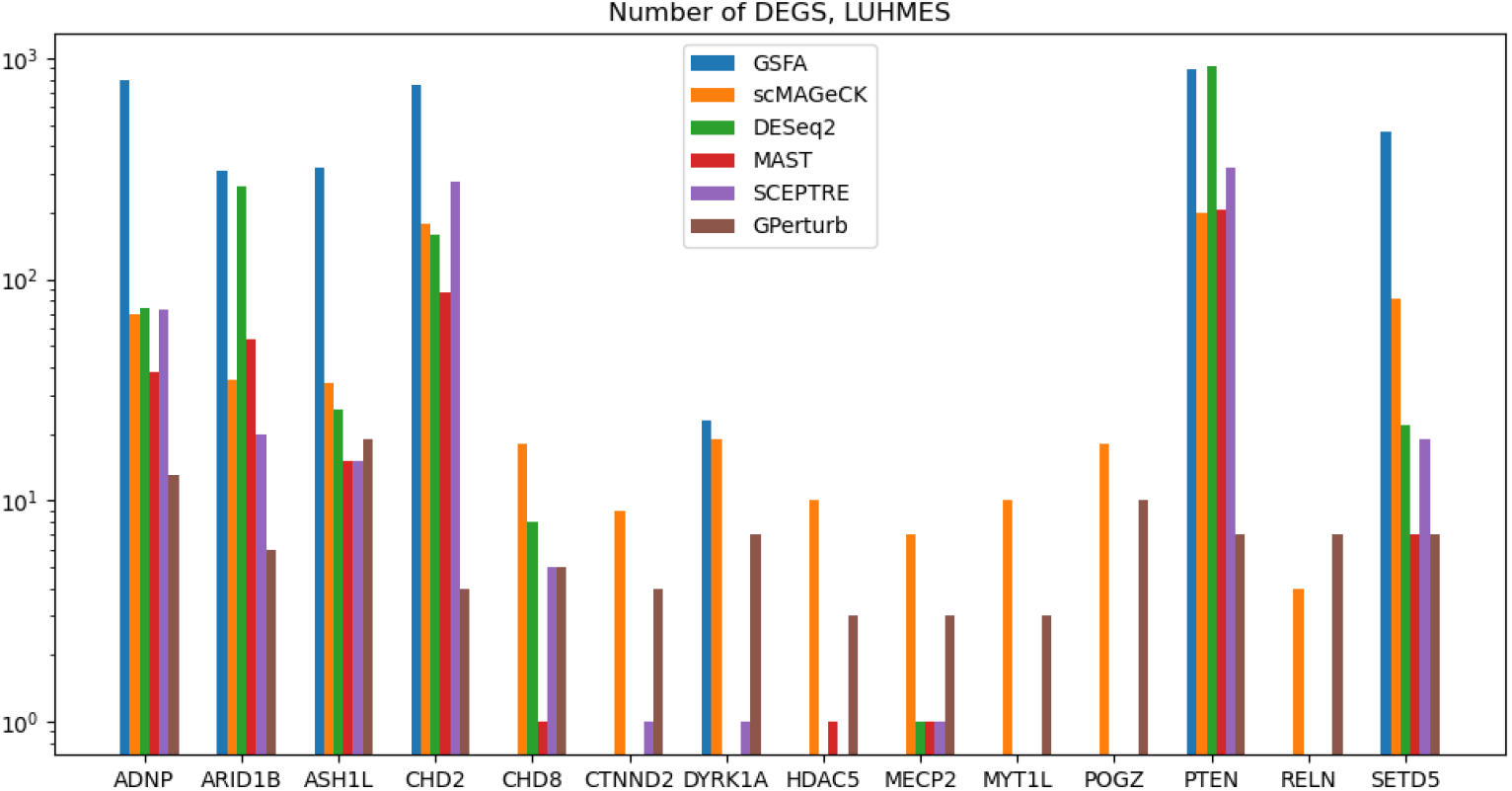
Histogram of the number of differentially expressed genes identified by different methods, LUHMES dataset. This figure is modified from Supplementary Fig 5e in ^11^.

We also applied Zero-inflated Poisson GPerturb to the raw counting data of the human T Cells dataset in Section S5.2. Results are reported in Supplementary Fig 24 and 25. We also report in Supplementary Fig 26, a histogram of the number of differentially expressed genes identified by Gaussian GPerturb and existing methods similar to Supplementary Fig 5e in Zhou et al. ^11^.

**Supplementary Figure 20:**
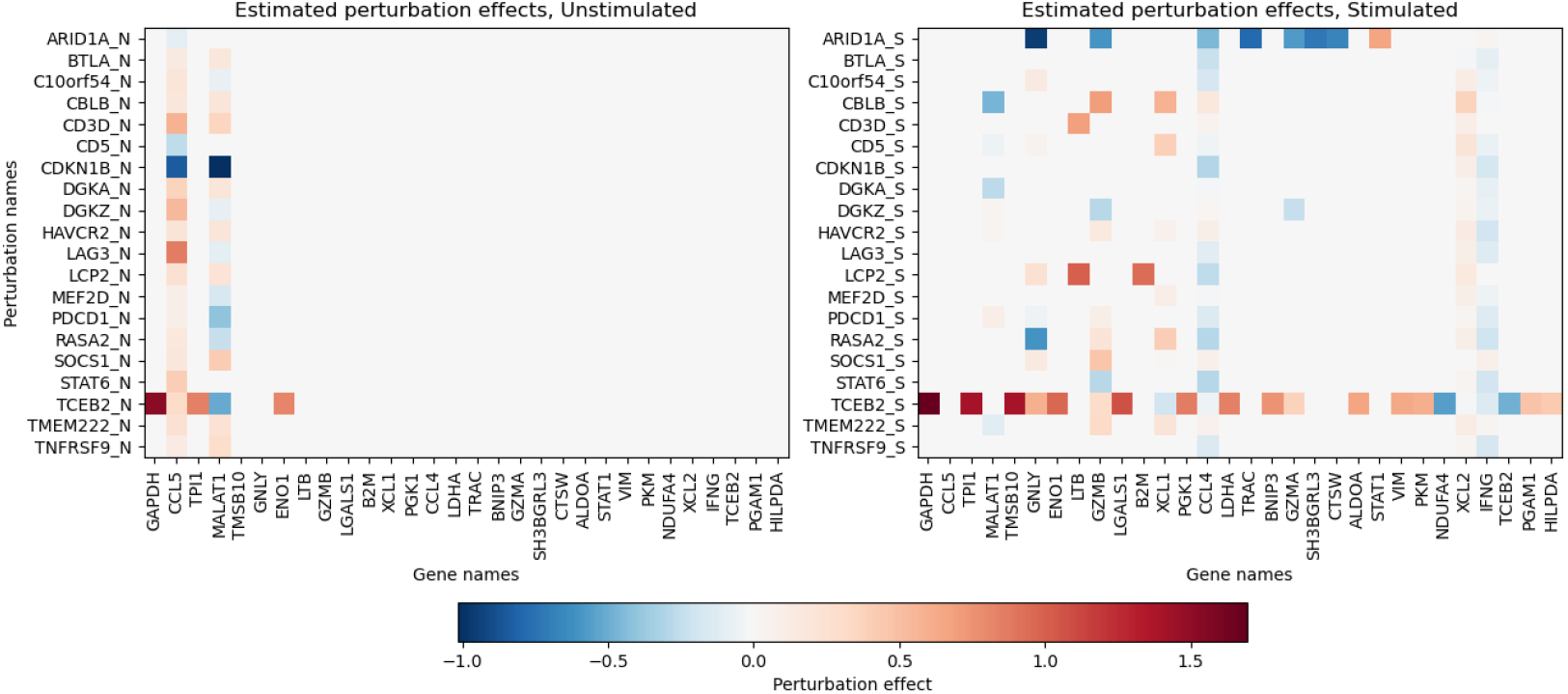
Heat map of perturbation effects estimated from the human T Cells dataset. **Left**: Estimated perturbation effects on unstimulated T cells. **Right**: Estimated perturbation effects on stimulated T cells. The perturbation effect of 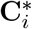 on gene *p* is included only if the associated posterior inclusion probability 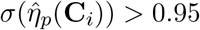. We only include the top 30 genes sorted by the magnitude of overall perturbation effects in the heat map for sake of visual.

**Supplementary Figure 21:**
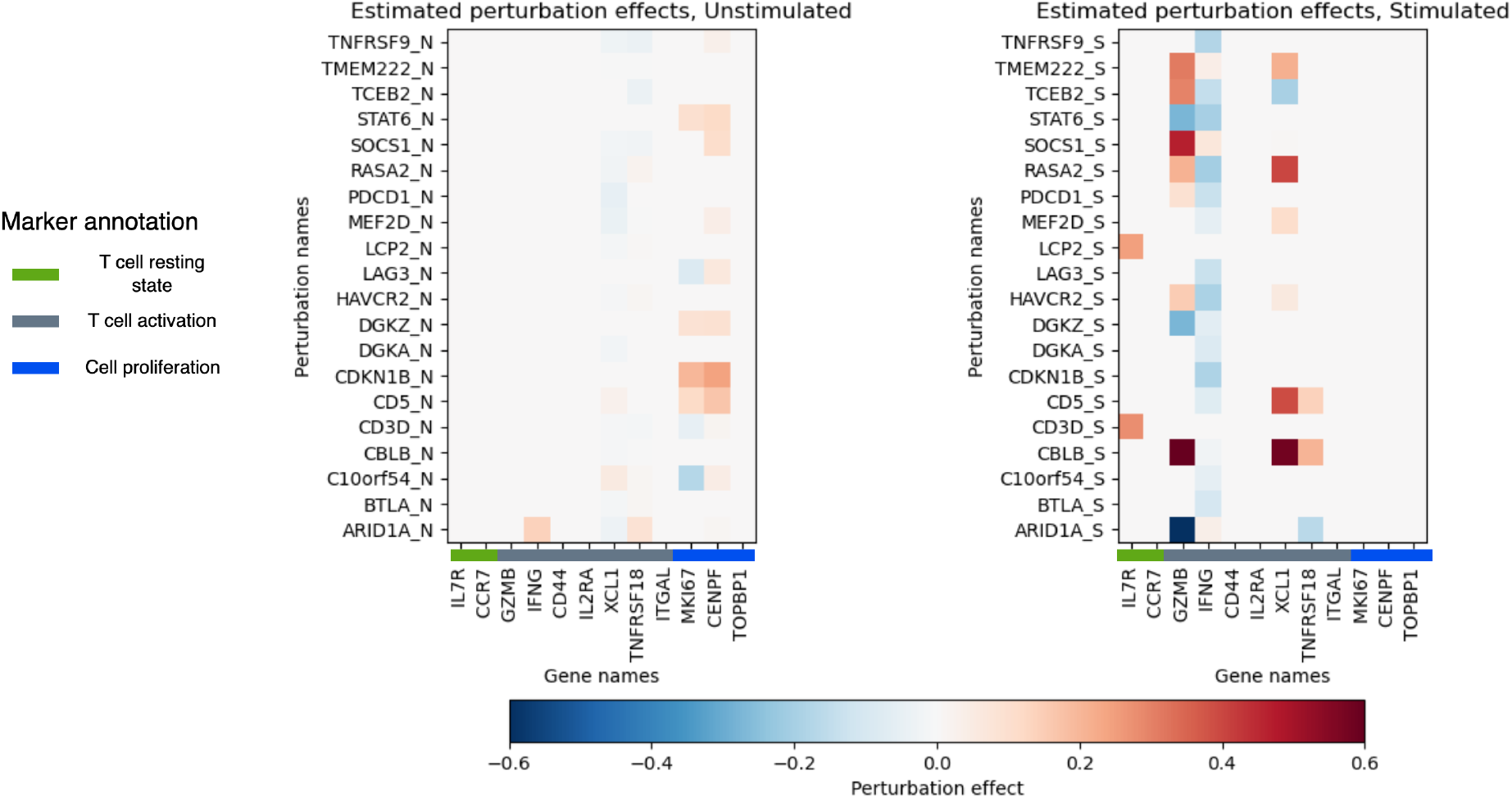
Heat map of perturbation effects on the marker genes studied in ^11^ estimated from the human T cell dataset. **Left**: Estimated perturbation effects on unstimulated T cells. **Right**: Estimated perturbation effects on stimulated T cells. Each row corresponds to one of the unique perturbation 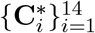.The perturbation effect of 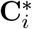 on gene *p* is included only if the associated posterior inclusion probability 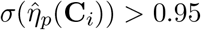.

**Supplementary Figure 22:**
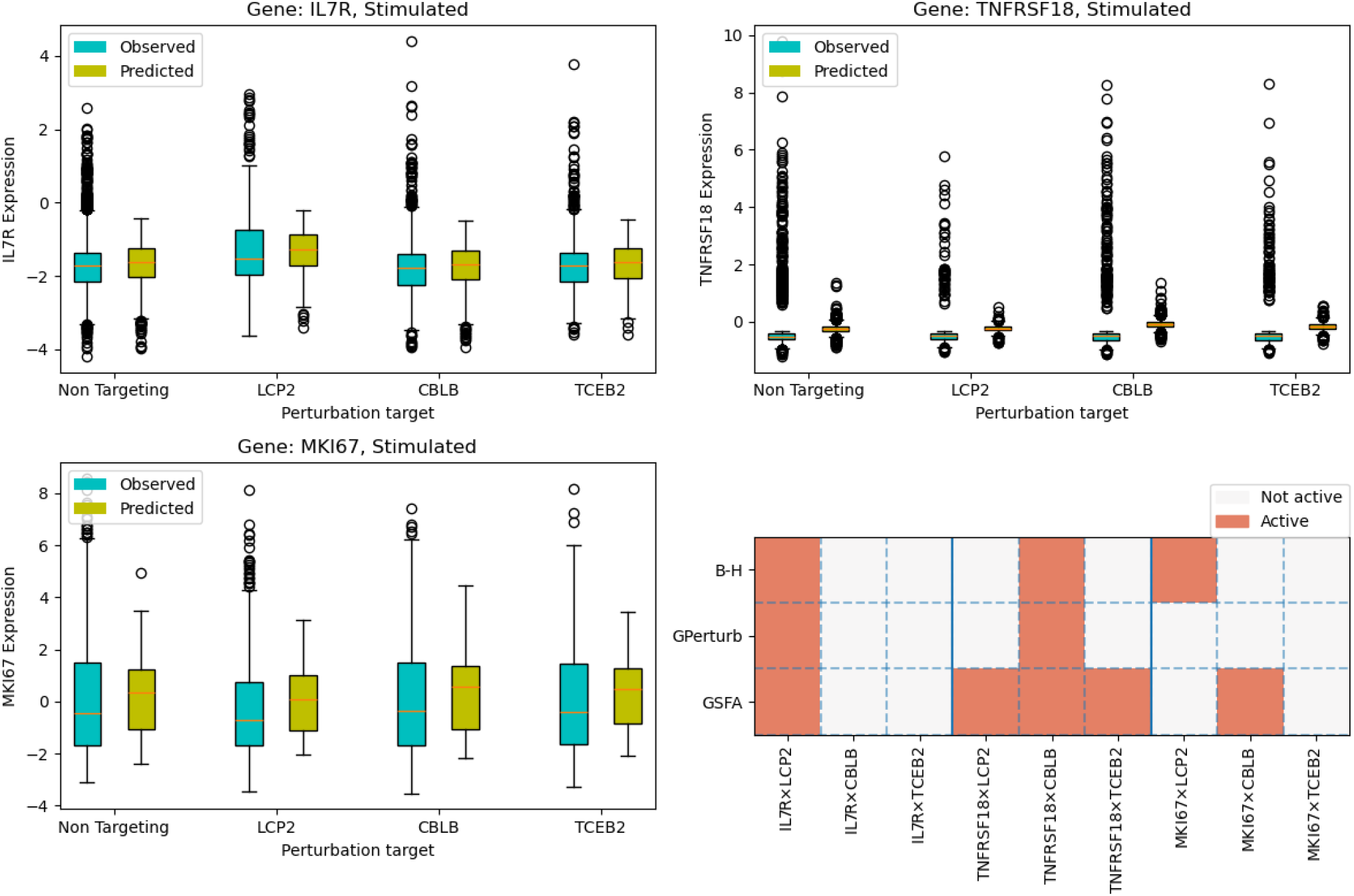
Comparing the expression levels in stimulated T cells under different perturbations. **Top left**: Boxplots of the observed and GPerturb predicted expression levels of gene IL7R in test set. **Top right**: Boxplots of the observed and GPerturb predicted expression levels of gene TNFTSF18 in test set. **Bottom left**: Boxplots of the observed and GPerturb predicted expression levels of gene MKI67 in test set. **Bottom right**: Subset of gene-perturbation pairs selected by Benjamini–Hochberg, GPerturb and GSFA respectively.

**Supplementary Figure 23:**
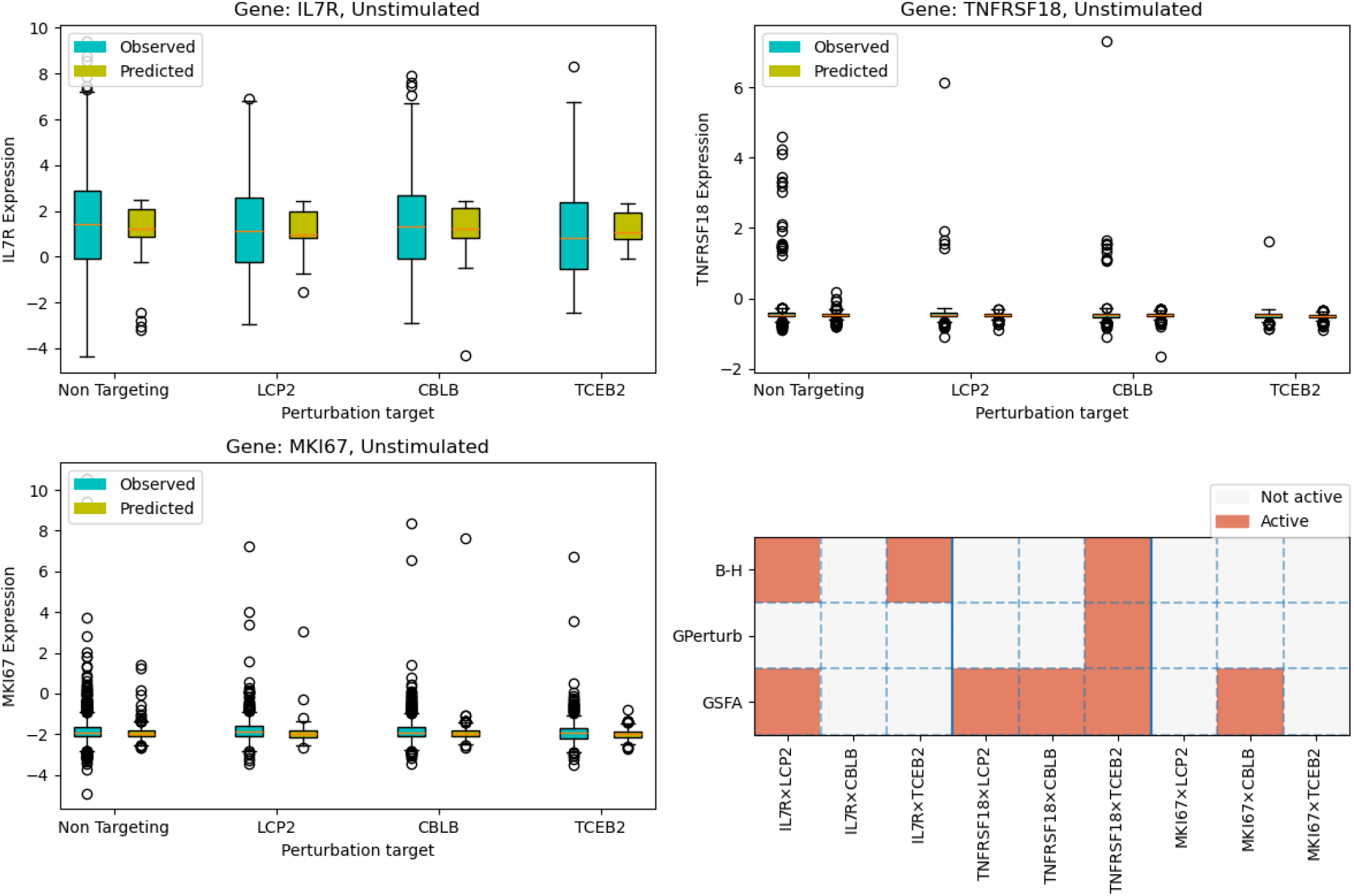
Comparing the expression levels in unstimulated T cells under different perturbations. **Top left**: Boxplots of the observed and GPerturb predicted expression levels of gene IL7R in test set. **Top right**: Boxplots of the observed and GPerturb predicted expression levels of gene TNFTSF18 in test set. **Bottom left**: Boxplots of the observed and GPerturb predicted expression levels of gene MKI67 in test set. **Bottom right**: Subset of gene-perturbation pairs selected by Benjamini–Hochberg, GPerturb and GSFA respectively.

**Supplementary Figure 24:**
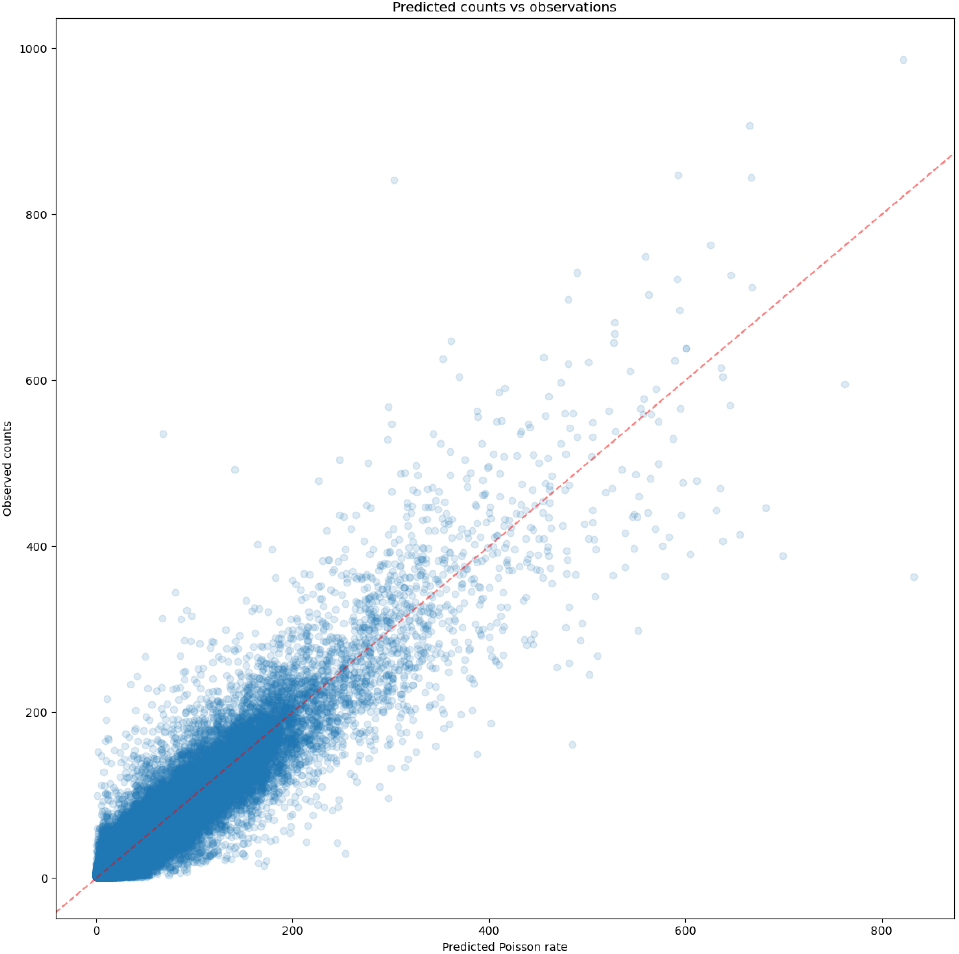
Human T Cells dataset, Non-zero observed counts for each cell-gene pair vs corresponding estimated Poisson rate for each cell-gene pair given by Poisson GPerturb

**Supplementary Figure 25:**
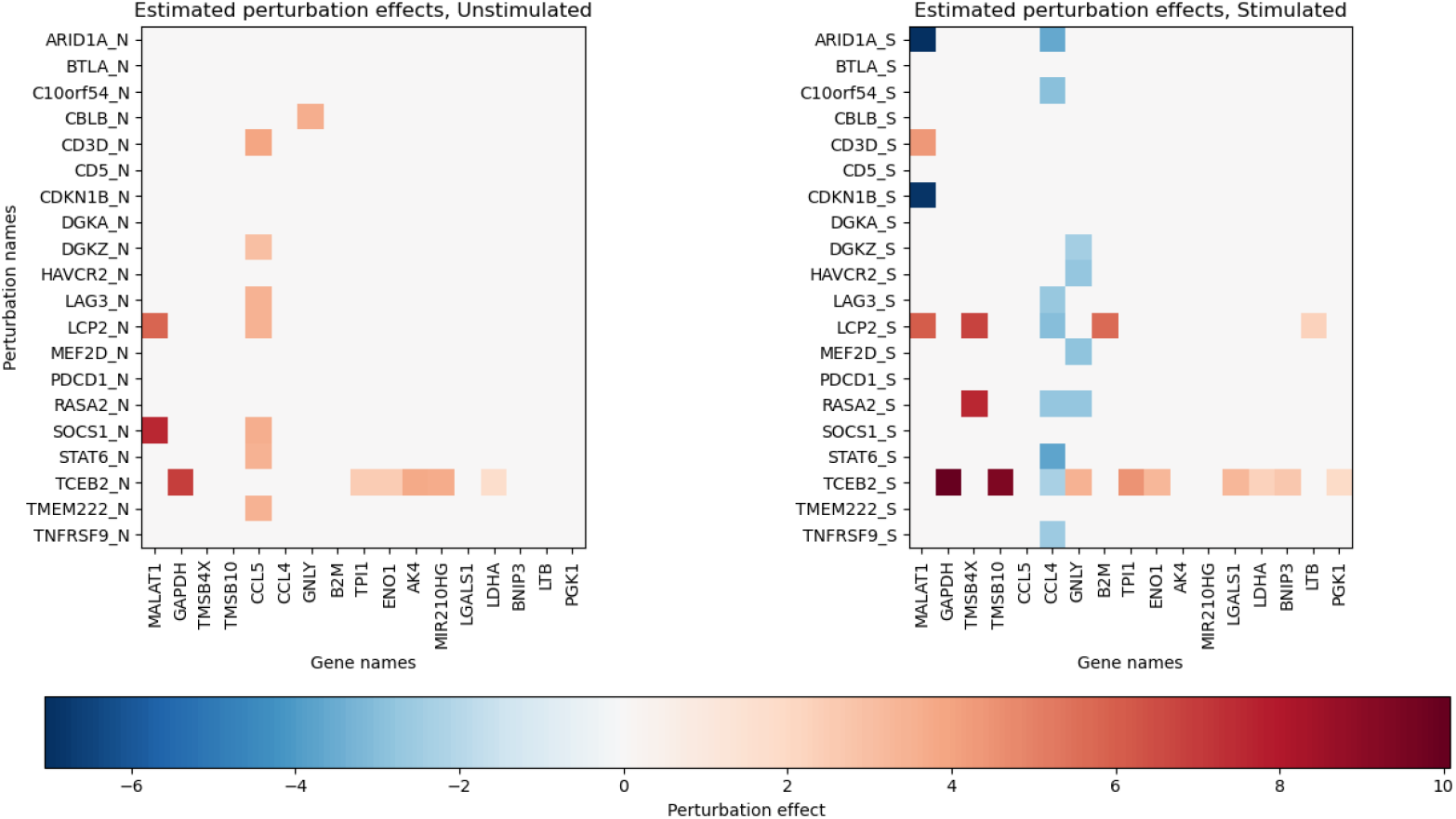
Heat map of the estimated perturbation effects given by Poisson GPerturb. Similar to Supplementary Fig 20, each row corresponds to the perturbation effects of a unique perturbation on simulated (S) or unstimulated (N) T Cells. The perturbation effects on gene *p* is included only if the associated posterior inclusion probability 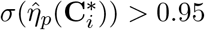.

**Supplementary Figure 26:**
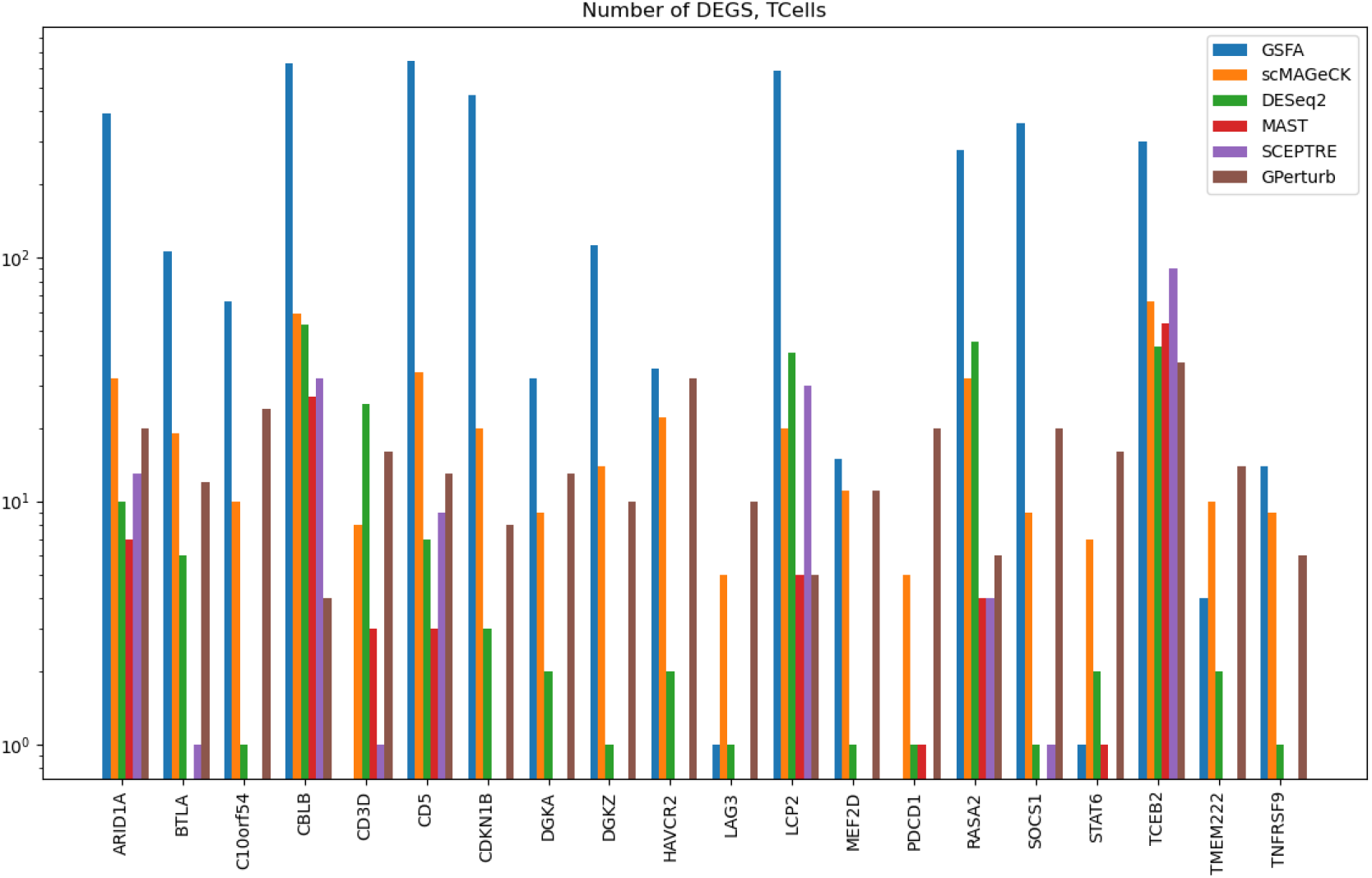
Histogram of the number of differentially expressed genes identified by different methods, human T cell dataset. This figure is modified from Supplementary Fig 5e in Zhou et al. ^11^.

## Notes

### Competing Interest Statement

The authors have declared no competing interest.

