## Supplementary figures, discussion and experiments for "GPerturb: Gaussian process modelling of single-cell perturbation data"

12 Estimated perturbation effects associated with exosome-related perturbations in Replogle et al[5] on a subset of differentially expressed genes identified by the model under different posterior inclusion probability thresholds. Note that as the threshold increases, less gene-perturbation pairs are deemed to be responsive. 16
13 LUHMES analysis. Comparison of GSFA and GPerturb predictions. . . . . . . . 18
14 LUHMES analysis. GPerturb estimated perturbation effects. . . . . . . . . . . . 19
15 LUHMES analysis. GPerturb marker gene effects. . . . . . . . . . . . . . . . . . . 20
16 LUHMES analysis. Comparison of expression levels from GSFA and GPerturb. . 21
17 LUHMES analysis. ZIP GPerturb predictions. . . . . . . . . . . . . . . . . . . . . . 22
18 LUHMES analysis. ZIP GPerturb perturbation effects. . . . . . . . . . . . . . . . . 22
19 LUHMES analysis. Comparison of differentially expressed genes by different methods. . . . . . . . . . . . . . . . . . . . . . . . . . . . . . . . . . . . . . . . . . . 23
20 T cells analysis. GPerturb estimated perturbation effects. . . . . . . . . . . . . . . 24
21 T cells analysis. GPerturb estimated marker gene perturbation effects. . . . . . . 25
22 T cells analysis. Expression effects under perturbations. . . . . . . . . . . . . . . . 26
23 T cells analysis. Comparison of expression levels. . . . . . . . . . . . . . . . . . . . 27
24 T cells analysis. ZIP GPerturb predictions. . . . . . . . . . . . . . . . . . . . . . . . 28
25 T cells analysis. ZIP GPerturb estimated perturbation effects. . . . . . . . . . . . 28
26 T cells analysis. Comparison of differentially expressed genes across methods. . . 29

### Preamble

This supplementary document is structured as follows: In Section S1, we give further details into the background and setup of the problem expanding on the Methods description. In Section S2, we discuss its connection to some existing state-of-the-art methods. We then study the properties of the method in Section S3 using simulated data. Finally, in Section S4, we provide further details of the analysis of a number of datasets including some not included in the main manuscripts, and compare its performance with state-of-the-art methods discussed in Section S2.

### S1 Further background and motivation

Our research is motivated by guided sparse factor analysis (GSFA) [11], a Bayesian sparse factor analysis model which aims to infer both the effects of genetic perturbations on individual genes, and the groups of genes or gene modules that are co-regulated. In GSFA, the groups of co-regulated genes are encoded as sparse loading vectors, and the effects of genetic perturbations on individual genes are assumed to be linear functions of the encoded loading vectors and the perturbation vectors. Before we give the details, we first introduce the notation. Let $N$ be the total number of samples (usually cells). For $i = 1, \ldots, N$, let $\mathbf{K}_i$ be the $D$ dimensional cell-level information vector, $\mathbf{C}_i$ the $L$ dimensional perturbation vector, and $\mathbf{X}_i$ the $P$ dimensional observed gene-expression vector associated with the $i$th sample, where $P$ is the total number of genes. Each entry $X_{ip}$ in $\mathbf{X}$ represents the observed expression level of the $p$th gene in the $i$th sample. Let $\mathbf{K} = \{\mathbf{K}_i\}_{i=1}^N$, $\mathbf{C} = \{\mathbf{C}_i\}_{i=1}^N$, $\mathbf{X} = \{\mathbf{X}_i\}_{i=1}^N$. In GSFA, the authors first apply a deviance-statistics transformation [9] to the raw counting data matrix, which returns to a continuous response matrix $\mathbf{X} \in \mathbb{R}^{N \times P}$. Then the cell-level information is decoupled/removed from $\mathbf{X}$ by first regressing each column of $\mathbf{X}$ on $\mathbf{K}$ using linear regression, then subtract the fitted value from $\mathbf{X}$. In other words, the resulting matrix $\tilde{\mathbf{X}}$ would be the residuals of the linear regressions described above. Finally, the author modelled the pre-processed $\tilde{\mathbf{X}}$ as

$$\tilde{\mathbf{X}} = \mathbf{C}\boldsymbol{\beta}\boldsymbol{W} + \boldsymbol{\epsilon}, \tag{1}$$

where $d$ is the number of latent factors chosen by the user, $\boldsymbol{\beta} \in \mathbb{R}^{L \times d}$ is a linear transformation of $\mathbf{C}$, $\mathbf{C}\boldsymbol{\beta}$ is the factor matrix informed by the perturbation vector $\mathbf{C}$, $\boldsymbol{W} \in \mathbb{R}^{d \times P}$ is the loading matrix, and $\boldsymbol{\epsilon} \in \mathbb{R}^{N \times P}$ is the residual matrix.

To improve interpretability, the authors put sparsity inducing priors on $\boldsymbol{\beta}$ and $\boldsymbol{W}$. Even though GSFA gives promising results on multiple real-world datasets, it suffers from a few limitations. Firstly, we do not expect a perturbation $\mathbf{C}_i$ to affect all $P$ genes in a dataset simultaneously. In particular, users are often interested in the questions "Given a perturbation, which subset of genes does it target?" Hence it would be desirable if a model could directly give sparse estimates of gene-level perturbation effects. However, even though the posterior of both $\boldsymbol{\beta}$ and $\boldsymbol{W}$ in GSFR are sparse, the resulting estimated perturbation effects on individual genes are not necessarily so, and users have to test for non-zero perturbation effects using LSFR [8], which is not straightforward to interpret. Secondly, GSFR is only applicable to continuous response matrix $\mathbf{X}$, which relies on some form of data-preprocessing. In practice, it is also of interest to study datasets in different formats. For example, users may be interested in directly analyzing the unique molecular identifier (UMI) counts instead of the continuous deviance statistics or $z$-scores. But it is not straightforward to generalize GSFR to non-Gaussian likelihoods in a computationally efficient way, which restricts GSFR's applicability. In addition, the linear assumption in GSFR may not be flexible enough to capture the complex biological process behind the experiment results, and therefore would affect its prediction power. In the following section, we propose a model to address these limitations by directly estimating sparse gene-level perturbation effects using a Gaussian process based additive model and armortized variational inference.

#### S1.1 GPerturb Schematic Models

We present schematic illustrations of the Gaussian and ZIP-GPerturb in Fig 1 and 2.

(a)

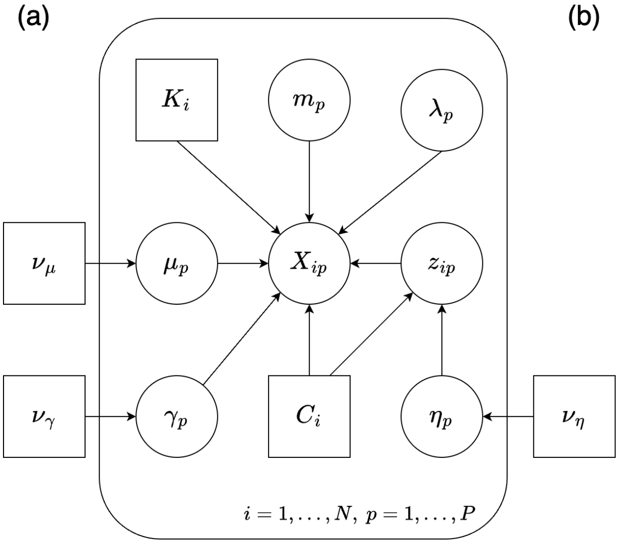

(b)

| Table of variables in the generative process | | |
|---|---|---|
| Name | Distribution | Description |
| $g_\mu, g_\gamma, g_\eta$ | Consant | Mean functions of the Gaussian processes |
| $\nu_\mu, \nu_\gamma, \nu_\eta$ | Constant | Kernel hyperparameters of the Gaussian processes |
| $C_i$ | Constant | Perturbation vector associated with the $i$th sample |
| $K_i$ | Constnat | Cell-level information vector associated with the $i$th sample |
| $m_p$ | Fixed & Unknown | Expected value of the basal expression level of the $p$th gene |
| $\lambda_p$ | Fixed & Unknown | Variability of the basal expression level of the $p$th gene |
| $\mu_p$ | Gaussian process | Perturbation effect on the expected value of expression level of the $p$th gene |
| $\gamma_p$ | Gaussian process | Perturbation effect on the variability of expression level of the $p$th gene |
| $\eta_p$ | Gaussian process | Logit score of the probability of presence of perturbation on the $p$th gene |
| $z_{ip}$ | Bernoulli | Indicator of presence of perturbation on the $p$th gene of the $i$th sample |
| $X_{ip}$ | Normal | Observed deviance statistics associated with the $p$th gene of the $i$th sample |

**Supplementary Figure 1:** (a): A graphical representation of the proposed deviance-based Normal model. (b): Table of parameters used in the proposed model.

#### S1.2 Zero-inflated Gamma-Poisson model

Here we give details of the zero-inflated Gamma-Poisson model variant:

$$m_p : \mathbb{R}^D \to \mathbb{R}; \quad \mu_p \sim \mathcal{GP}(g_\mu, k_{\nu_\mu}); \quad \eta_p \sim \mathcal{GP}(g_\gamma, k_{\nu_\gamma}); \tag{2}$$

$$\alpha_p \in \mathbb{R}^+; \quad \pi_p \in (0,1); \quad z_{ip} \sim \text{Bernoulli}\big(\sigma(\eta_p(\mathbf{C}_i))\big); \tag{3}$$

$$X_{ip} \sim \text{ZIGP}\left(\log(\exp(m_p(\mathbf{K}_i) + z_{ip}\mu_p(\mathbf{C}_i)) + 1), \alpha_p, \pi_p\right), \tag{4}$$

where $m_p, \mu_p, \eta_p, \pi_p, z_{ip}$ have the same interpretation as in the Zero-inflated Poisson model seen previously, $\alpha_p$ is the dispersion parameter associated with the $p$th gene, and $\text{ZIGP}(\mu, \alpha, \pi)$ is a Zero-inflated Gamma-Poisson distribution with p.m.f.

$$\text{ZIGP}(y; \mu, \alpha, \pi) = \pi \mathbf{1}(x = 0) + (1 - \pi)\frac{\Gamma(y + \alpha^{-1})}{\Gamma(\alpha^{-1})\Gamma(y + 1)} \left(\frac{\mu\alpha}{1 + \mu\alpha}\right)^y \left(\frac{1}{1 + \mu\alpha}\right)^{\alpha^{-1}}. \tag{5}$$

The ELBO estimate and variational posterior inference of the ZIGP model above can be carried out using the same procedure as the Zero-inflated Poisson model.

(a)

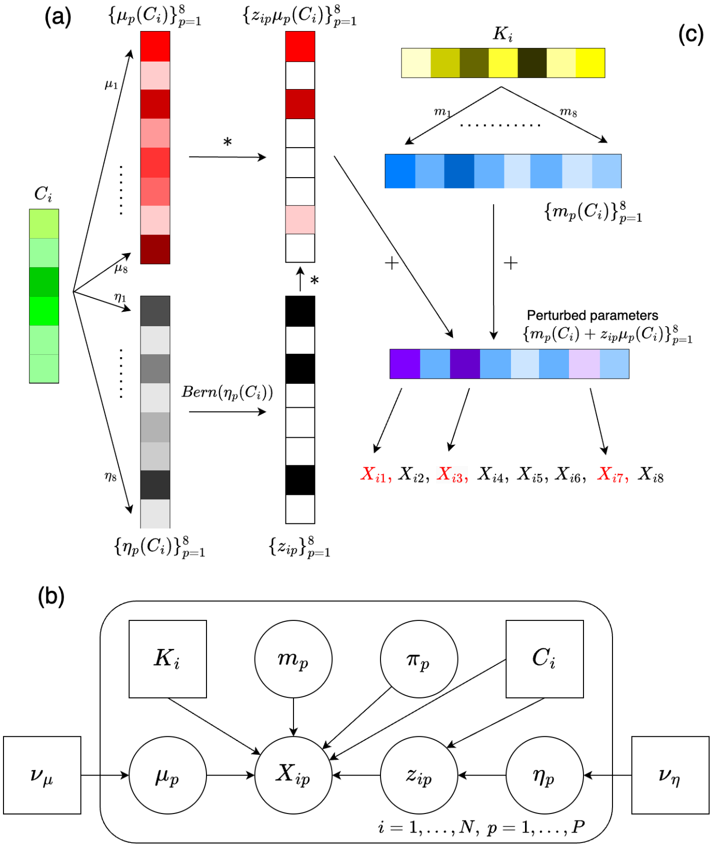

(c)

Table of variables in the generative process

| Name | Distribution | Description |
|---|---|---|
| $g_\mu, g_\eta$ | Consant | Mean functions of the Gaussian processes |
| $\nu_\mu, \nu_\eta$ | Constant | Kernel hyperparameters of the Gaussian processes |
| $C_i$ | Constant | Perturbation vector associated with the $i$th sample |
| $K_i$ | Constnat | Cell-level information vector associated with the $i$th sample |
| $m_p$ | Fixed & Unknown | Poisson rate of the basal expression level of the $p$th gene |
| $\pi_p$ | Fixed & Unknown | Proportion of excessive zeros associated with the $p$th gene |
| $\mu_p$ | Gaussian process | Perturbation effect on the Poisson rate of expression level of the $p$th gene |
| $\eta_p$ | Gaussian process | Logit score of the probability of presence of perturbation on the $p$th gene |
| $z_{ip}$ | Bernoulli | Indicator of presence of perturbation on the $p$th gene of the $i$th sample |
| $X_{ip}$ | Zero-inflated Poisson | Observed counting data associated with the $p$th gene of the $i$th sample |

**Supplementary Figure 2:** (a): A schematic illustration of the generative process of the raw counting data vector $\mathbf{X}_i$ associated with the $i$th sample. Here we assume $L = 6, D = 7, P = 8$. $*$ denotes elementwise product. The perturbed gene expressions $\{X_{i1}, X_{i3}, X_{i7}\}$ are highlighted in red. (b): A graphical illustration of the proposed Zero-inflated Poisson model. (c): Table of variables used in the proposed Zero-inflated Poisson model

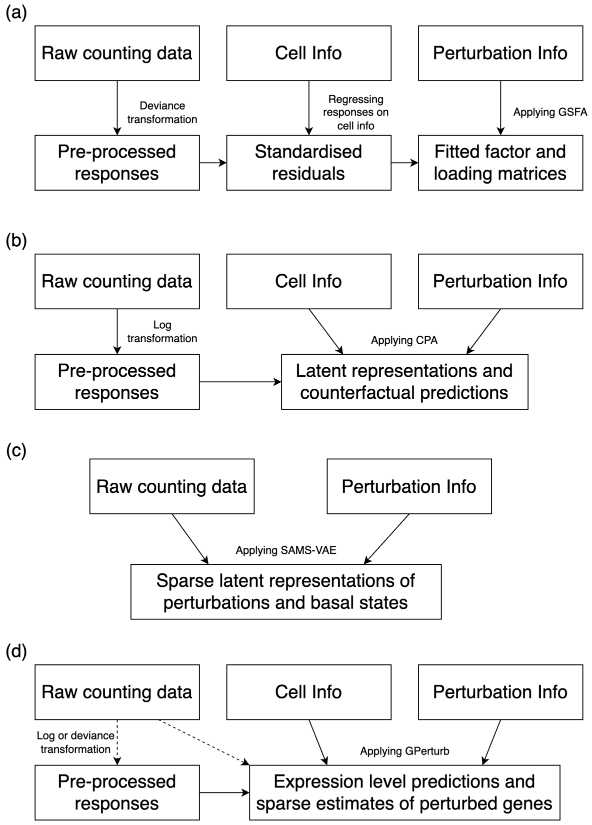

**Supplementary Figure 3:** Schematic illustration of training pipelines (a) GSFA (b) CPA (c) SAMS-VAE and (d) GPerturb

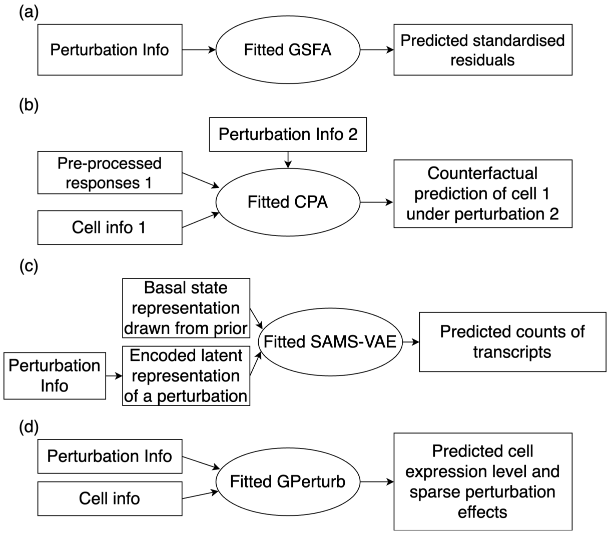

**Supplementary Figure 4:** Schematic illustration of inference/prediction pipelines (a) GSFA (b) CPA (c) SAMS-VAE and (d) GPerturb.

### S3 Simulation Experiments

Here we demonstrate that our proposed methods can learn sparse combinatorial perturbation effects using two simulated datasets.

#### S3.1 Gaussian GPerturb

We first demonstrate the efficacy of our proposed models using a simulated dataset consisting of continuous expression levels. Let the dimension of gene expression vector $P = 6000$, dimension of perturbation vector $L = 15$, dimension of cell-level information $D = 4$, number of cells $N = 4000$. The simulated data is generated as follows: Let $\mathbf{K}$ be a $N \times D$ matrix such that the entries in the first two columns of $K$ are samples drawn from i.i.d. $\mathcal{N}(0, 1)$, and the last two entries in each row of $\mathbf{K}$ are the one-hot encoding of a categorical sample drawn form Cat $\left(\{1, 2, 3\}, \{\frac{1}{3}, \frac{1}{3}, \frac{1}{3}\}\right)$. We generate $\mathbf{K}$ in this fashion since cell-level information can either be continuous or categorical in real world applications. Let $\mathbf{C}$ be a $N \times L$ binary matrix with each entry being a sample from Bernoulli(0.2). Let $\lambda_p$ be samples from i.i.d. $\mathcal{N}(0, 1)$ for $p = 1, ..., P$. Let $\mathbf{H}_1 \in \mathbb{R}^{K \times P}$, $\mathbf{H}_2, \mathbf{H}_3, \mathbf{H}_4 \in \mathbb{R}^{D \times P}$ be matrices whose entries are drawn from i.i.d. $\mathcal{N}(0, 1)$. For each $i = 1, ..., N$ and $p = 1, ..., P$, we set $m_p(K_i) = (\mathbf{KH}_1)_{ip}$ (i.e. the corresponding entry in the matrix $(\mathbf{KH}_1^T)$), $\mu_p(\mathbf{C}_i) = (\mathbf{CH}_2)_{ip}$, $\gamma_p(\mathbf{C}_i) = (\mathbf{CH}_3)_{ip}$, $z_{ip} = \mathbb{1}(\sigma((\mathbf{CH}_4)_{ip}) > 0.95)$, and $\mathbf{X}_{ip} \sim \mathcal{N}(m_p(\mathbf{K}_i) + z_{ip}\mu_p(\mathbf{C}_i), \log(\exp(\lambda_p + z_{ip}\gamma_p(\mathbf{C}_i)) + 1)$. In the simulated dataset, roughly 5% of the $z_{ip}$s are ones. In other words, roughly 5% of the sample-gene pairs in this simulated dataset are perturbed.

In order to assess the generalization performance of our proposed model on unobserved perturbation patterns, we split the simulated dataset into a training set and a test set in the following way: Let $E$ be the set consisting of 40 unique perturbation vectors uniformly and randomly selected from the rows of $C$. Let $F = \{i \in \{1, ..., N\} | \mathbf{C}_i \in E\}$ be the index set of all samples whose associated perturbation vector is in $E$. Let $\{\mathbf{X}_{-F}, \mathbf{C}_{-F}, \mathbf{K}_{-F}\}$ be the training set (i.e. samples whose indices are not in $F$), and $\{\mathbf{X}_F, \mathbf{C}_F, \mathbf{K}_F\}$ be the test set. By doing so, we ensure that the perturbation vectors in the test set are unseen in the model's training process. The size of test set is roughly 15% of the full synthetic dataset. Performance of the fitted model on test set is reported in Supplementary Fig 5. We see the model estimates both basal states and perturbation effects associated with unseen perturbation vectors accurately. In addition, the model also accurately identifies whether or not a sample-gene pair is perturbed by an unseen perturbation vectors (AUC = 0.966).

#### S3.2 Zero-inflated Poisson GPerturb

In this section, we demonstrate the efficacy of the proposed zero-inflated Poisson model using a simulated example. The dimension of the dataset and the synthetic data $\mathbf{C}, \mathbf{K}, \mathbf{H}_1, \mathbf{H}_2, \mathbf{H}_3$ are

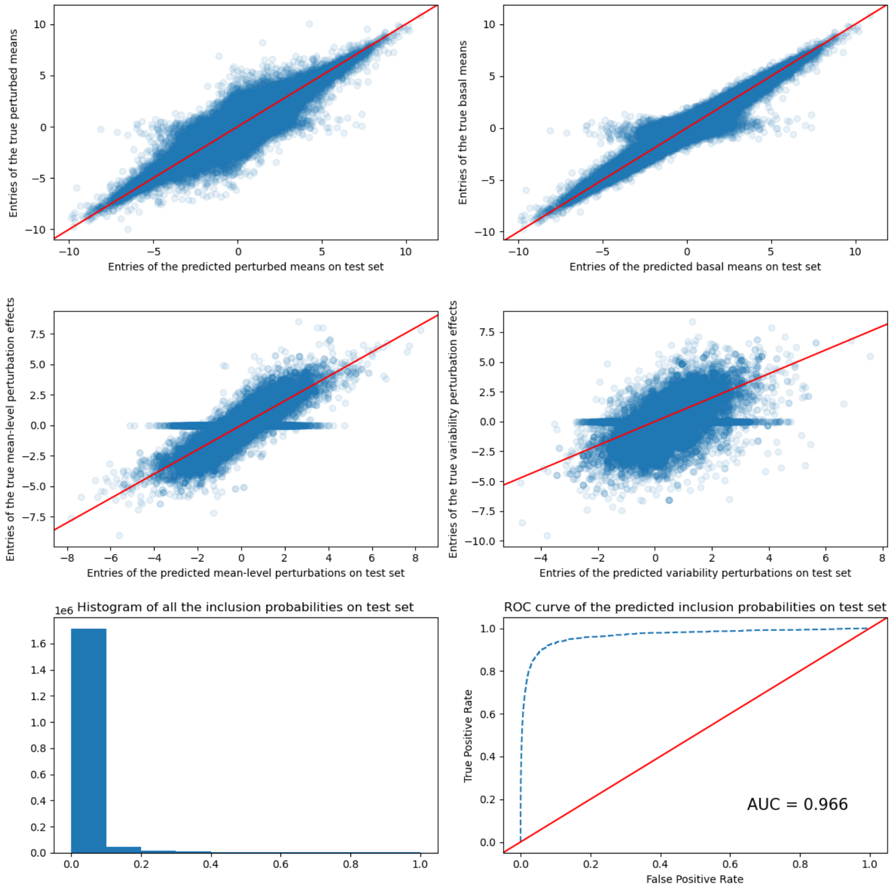

**Supplementary Figure 5:** Estimated perturbation effects and inclusion probabilities on the test set consisting of unseen perturbation patterns. Top: Scatter plots of the estimated perturbed mean v.s. the true perturbed mean and estimated basal mean v.s. true basal mean on test set. Mid: Scatter plots of estimated mean-level and variability perturbation vs the truth (i.e. $z_{ip}\mu_p(\mathbf{C}_i)$ and $z_{ip}\gamma_p(\mathbf{C}_i)$ respectively) on test set. Bottom left: Histogram of estimated inclusion probabilities on test set. Bottom right: ROC curve and AUC indicating how well the estimated inclusion probabilities predict the binary toggle $z_{ip}$s on test set.

chosen and generated in the same fashion as in the previous example. Let $\pi_p$ be samples from i.i.d. Beta$(2, 10)$ for $p = 1, ..., P$. In this example, we set $m_p(\mathbf{K}_i) = 5(\mathbf{KH}_1)_{ip} + 50$, $\mu_p(\mathbf{C}_i) = 5(\mathbf{CH}_2)_{ip}$, $z_{ip} = \mathbb{1}(\sigma((\mathbf{CH}_3)_{ip}) > 0.95)$ and $\mathbf{X}_{ip} \sim \text{ZIP}(\log(\exp(m_p(\mathbf{K}_i) + z_{ip}\mu_p(\mathbf{C}_i)) + 1), \pi_p)$. The synthetic basal rates and perturbation effects are scaled to mimic the size of counts in real datasets. The training and test set are split in the same way as in the previous example. Performance of the fitted model on test set is reported in Supplementary Fig 6. We see it predicts unseen basal states and perturbation patterns accurately, and is able to correctly identify the sparse perturbation effects of unseen perturbation vectors (AUC = 0.901).

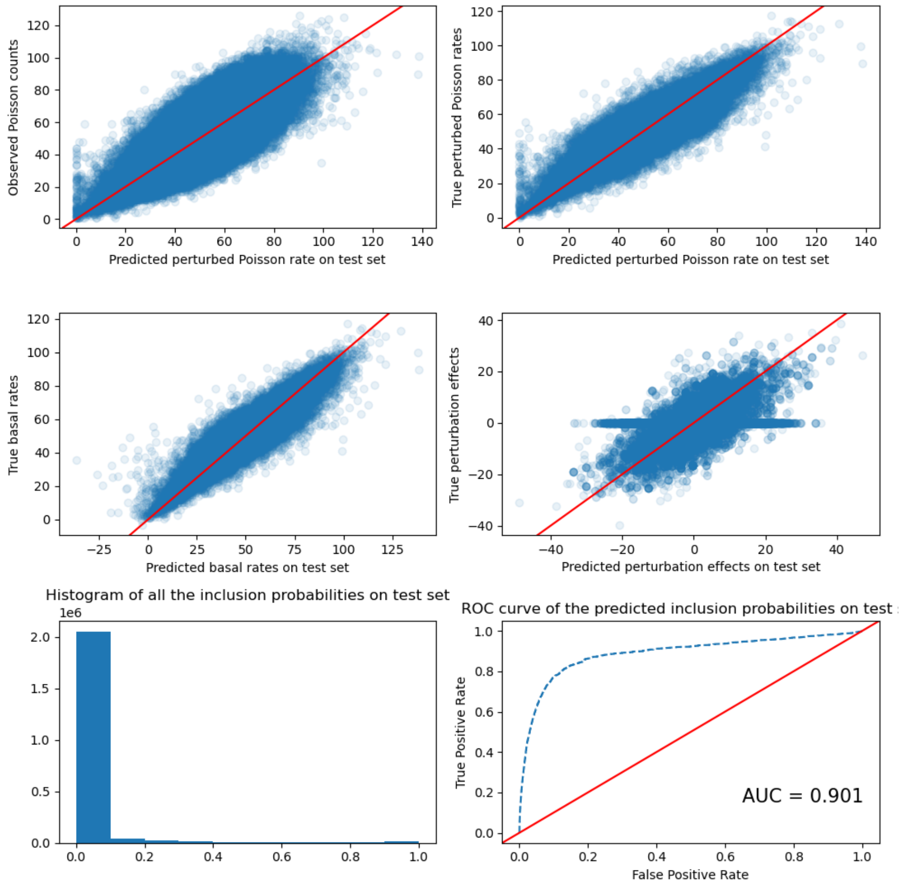

**Supplementary Figure 6:** Estimated perturbation effects and inclusion probabilities on the test set consisting of unseen perturbation patterns. Top: Scatter plots of the estimated perturbed Poisson mean v.s. the true observations and the estimated perturbed Poisson mean v.s. true Poisson means of the non-zero entries in test set. Mid: Scatter plots of the estimated basal rate v.s. true basal rate and estimated mean-level perturbation vs the truth perturbation effects of the non-zero entries in test set. Bottom left: Histogram of estimated inclusion probabilities of the non-zero entries in test set. Bottom right: ROC curve and AUC indicating how well the estimated inclusion probabilities predict the binary toggle $z_{ip}$s of the non-zero entries in test set.

### S4 Further Results

#### S4.1 Zero-inflated Gamma-Poisson GPerturb

In this section, we provide further details of the analysis of datasets discussed in the main text using the Zero-inflated Gamma Poisson variant of GPerturb. Supplementary Figures 7, 8, 9 and 10 show comparative performance between this and the Zero-inflated Poisson model used in the main text. Supplementary Figure 11 shows the empirical distribution of the estimated dispersion parameters $\alpha_p$) across all genes. The results indicate that due to the low dispersion, the simpler Zero-inflated Poisson model used in the main text is sufficient for these datasets.

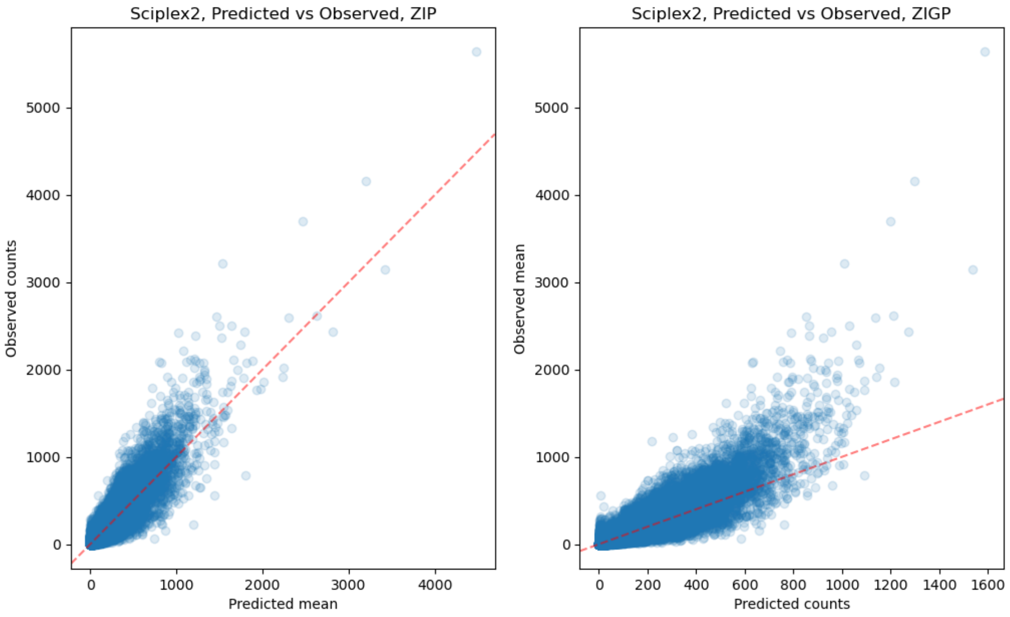

**Supplementary Figure 7:** Non-zero observed counts for each cell-gene pair vs corresponding estimated mean for each cell-gene pair given by ZIP and ZIGP GPerturb on test set.

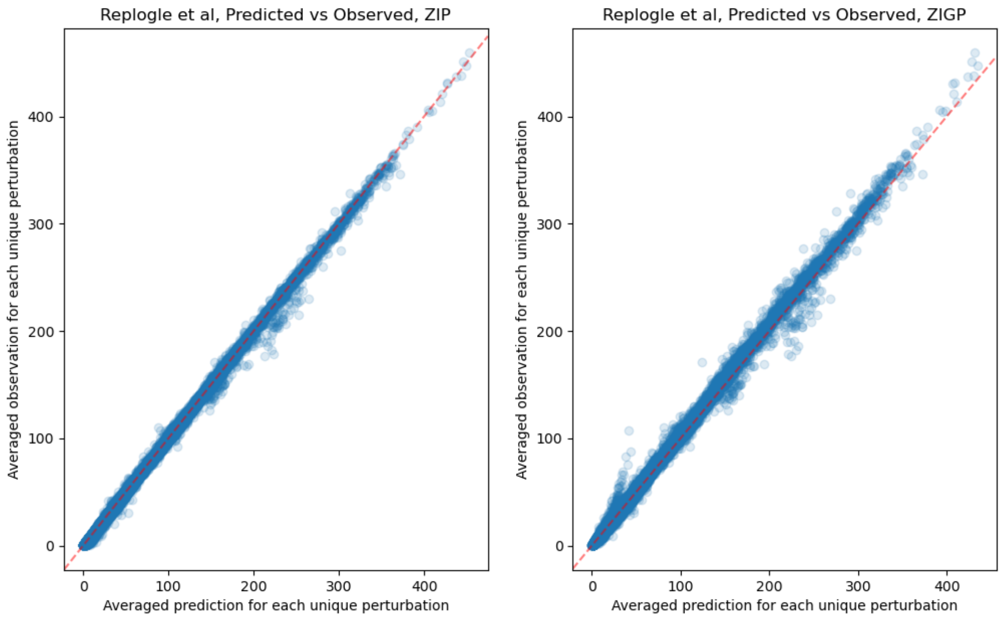

**Supplementary Figure 8:** Non-zero observed counts for each cell-gene pair vs corresponding estimated mean for each cell-gene pair given by ZIP and ZIGP GPerturb on test set.

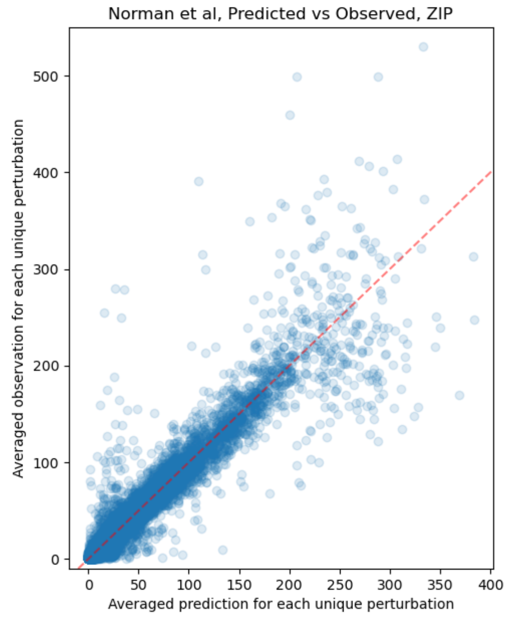

**Supplementary Figure 9:** Non-zero observed counts for each cell-gene pair vs corresponding estimated mean for each cell-gene pair given by ZIP and ZIGP GPerturb on test set.

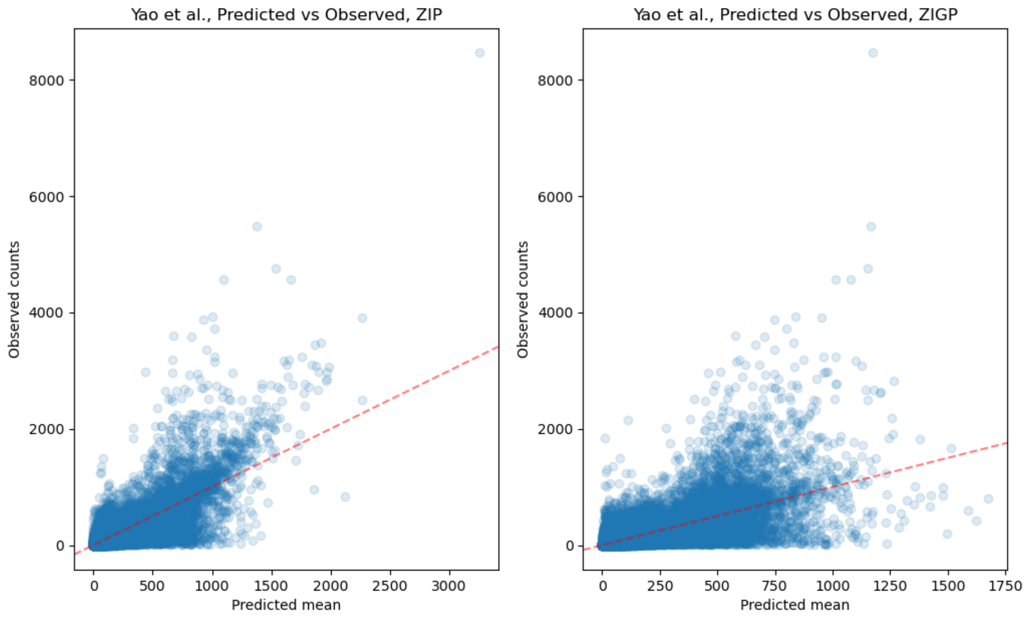

**Supplementary Figure 10:** Non-zero observed counts for each cell-gene pair vs corresponding estimated mean for each cell-gene pair given by ZIP and ZIGP GPerturb on test set.

#### S4.2 Computation cost

In this section we compare the computation cost of GPerturb with existing methods. Since distinct modeling architecture are used in different methods, we choose to report the wall clock

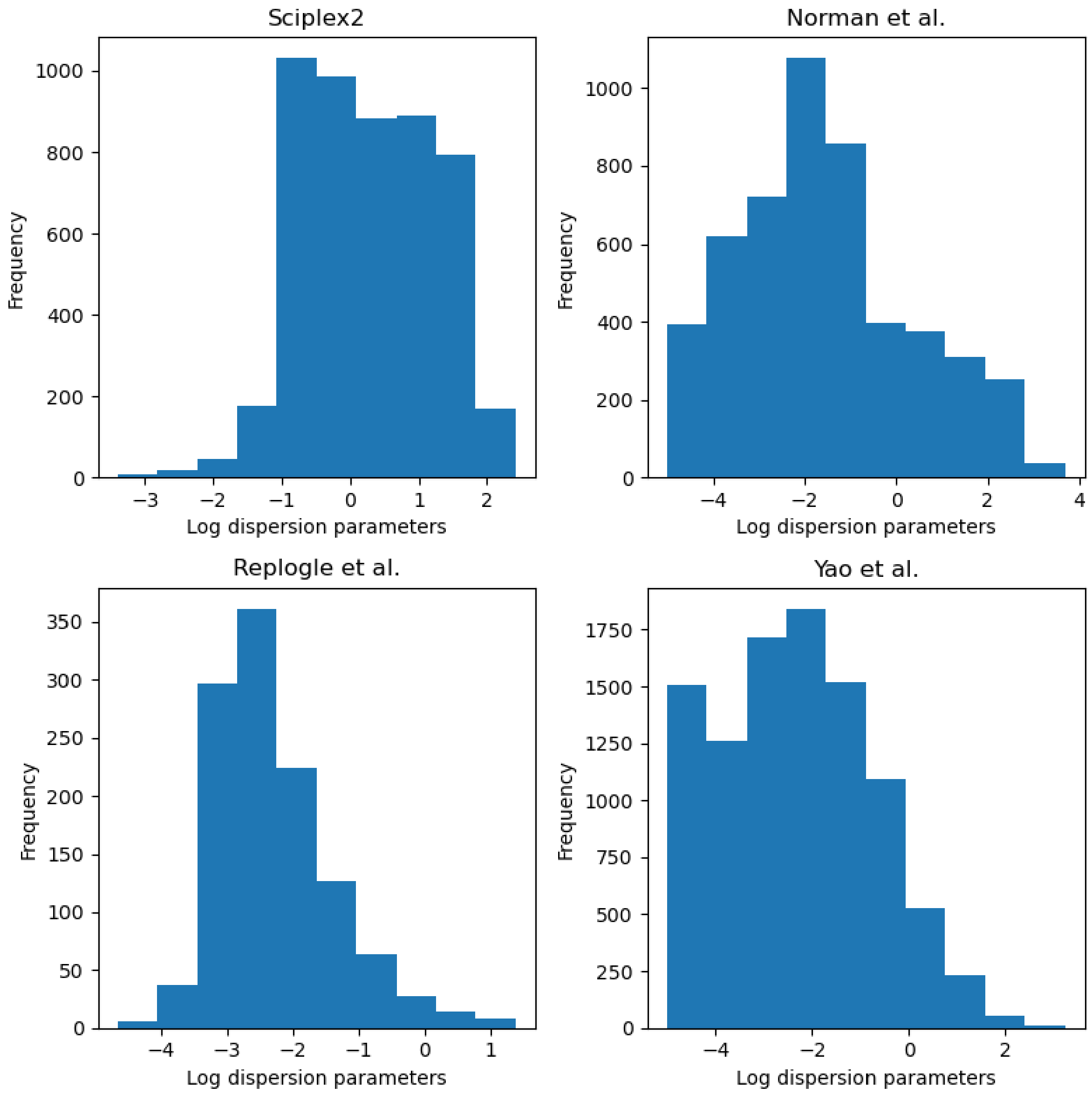

**Supplementary Figure 11:** Histograms of estimated log dispersion parameters for each of the datasets we studied in the paper.

running time as a fair benchmark for computation cost (Supplementary Table 1). We see that the computation cost of GPerturb in term of running time is on a similar level to existing methods. All experiments are conducted on our machine with an AMD Ryzen 7 2700 CPU and an NVidia RTX 2060 GPU.

##### S4.3 GPerturb's Bayesian probabilistic modeling framework

The probabilistic Bayesian modeling framework improves interpretability and uncertainty quantification of the results. Existing methods such as SAMS-VAE[2] and CPA[3] focus on predicting counterfactual perturbed gene expressions using latent embeddings which lack natural biological meaning. GEARS[6] is able to predict the effect of a given perturbation on individual genes, and give ad-hoc uncertainty quantification. However, the lack of sparsity in GEARS predictions

| Expression Input Type | Approach | Dataset | | | |
|---|---|---|---|---|---|
| | | Sciplex2[7] | Single-gene perturbation[5] | Multi-gene perturbation[4] | Multi-gene perturbation[10] |
| Continuous, transformed | GPerturb-Gaussian | $1.65 \times 10^3$ | $1.35 \times 10^4$ | $7.82 \times 10^4$ | $3.78 \times 10^3$ |
| | CPA-logsig | $1.38 \times 10^3$ | $2.27 \times 10^4$ | $1.01 \times 10^5$ | - |
| | CPA-MLP | $1.40 \times 10^3$ | $2.28 \times 10^4$ | $1.03 \times 10^5$ | - |
| | GEARS | - | $1.17 \times 10^4$ | $5.94 \times 10^4$ | $4.29 \times 10^3$ |
| Count-based | GPerturb-ZIP | $1.60 \times 10^3$ | $1.32 \times 10^4$ | $8.80 \times 10^4$ | $3.71 \times 10^3$ |
| | GPerturb-ZIGP | $1.64 \times 10^3$ | $1.38 \times 10^4$ | $8.87 \times 10^4$ | $3.82 \times 10^3$ |
| | SAMS-VAE | - | $2.09 \times 10^4$ | $6.55 \times 10^4$ | - |

**Supplementary Table 1: Comparison of wall clock running time**. Values show the averaged wall clock running time in seconds for each method and each data set over 3 repetitions.

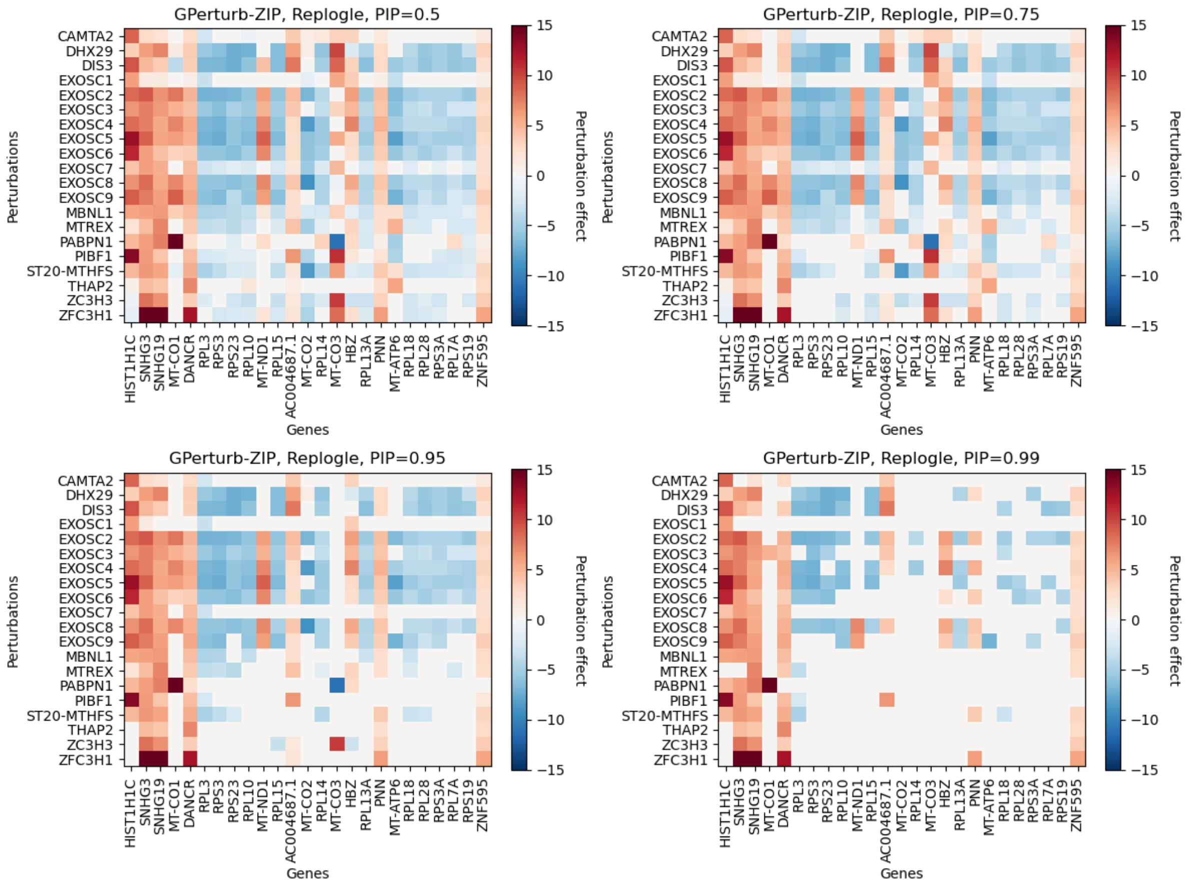

**Supplementary Figure 12:** Estimated perturbation effects associated with exosome-related perturbations in Replogle et al[5] on a subset of differentially expressed genes identified by the model under different posterior inclusion probability thresholds. Note that as the threshold increases, less gene-perturbation pairs are deemed to be responsive.

14 distinct perturbations). The cell information $\mathbf{K}_i \in \mathbb{R}^D$ is a real vector of length $D = 4$ (`lib_size`: number of total UMI counts, `n_features`: number of genes with non-zero UMI readings, `mt_percent`: percentage of mitochondrial gene expression and `batch`: batch ID). In addition to the one-hot perturbations, the dataset also consists of negative control gRNAs whose perturbations are encoded as $\mathbf{C}_i = \mathbf{0}$. Recall that our choices of generative process and variational family ensure that the negative controls with $\mathbf{C}_i = \mathbf{0}$ have zero perturbation effects. By doing so, users can view the negative controls as the baseline level, and the perturbation effects associated with the non-zero $\mathbf{C}_i$s as the perturbation strength relative to the negative controls. In Zhou et al.[11], the authors first remove cell level information from the transformed responses by regressing $\mathbf{X}$ on $\mathbf{K}$ using linear regression, then apply GSFA to the corresponding standardized residual matrix. In contrast, our proposed method disentangles and estimates cell-level and perturbation-induced variations simultaneously, and does not require any standardisation. For our proposed method, we randomly select 20% of the dataset as the test set, and use the rest to train GPerturb. The default priors discussed in Methods are used here. For GSFA, the results are obtained based on the recommended settings given in Zhou et al.[11].

In this example, we are interested in comparing the perturbation-induced variations captured by the two methods. Compared with our approach, GSFA requires additional dataset-dependent pre-processing and standardisation steps, which could potentially restrict its interpretability and generalisation power. (See Supplementary Fig 3, 4 for an illustration of the different training and inference pipelines of the methods discussed in this paper.) To make the results of the two methods comparable, we apply the following transformations to the fitted Gaussian GPerturb, mimicking the pre-processing steps in GSFA[11]: Let $N' = 0.2\lfloor N \rfloor$ be the number of samples in the test set. For each sample $i = 1, \ldots, N'$ in the test set, we first let $\bar{\mathbf{X}}' = \{\mathbf{X}_{ip} - \hat{m}_p(\mathbf{K}_i)\}_{i,p=1}^{N',P}$ to be the residual matrix (i.e. subtract the estimated cell-level variation $\hat{m}_p(\mathbf{K}_i)$ from the observed response $\mathbf{X}_{ip}$), then we standardise the columns of $\bar{\mathbf{X}}'$, and apply the same standardisation to the estimated mean perturbation effect matrix $\hat{\boldsymbol{\mu}}' = \{\sigma(\hat{\eta}_p(\mathbf{C}_i))\hat{\mu}_p(\mathbf{C}_i)\}_{i,p=1}^{N',P}$. Let $\bar{\mathbf{X}}'_{\text{GPerturb}}$ and $\hat{\boldsymbol{\mu}}'_{\text{GPerturb}}$ be the standardised residual matrix and estimated mean perturbation effect matrix given by GPerturb. Let $\bar{\mathbf{X}}_{\text{GSFA}}$ and $\hat{\boldsymbol{\mu}}_{\text{GSFA}}$ be the corresponding standardised residual matrix and estimated gene-level perturbation effect given by GSFA on the full dataset (GSFA uses the *entire* dataset to estimate the factor/loading matrices). In other words, one can view the standardised residuals $\bar{\mathbf{X}}_{\text{GSFA}}$, $\bar{\mathbf{X}}'_{\text{GPerturb}}$ as transformed, noisy observations associated with perturbation treatments, and $\hat{\boldsymbol{\mu}}_{\text{GSFA}}$, $\hat{\boldsymbol{\mu}}'_{\text{GPerturb}}$ as the estimated mean perturbation effects. We stress that the fitted Gaussian GPerturb is more interpretable on the original scale, and transformations applied to it may affect its prediction power. The purpose of the transformations above is only to map the fitted results onto a scale comparable with GSFA, and is not necessary in practice.

In this comparison, we focus on predictive performance on cell-gene pairs whose expression levels are more likely to be perturbed. To do so, we first select entries in $\hat{\boldsymbol{\mu}}_{\text{GPerturb}}$ whose corresponding estimated posterior inclusion probability $\sigma(\hat{\eta}_p(\mathbf{C}_i)) > 0.95$, then report the scatter plot of the selected entries in perturbation effects $\hat{\boldsymbol{\mu}}_{\text{GPerturb}}$ verses the same set of selected entries in standardised residuals $\bar{\mathbf{X}}_{\text{GPerturb}}$ in Supplementary Fig 13. We also report the scatter plot of the same set of selected entries in $\hat{\boldsymbol{\mu}}_{\text{GSFA}}$ verses $\bar{\mathbf{X}}_{\text{GSFA}}$. We find that under this comparison framework in favour of GSFA, the post-processed GPerturb achieves similar performance to GSFA on the set of selected entries (Pearson correlation $r_{\text{GPerturb}} = 0.248$, $r_{\text{GSFA}} = 0.182$) in term of Pearson correlation between the transformed observations ($\bar{\mathbf{X}}_{\text{GSFA}}$, $\bar{\mathbf{X}}'_{\text{GPerturb}}$) and predictions ($\hat{\boldsymbol{\mu}}_{\text{GSFA}}$, $\hat{\boldsymbol{\mu}}'_{\text{GPerturb}}$), indicating that GPerturb is able to capture details of perturbation effects. The fitted verses observed scatter plot on test set is reported in Supplementary Fig 13. We remind the reader that this comparison is qualitative and not rigorous as the two models

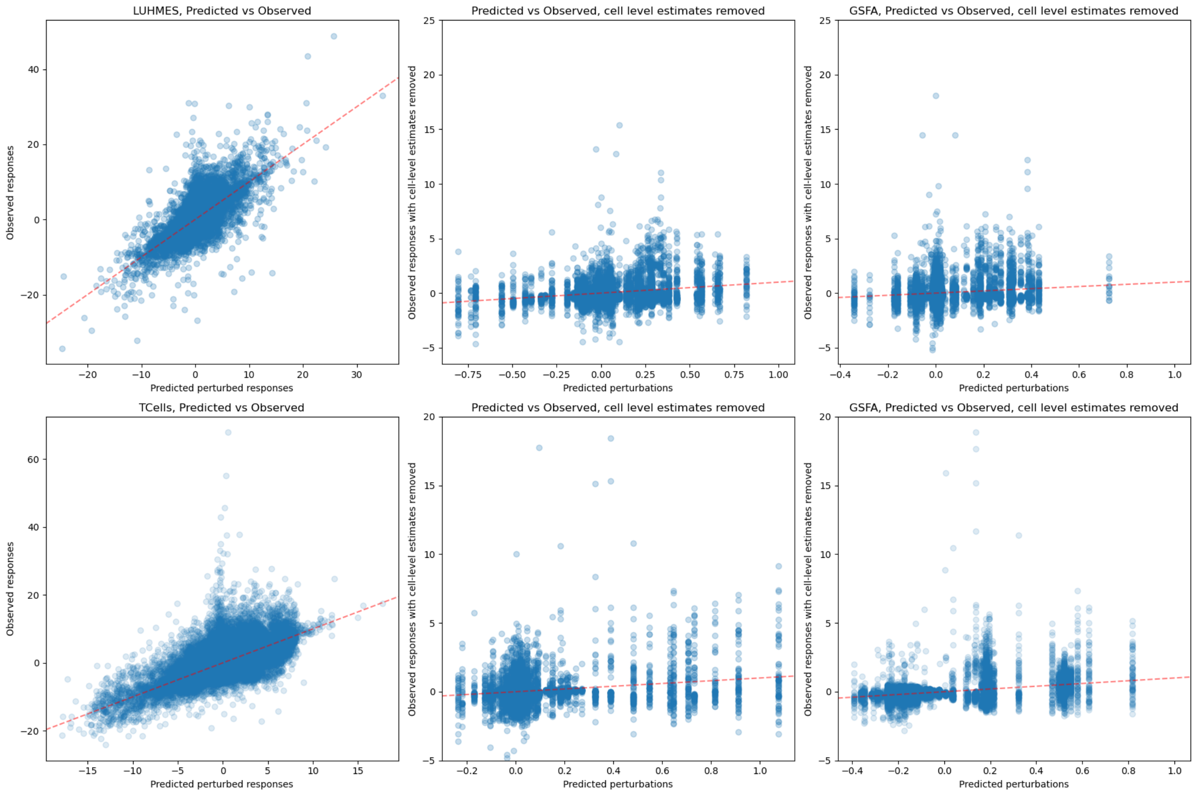

**Supplementary Figure 13: Top**: Fitted values of Gaussian GPerturb and GSFA on the LUHMES dataset. **Top left**: Predicted expression level given by Gaussian GPerturb vs observed expression level in the LUHMES test set. **Top middle**: Standardised perturbation effects $\hat{\boldsymbol{\mu}}'_{\text{GPerturb}}$ verses standardised residuals $\bar{\mathbf{X}}'_{\text{GPerturb}}$ of the selected "active" gene-sample pairs whose corresponding posterior inclusion probability $\sigma(\hat{\eta}_p(\mathbf{C}_i)) > 0.95$. **Top right**: Standardised perturbation effects $\hat{\boldsymbol{\eta}}_{\text{GSFA}}$ verses standardised residuals $\bar{\mathbf{X}}_{\text{GSFA}}$ of the same set of selected "active" gene-sample pairs. **Bottom**: Fitted values of Gaussian GPerturb and GSFA on the human T Cells dataset. Figures have the same interpretation as in the top row.

are estimated using different pre-processing steps and objectives.

We also report heat map of the estimated perturbation effects associated with each of the 14 unique perturbations $\{\mathbf{C}_i^*\}_{i=1}^{14}$ given by Gaussian GPerturb in Supplementary Fig 14. Here a perturbation-gene pair is considered "active" only if the corresponding estimated posterior inclusion probability $\sigma(\hat{\eta}_p(\mathbf{C}_i^*)) > 0.95$, and a gene $p$ is included only if at least one of the $\{\sigma(\hat{\eta}_p(\mathbf{C}_i^*))\}_{i=1}^{14}$ is greater than 0.95. The resulting heat map provides an intuitive visualization of perturbation effects on different genes. In addition to the genes associated with large estimated perturbation effects, we also report a similar heat map on a collection of marker genes studied in Zhou et al.[11] in Supplementary Fig 15. Comparing with Zhou et al.[11], we found that GPerturb identifies less active perturbation-gene pairs than GSFA (See also Supplementary Fig 19). To further investigate this difference, we select three marker genes (NES, CRABP2, HDAC2) from different pathway groups, and compare the observed expression level of these genes with the GPerturb prediction under three different perturbations (CHD2, PTEN, SETD5) (Supplementary Fig 16). For each perturbation-gene pair, we use a two-sample $t$-test to test if the mean of the expression levels under the perturbation is different from the baseline (i.e. mean expression level under the non-targeting perturbation). The $p$-values are then

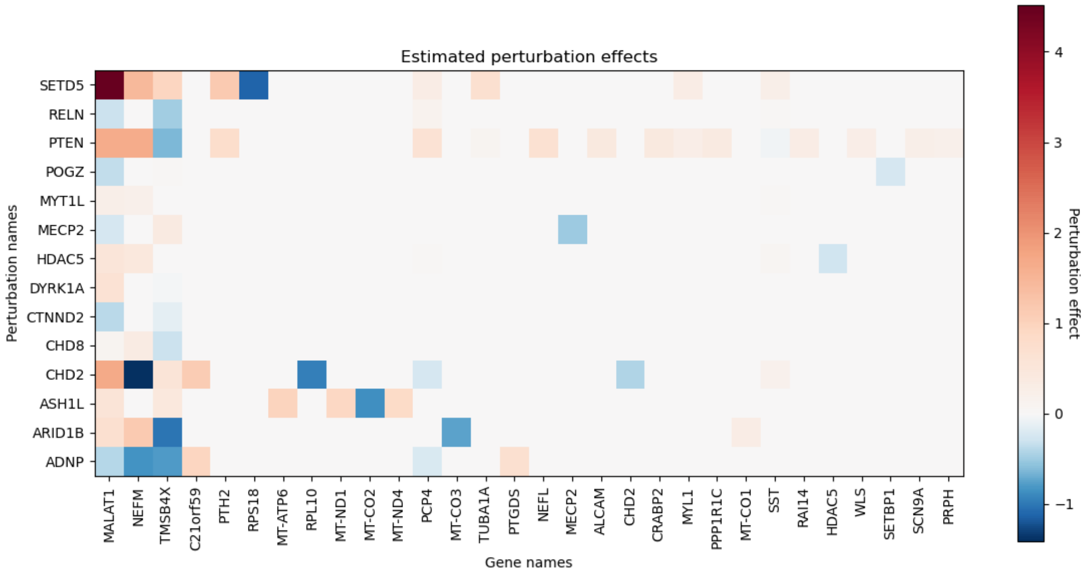

**Supplementary Figure 14:** Heat map of selected perturbation effects estimated from the LUHMES dataset. Each row corresponds to one of the unique perturbation $\{\mathbf{C}_i^*\}_{i=1}^{14}$. The perturbation effect of $\mathbf{C}_i^*$ on gene $p$ is included only if the associated posterior inclusion probability $\sigma(\hat{\eta}_p(\mathbf{C}_i)) > 0.95$.

corrected using Benjamini–Hochberg procedure (B-H)[1], and a gene-perturbation pair is considered active if the corrected $p$-value is less than 0.05. We then compare the subset of active perturbation-gene pairs identified by B-H with the ones identified by GPerturb and GSFA. From Supplementary Fig 16 we see GPerturb's predicted expression levels agree with the observed values, and the subset of pairs identified by GPerturb agrees better with the one identified by B-H, suggesting that GSFA may consist of more false positives.

To demonstrate scalability and versatility of our methods, we also apply Zero-inflated Poisson GPerturb to the raw counting data. We removed all zero columns from the raw counting matrix (i.e. genes with zero count across all samples), and apply Poisson GPerturb to the resulting data matrix consisting of $N = 8708$ samples and $P = 21688$ genes. The results are reported in Supplementary Fig 17, 18. Even though both the Gaussian and Poisson GPerturb fit the corresponding datasets reasonably well, the two approaches give slightly different estimates of the "active" perturbation-gene pairs, which is likely due to the data pre-processing and transformation steps. Hence we recommend users to analyse the dataset with different data pre-processings and compare their results. We also report in Supplementary Fig 19 histogram of the number of differentially expressed genes identified by Gaussian-GPerturb and existing methods similar to Supplementary Fig 5e in Zhou et al.[11].

#### S5.2 CD8+ T cell CROP-seq study

In this section, we apply Gaussian GPerturb to the primary human CD8+ T cells dataset (GSE119450) studied in Zhou et al.[11] in a similar fashion to the previous section. This study targets 20 genes associated with the T cell response, in both stimulated and unstimulated T cells. The processed dataset $\mathbf{X} \in \mathbb{R}^{N \times P}$ consists of $N = 24955$ samples and $P = 6000$ genes. For $i = 1, ..., N$, the perturbations $\mathbf{C}_i \in \{0, 1\}^L$ are one-hot vectors of length $L = 20$, which correspond to the 20 targeted genes in the study, and $\mathbf{K}_i \in \mathbb{R}^D$ is a real vector of length

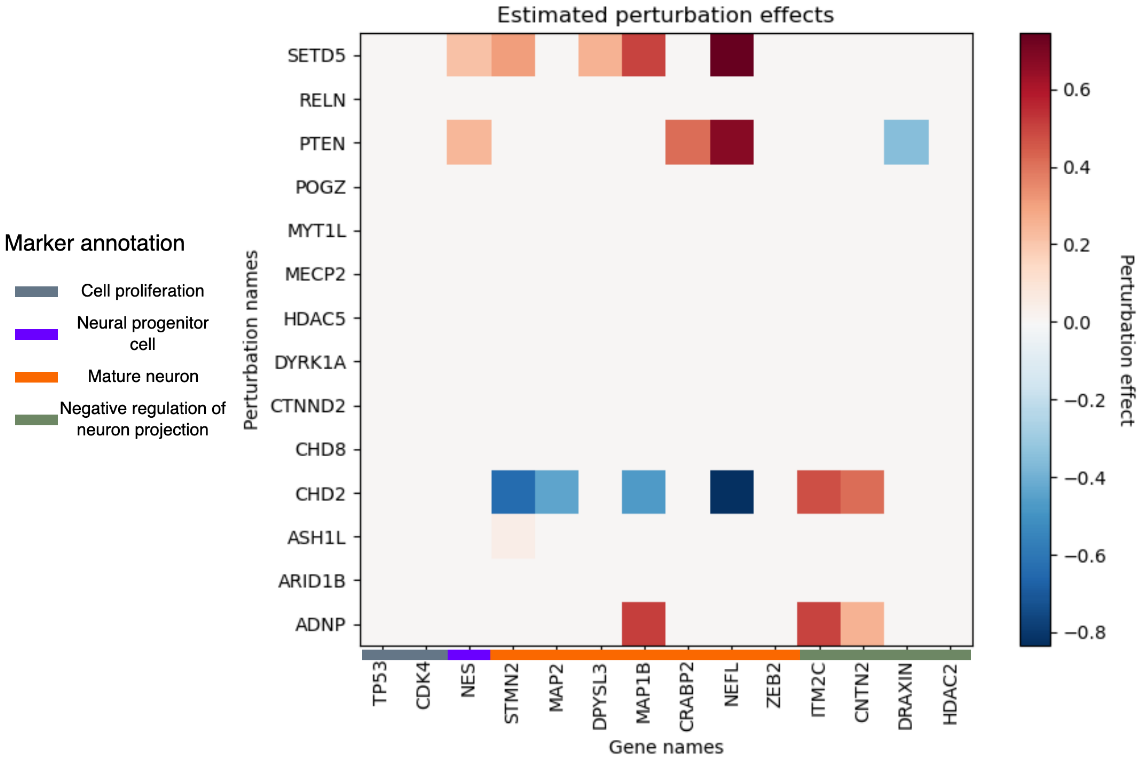

**Supplementary Figure 15:** Heat map of perturbation effects on the marker genes studied in Zhou et al.[11] estimated from the LUHMES dataset. Each row corresponds to one of the unique perturbation $\{\mathbf{C}_i^*\}_{i=1}^{14}$. The perturbation effect of $\mathbf{C}_i^*$ on gene $p$ is included only if the associated posterior inclusion probability $\sigma(\hat{\eta}_p(\mathbf{C}_i)) > 0.95$.

$D = 5$ (`lib_size`: number of total UMI counts, `n_features`: number of genes with non-zero UMI readings, `mt_percent`: percentage of mitochondrial gene expression, `donor`: T Cell donor ID and `stimulated`: whether or not the T Cell is stimulated). In Zhou et al.[11], the authors hypothesised that perturbation effects are different in stimulated and unstimulated cells, and used a modified GSFA to capture such difference.

In this example, we use a modified Gaussian GPerturb model to accommodate potentially different perturbation effects for stimulated and unsimulated T Cells in a similar fashion. Our objective is to capture potentially different perturbation effects associated with different cell groups under the same perturbation. For simplicity, we demonstrate the modified model with two cell groups, which corresponds to the stimulated and unstimulated T cells. We assume $\mathbf{K}_i^{(1)} \in \{0, 1\}$, the first entry of cell information, encodes the cell group indicator. Extension to larger number of cell groups or other cell conditions is straightforward.

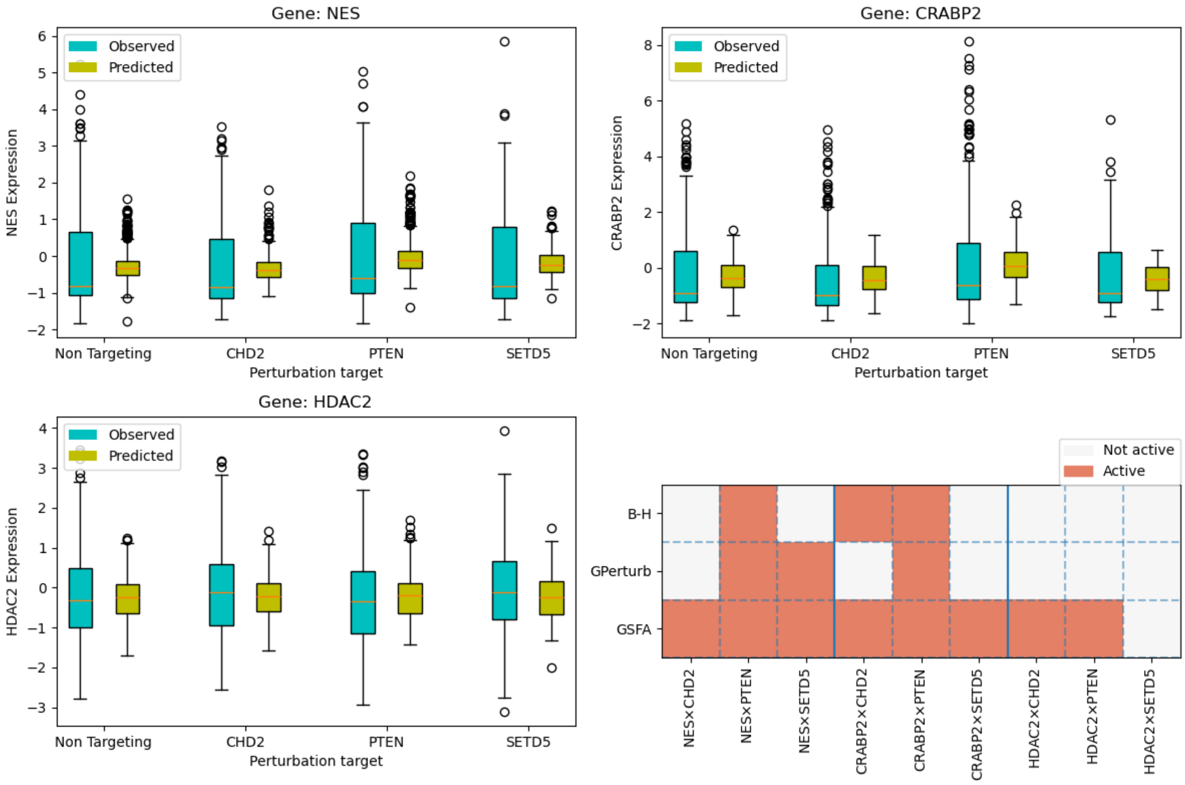

**Supplementary Figure 16:** Comparing the expression levels under different perturbations. **Top left**: Boxplots of the observed and GPerturb predicted expression levels of gene NES in test set. **Top right**: Boxplots of the observed and GPerturb predicted expression levels of gene CRABP2 in test set. **Bottom left**: Boxplots of the observed and GPerturb predicted expression levels of gene HDAC2 in test set. **Bottom right**: Subset of gene-perturbation pairs selected by Benjamini–Hochberg, GPerturb and GSFA respectively.

We start from the modified generative process

$$m_p : \mathbb{R}^D \to \mathbb{R}; \quad \lambda_p \in \mathbb{R}; \tag{6}$$

$$\mu_p^{(0)} \sim \mathcal{GP}(g_\mu, k_{\nu_\mu}); \quad \mu_p^{(1)} \sim \mathcal{GP}(g_\mu, k_{\nu_\mu}); \quad \gamma_p^{(0)} \sim \mathcal{GP}(g_\gamma, k_{\nu_\gamma}); \quad \gamma_p^{(1)} \sim \mathcal{GP}(g_\gamma, k_{\nu_\gamma}); \tag{7}$$

$$\eta_p^{(0)} \sim \mathcal{GP}(g_\eta, k_{\nu_\eta}); \quad z_{ip}^{(0)} \sim \text{Bernoulli}\left(\sigma(\eta_p^{(0)}(\mathbf{C}_i))\right); \tag{8}$$

$$\eta_p^{(1)} \sim \mathcal{GP}(g_\eta, k_{\nu_\eta}); \quad z_{ip}^{(1)} \sim \text{Bernoulli}\left(\sigma(\eta_p^{(1)}(\mathbf{C}_i))\right); \tag{9}$$

$$X_{ip} \sim \mathcal{N}\left(m_p(\mathbf{K}_i) + z_{ip}^{(\mathbf{K}_i^{(1)})}\mu_p^{(\mathbf{K}_i^{(1)})}(\mathbf{C}_i),\ \log(\exp(\lambda_p + z_{ip}^{(\mathbf{K}_i^{(1)})}\gamma_p^{(\mathbf{K}_i^{(1)})}(\mathbf{C}_i)) + 1)\right), \tag{10}$$

In other words, the generative process assumes that two cell groups share the common basal mean function $\mu_p$ (whose output also depends on the cell group indicator), but are associated with different perturbation effects $\{\mu_p^{(0)}, \gamma_p^{(0)}, \eta_p^{(0)}\}$ and $\{\mu_p^{(1)}, \gamma_p^{(1)}, \eta_p^{(1)}\}$.

The variational family is then modified accordingly: We replace $f_{\boldsymbol{\xi}} : \mathbb{R}^L \to \mathbb{R}^{6P}$ in Eqn (13) by $g_{\boldsymbol{\xi}} : \mathbb{R}^{L+1} \to \mathbb{R}^{6P}$, which takes the augmented perturbation vector $\{\mathbf{C}_i, \mathbf{K}_i^{(1)}\} \in \mathbb{R}^{L+1}$ as its input. The new function $g_{\boldsymbol{\xi}}$ alongside with $f_{\boldsymbol{\phi}}$ and $\boldsymbol{\lambda}$ are estimated by minimizing the ELBO given in Eqn (10) in a similar fashion to the original Gaussian GPerturb. The Poisson GPerturb is modified in a similar fashion.

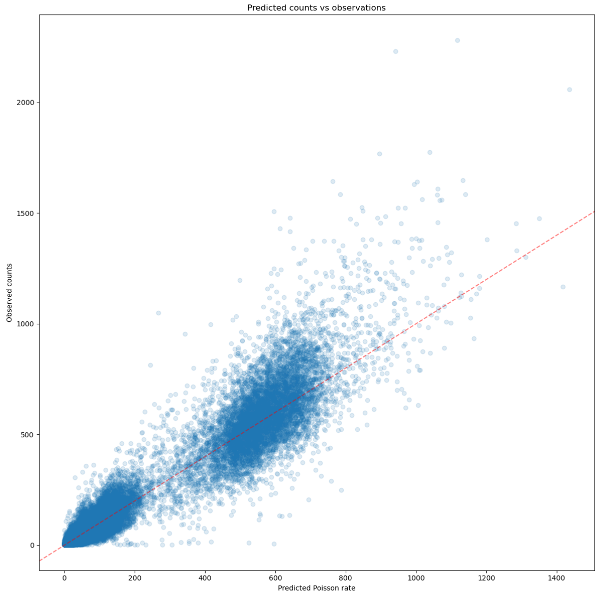

**Supplementary Figure 17:** Non-zero observed counts for each cell-gene pair vs corresponding estimated Poisson rate for each cell-gene pair given by Poisson GPerturb

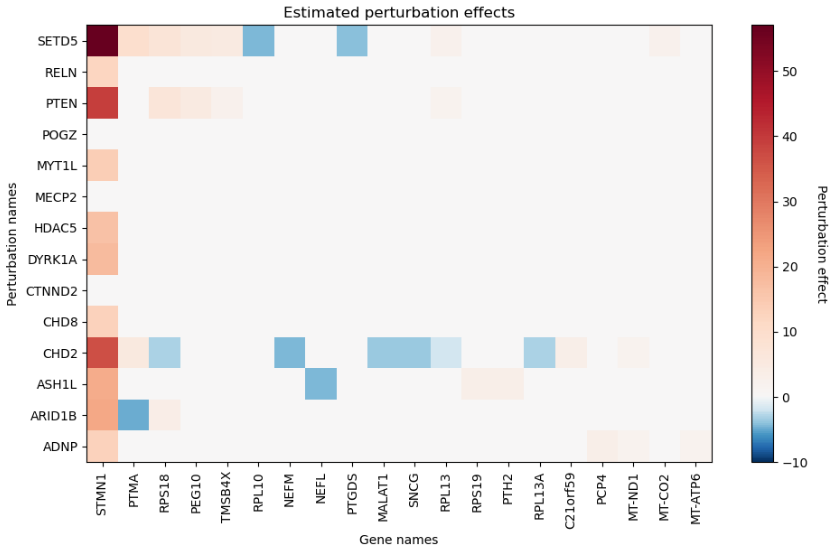

**Supplementary Figure 18:** Heat map of the estimated perturbation effects given by Poisson GPerturb. Similar to Supplementary Fig 14, the perturbation effects of $\mathbf{C}_i^*$ on gene $p$ is included only if the associated posterior inclusion probability $\sigma(\hat{\eta}_p(\mathbf{C}_i)) > 0.95$.

Similar to the previous example, we randomly select 20% of the dataset as the test set, and use the rest to train GPerturb. We report the fitted results in Supplementary Fig 13. Similar to the previous section, we compare the predictive performance between GPerturb and GSFA on the cell-gene pairs in test set whose estimated posterior inclusion probability is greater than 0.95. Here we found post-processed Gaussian GPerturb and GSFA achieve comparable prediction performance ($r_{\text{GPerturb}} = 0.271$, $r_{\text{GSFA}} = 0.335$).

The heat maps of estimated perturbation effects associated with unique perturbations are reported in Supplementary Fig 20. Here we only include the top 30 genes sorted by the magnitude

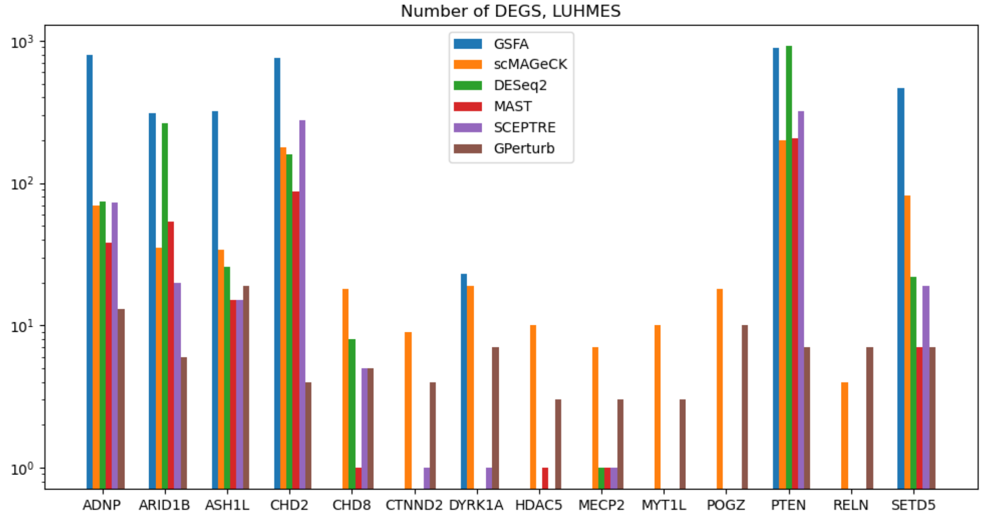

**Supplementary Figure 19:** Histogram of the number of differentially expressed genes identified by different methods, LUHMES dataset. This figure is modified from Supplementary Fig 5e in [11].

of overall perturbation effects in the heat map for sake of visual. We also report the heat map of perturbation effects on the collection of marker genes studied in Zhou et al. [11] in Supplementary Fig 21. In Supplementary Fig 22 and 23, we compare the expression level of three marker genes (IL7R, TNFRSF18, MKI67) under three perturbations (LCP2, CBLB, TCEB2) in a similar fashionn to the previous section. Again we see that GPerturb agrees better with the B-H corrected two-sample $t$-tests, and GSFA tends to pick up more perturbation-gene pairs whose mean expression levels are not significantly different from the baseline. We also apply the Zero-inflated Poisson GPerturb model to the raw counting data consists of $N = 8708$ samples and $P = 22400$ genes. The results are reported in Supplementary Fig 24, 25.

We also applied Zero-inflated Poisson GPerturb to the raw counting data of the human T Cells dataset in Section S5.2. Results are reported in Supplementary Fig 24 and 25. We also report in Supplementary Fig 26, a histogram of the number of differentially expressed genes identified by Gaussian GPerturb and existing methods similar to Supplementary Fig 5e in Zhou et al. [11].

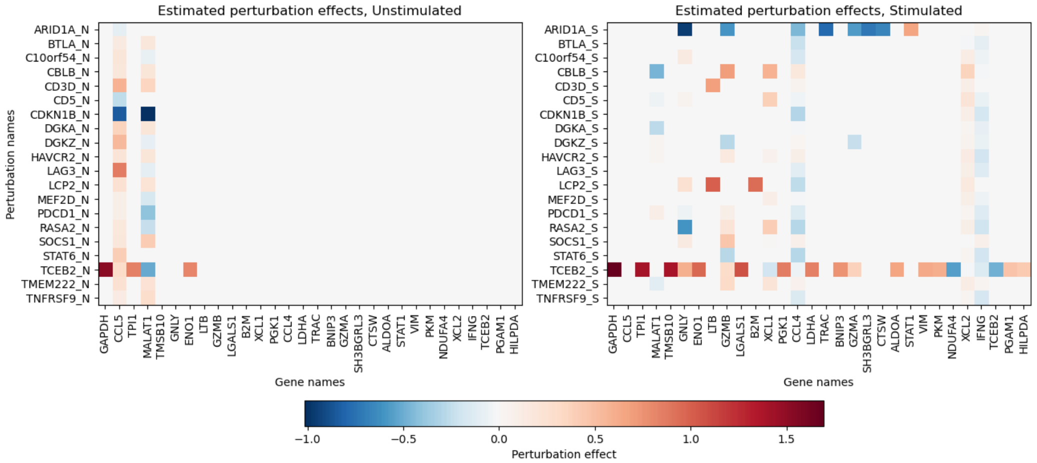

**Supplementary Figure 20:** Heat map of perturbation effects estimated from the human T Cells dataset. **Left**: Estimated perturbation effects on unstimulated T cells. **Right**: Estimated perturbation effects on stimulated T cells. The perturbation effect of $\mathbf{C}_i^*$ on gene $p$ is included only if the associated posterior inclusion probability $\sigma(\hat{\eta}_p(\mathbf{C}_i)) > 0.95$. We only include the top 30 genes sorted by the magnitude of overall perturbation effects in the heat map for sake of visual.

[7] Srivatsan, S. R., McFaline-Figueroa, J. L., Ramani, V., Saunders, L., Cao, J., Packer, J.,

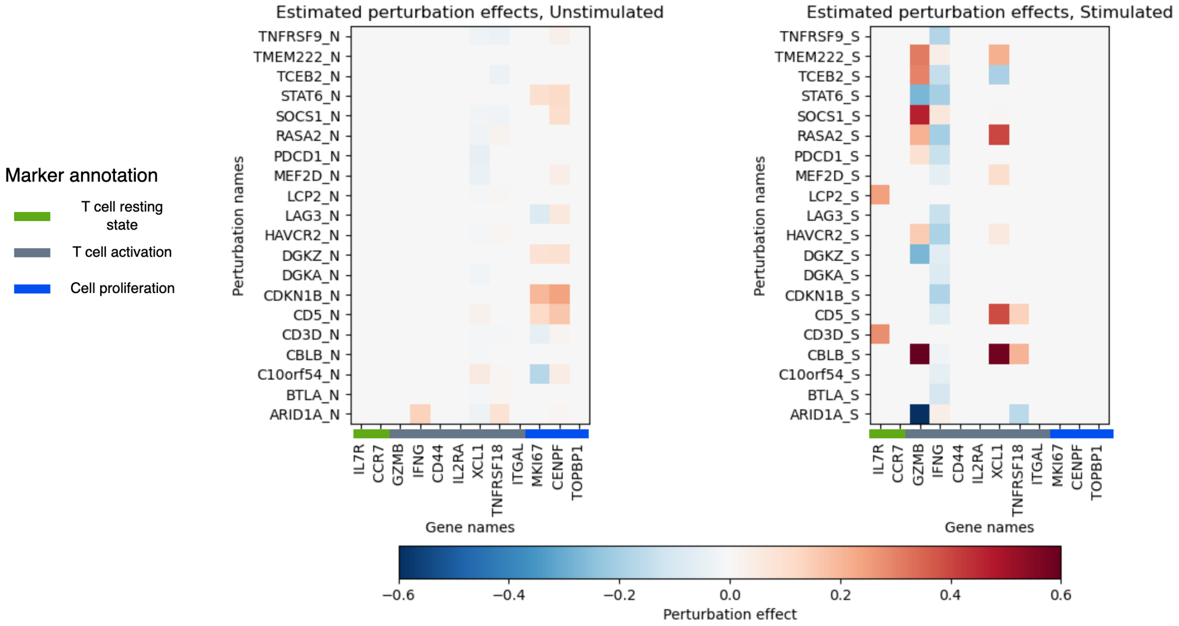

**Supplementary Figure 21:** Heat map of perturbation effects on the marker genes studied in [11] estimated from the human T cell dataset. **Left**: Estimated perturbation effects on unstimulated T cells. **Right**: Estimated perturbation effects on stimulated T cells. Each row corresponds to one of the unique perturbation $\{\mathbf{C}_i^*\}_{i=1}^{14}$. The perturbation effect of $\mathbf{C}_i^*$ on gene $p$ is included only if the associated posterior inclusion probability $\sigma(\hat{\eta}_p(\mathbf{C}_i)) > 0.95$.

Pliner, H. A., Jackson, D. L., Daza, R. M., Christiansen, L., et al. (2020). Massively multiplex chemical transcriptomics at single-cell resolution. *Science*, 367(6473):45–51.

[8] Stephens, M. (2017). False discovery rates: a new deal. *Biostatistics*, 18(2):275–294.

[9] Townes, F. W., Hicks, S. C., Aryee, M. J., and Irizarry, R. A. (2019). Feature selection and dimension reduction for single-cell rna-seq based on a multinomial model. *Genome biology*, 20:1–16.

[10] Yao, D., Binan, L., Bezney, J., Simonton, B., Freedman, J., Frangieh, C. J., Dey, K., Geiger-Schuller, K., Eraslan, B., Gusev, A., et al. (2023). Scalable genetic screening for regulatory circuits using compressed perturb-seq. *Nature Biotechnology*, pages 1–14.

[11] Zhou, Y., Luo, K., Liang, L., Chen, M., and He, X. (2023). A new bayesian factor analysis method improves detection of genes and biological processes affected by perturbations in single-cell crispr screening. *Nature Methods*, 20(11):1693–1703.

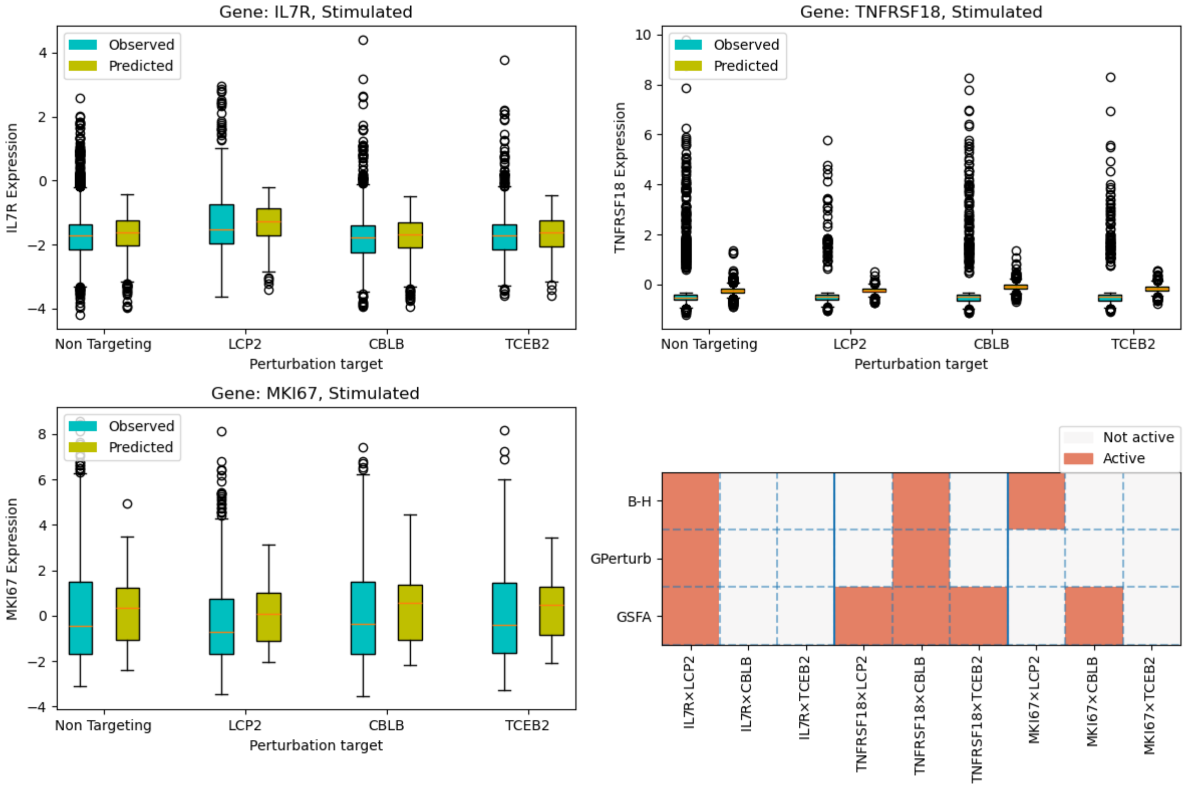

**Supplementary Figure 22:** Comparing the expression levels in stimulated T cells under different perturbations. **Top left**: Boxplots of the observed and GPerturb predicted expression levels of gene IL7R in test set. **Top right**: Boxplots of the observed and GPerturb predicted expression levels of gene TNFTSF18 in test set. **Bottom left**: Boxplots of the observed and GPerturb predicted expression levels of gene MKI67 in test set. **Bottom right**: Subset of gene-perturbation pairs selected by Benjamini–Hochberg, GPerturb and GSFA respectively.

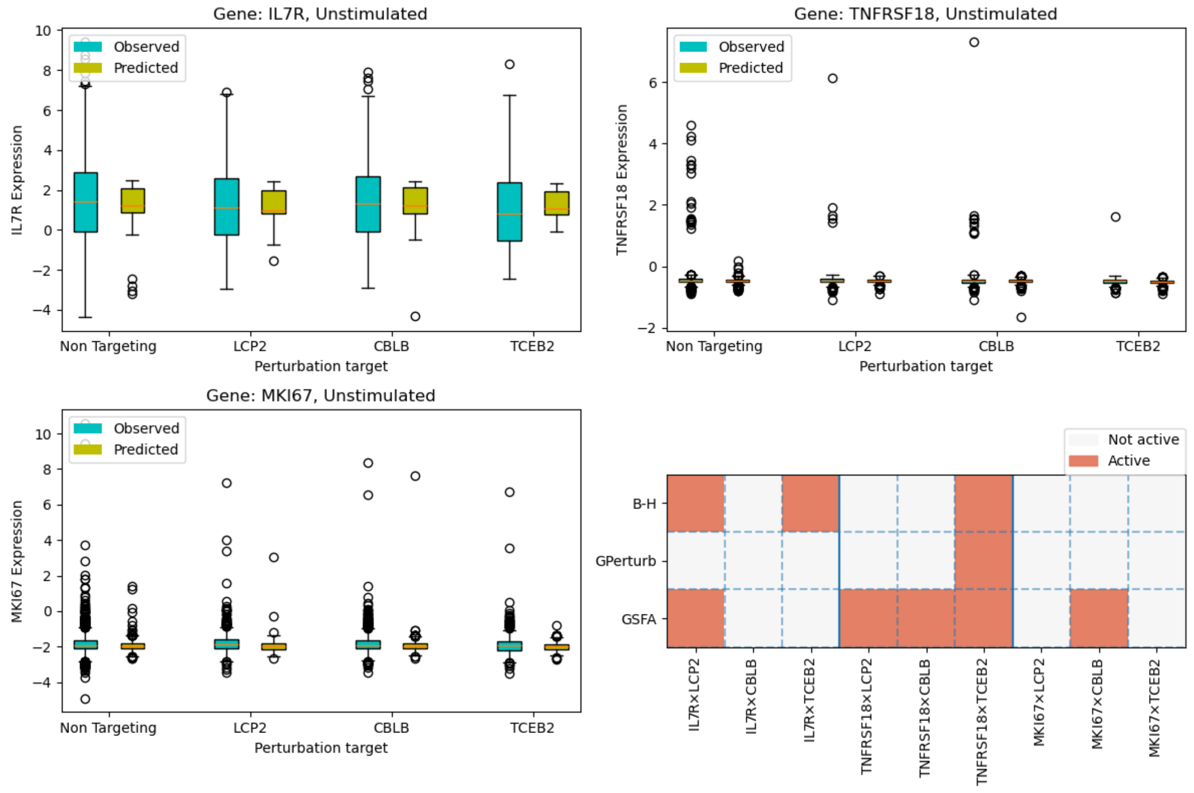

**Supplementary Figure 23:** Comparing the expression levels in unstimulated T cells under different perturbations. **Top left**: Boxplots of the observed and GPerturb predicted expression levels of gene IL7R in test set. **Top right**: Boxplots of the observed and GPerturb predicted expression levels of gene TNFTSF18 in test set. **Bottom left**: Boxplots of the observed and GPerturb predicted expression levels of gene MKI67 in test set. **Bottom right**: Subset of gene-perturbation pairs selected by Benjamini–Hochberg, GPerturb and GSFA respectively.

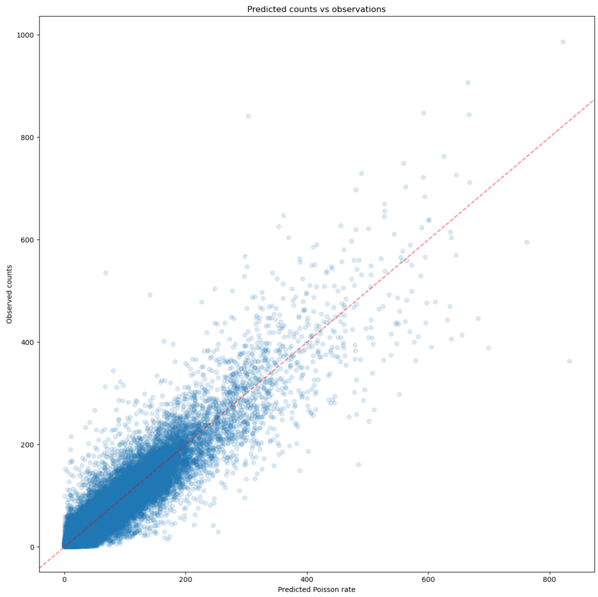

**Supplementary Figure 24:** Human T Cells dataset, Non-zero observed counts for each cell-gene pair vs corresponding estimated Poisson rate for each cell-gene pair given by Poisson GPerturb

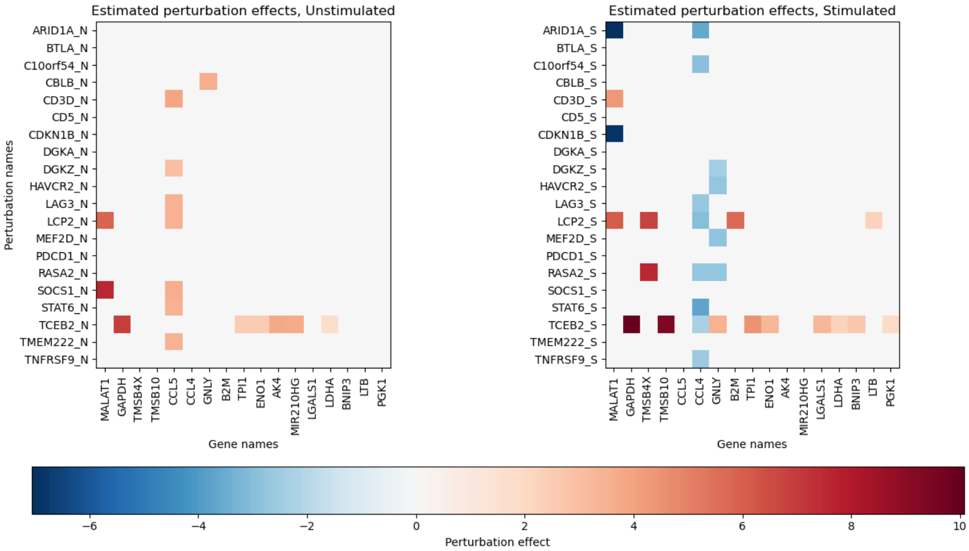

**Supplementary Figure 25:** Heat map of the estimated perturbation effects given by Poisson GPerturb. Similar to Supplementary Fig 20, each row corresponds to the perturbation effects of a unique perturbation on simulated (_S) or unstimulated (_N) T Cells. The perturbation effects on gene $p$ is included only if the associated posterior inclusion probability $\sigma(\hat{\eta}_p(\mathbf{C}_i^*)) > 0.95$.

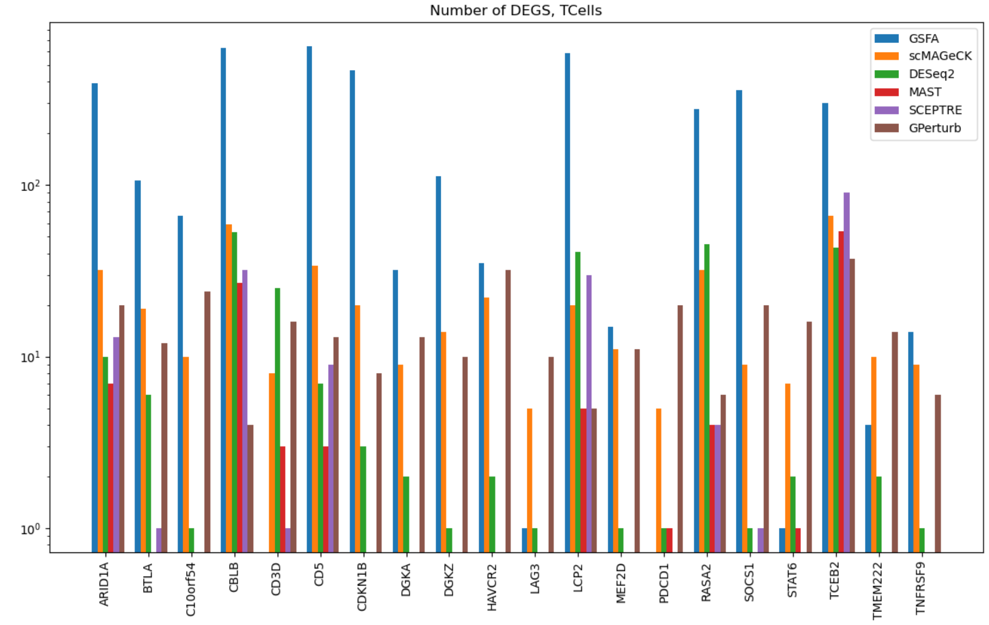

**Supplementary Figure 26:** Histogram of the number of differentially expressed genes identified by different methods, human T cell dataset. This figure is modified from Supplementary Fig 5e in Zhou et al.[11].
